# Robust geometric drift correction and identity reconstruction for biological 3D time-lapse point clouds

**DOI:** 10.64898/2026.10.17.759823

**Authors:** Fabrizio Musacchio, Martin Fuhrmann

## Abstract

Structural time-lapse microscopy experiments aim to follow cells or subcellular structures over time, typically by detecting objects in each frame and linking them into trajectories. In practice, this process yields sequences of spatial coordinates derived from segmentation, while identity information across time is often absent or unreliable. In addition, the observed coordinates may be affected by systematic geometric drift. As a result, two coupled inference problems arise: geometric drift must be corrected, and object identities must be reconstructed from unlabeled point clouds.

Here, we formulate both drift correction and identity reconstruction within a unified point-cloud framework and evaluate them jointly using controlled synthetic benchmarks and biological datasets. For drift correction, we compare rigid, local, and nonparametric models across spatially varying and non-rigid drift regimes. For identity reconstruction, we analyze a family of Hungarian assignment methods with constant-velocity refinement and different gating strategies, including innovation-based normalization.

Across benchmarks, we find that increased model flexibility does not necessarily improve drift correction. A simple rigid stepwise model provides the most robust performance across a wide range of conditions, with a local community model as a competitive alternative under structured spatial heterogeneity. On the association side, innovation-based gating emerges as the key determinant of performance. Hungarian constant-velocity assignment with normalized innovation gating consistently outperforms coordinate-space variants, particularly under heterogeneous motion scales, birth and death processes, and density-induced ambiguity.

Application to biological datasets confirms that the combined pipeline improves trajectory consistency and restores interpretable motion statistics, while highlighting the role of proxy references when ground truth is unavailable. Together, these results establish a practical and robust workflow for post hoc trajectory analysis in point-cloud-based time-lapse data, and clarify the methodological trade-offs between model expressivity and robustness in both drift correction and identity reconstruction.

**Author Summary:** Time-lapse microscopy is widely used to study how cells and subcellular structures move and change over time. A common analysis goal is to track these objects by detecting them in each frame and linking them into trajectories. In practice, however, this process often produces only their spatial coordinates, while consistent identity information across time is missing or unreliable. In addition, the entire field of view may shift or rotate during imaging, a problem known as drift.

We developed and systematically evaluated a computational framework that addresses both challenges using coordinate-based data alone. By combining controlled synthetic benchmarks with real biological datasets, we show that simple and robust approaches often outperform more complex ones. In particular, a rigid drift correction model remains effective even when drift varies across space, and trajectory reconstruction improves substantially when candidate matches are evaluated relative to their uncertainty rather than by fixed distance thresholds.

Our results provide practical guidance for analyzing time-lapse data when only coordinates are available. More broadly, they show that accounting for uncertainty and prioritizing robustness over model complexity leads to more reliable trajectory reconstruction in noisy and heterogeneous biological systems.

## Introduction

Time-lapse microscopy is widely used to quantify structural dynamics in living tissue, including the appearance, displacement, migration, and disappearance of cellular and subcellular objects. In many experimental settings, volumetric image stacks are recorded repeatedly over time, yielding a sequence of three-dimensional spatial volumes sampled at discrete time points. Such recordings are often referred to as 3D time-lapse or 3D+t acquisitions, emphasizing the three spatial dimensions and the additional temporal dimension. A primary goal in these experiments is frequently to quantify how individual cells or other segmented structures evolve over time, for example by counting objects, measuring their spatial distribution, and reconstructing their trajectories across successive volumes.

To enable such quantitative analysis, each three-dimensional volume is typically segmented to identify objects of interest. In practice, segmentation is performed using a variety of established tools and methods. Commercial platforms such as Imaris (Oxford Instruments, Bitplane, 2024) provide workflows based on intensity thresholding, region growing, and model-based fitting, while open-source and deep learning–based approaches such as Cellpose (Stringer et al., 2021) employ convolutional neural networks trained to delineate cellular boundaries. Depending on image quality and experimental design, segmentation may rely on classical intensity-based thresholding, watershed methods, or modern convolutional neural network architectures. Irrespective of the specific approach, the output of segmentation is usually a set of spatially extended objects for each time point.

For many downstream analyses, the full voxel representation of each segmented object is not required. Instead, it is common to extract compact geometric descriptors such as centroid coordinates, volumes, or simple shape features. In particular, when the primary objective is to count cells and track them over time, the temporal sequence of centroid coordinates is often sufficient to approximate cell trajectories. From these reconstructed trajectories, one can compute displacement statistics, motility measures, persistence, and event rates such as object appearances and disappearances, which correspond to biological cell births and deaths. Thus, the practical workflow frequently reduces a sequence of volumetric images to a time-indexed three-dimensional point cloud representation.

However, this reduction has methodological consequences. Classical image registration algorithms operate directly on voxel intensities and can exploit spatially distributed texture and structural information to estimate transformations between volumes. Once the data have been reduced to point clouds of object centroids, much of this spatial context is no longer available. Any residual motion artifacts or imperfect alignment that remain after initial rigid or semi-rigid registration of the image stacks are therefore inherited by the centroid coordinates. As a result, the subsequent geometric analysis becomes vulnerable to systematic misalignment and drift that are not easily corrected at the image level without risking distortion of genuine structural changes.

A particularly challenging instance of this problem arises in in vivo multiphoton microscopy. Two-photon and three-photon imaging provide optical sectioning and deep tissue access, but they are susceptible to motion from breathing, heartbeat, and slow sample drift (Denk, Strickler and Webb, 1990; Helmchen and Denk, 2005; Wang et al., 2018; Fuhrmann, Nebenlin, Musacchio et al., 2025). In functional calcium imaging, this challenge is often mitigated by image-based motion correction methods that register each frame or each patch of the field of view to a reference template. Representative approaches include subpixel registration methods based on Fourier phase correlation (Guizar-Sicairos, Thurman and Fienup, 2008) as well as template matching and piecewise rigid models, both in dedicated motion correction algorithms and in broader analysis suites (Pnevmatikakis and Giovannucci, 2017; Giovannucci et al., 2019; Kaifosh et al., 2014). These methods are extremely successful in two-dimensional time-lapse (2D+t) and quasi-2D+t imaging regimes. Here, quasi-2D+t refers to volumetric acquisitions with limited axial extent or strongly anisotropic sampling, where motion correction is often approximated as predominantly planar. Nonetheless, structural time-lapse imaging introduces a different constraint. Here, the objects of interest can move, appear, disappear, or change morphology over time, and these dynamics are themselves the biological signal. A registration model that is too flexible, such as a high degree affine or deformable warp estimated via optical flow or local gradient-based methods (Lucas and Kanade, 1981; Farnebäck, 2003; Greenberg and Kerr, 2009), risks absorbing real biological motion into the estimated transformation field, thereby conflating correction with measurement. This is conceptually problematic: In structural time-lapse, one cannot assume that temporal changes in object geometry are nuisance variation that should be removed by an unconstrained transformation. Instead, the transformation class must be carefully chosen so that systematic acquisition induced misalignment is corrected while genuine structural changes remain observable.

The multiphoton setting also highlights an additional difficulty that is not fully addressed by standard two-dimensional motion correction. Volumetric imaging is intrinsically anisotropic, since axial resolution is typically worse than lateral resolution, the signal to noise ratio can vary strongly with depth, and scanning is performed sequentially in time rather than instantaneously across the volume. These factors complicate volumetric rigid registration that includes full 3D rotations, especially when the imaged content itself changes over time. While there are specialized hardware and acquisition strategies that enable real-time 3D movement correction with submicrometer precision, they rely on dedicated instrumentation, closed-loop scanning, or additional assumptions that are not available in many laboratories (Griffiths et al., 2020). Conversely, advanced deformable registration frameworks developed for medical imaging, such as ANTs and elastix, are powerful and broadly applicable, but they are designed to estimate high-dimensional transformations in regimes where the underlying anatomy is assumed to be comparable between images and where the goal is often to maximize alignment even under genuine deformation (Avants et al., 2011; Klein et al., 2010). For 3D time-lapse microscopy of dynamic cellular objects, such flexibility can be undesirable when it risks erasing or distorting the biological dynamics that motivate the experiment in the first place.

These considerations motivate a different perspective. Rather than treating motion correction as an image registration problem, one can formulate the task at the level of the extracted point clouds. Each time point provides a set of observed object positions in ℝ^3^. The observed positions are affected by (i) systematic geometric drift that depends on time and potentially on spatial location within the volume, (ii) object specific motion that may be heterogeneous across objects and across time, (iii) measurement noise, and (iv) changes in object population due to births and deaths. The computational goal is then to reconstruct consistent object trajectories and to correct systematic drift, using only geometric information. This setting is common in practice because many segmentation and visualization tools export centroid coordinates and, depending on the software and settings, the exported data may not contain stable track identities. Tracking tools exist and are widely used, for example in the Fiji ecosystem (Tinevez et al., 2017; Ershov et al., 2022), but robust tracking remains difficult under dense object populations, near collisions, heterogeneous motion statistics, and missing or unreliable identity labels. The combination of drift correction and identity reconstruction is therefore not an optional refinement. It is a requirement for quantitative structural time-lapse analysis in many realistic regimes.

Drift estimation from point trajectories presupposes trajectory continuity. If one estimates a drift field from the temporal evolution of points that are assumed to represent the same object across time, then identity errors such as ID switches and track fragmentation will bias the drift estimate. Drift correction in the point cloud setting is therefore sensitive to the consistency of object identities. At the same time, identity reconstruction itself depends on the geometry of the observations. Drift changes inter-point distances, apparent velocities, and relative displacements, thereby perturbing the cost structure that drives assignment decisions. Association algorithms that rely on spatial proximity or predicted motion are thus sensitive to uncorrected drift. These two dependencies imply a coupled inference problem. Accurate drift estimation requires consistent trajectories, while reliable trajectory reconstruction benefits from geometrically corrected observations. A rigorous methodological treatment must therefore address drift correction and identity reconstruction jointly, making explicit their mutual dependencies and their respective failure modes under realistic data regimes.

In this work, we study geometric drift correction and identity reconstruction in three-dimensional time-lapse point clouds. Although the problem is motivated by multiphoton structural imaging, the formulation is not restricted to a specific imaging modality. Instead, we consider the general setting of temporally sampled 3D point sets subject to systematic geometric drift, object-specific motion, and dynamic population changes. On the drift side, we consider a hierarchy of drift models that range from globally rigid motion to spatially varying drift fields. We benchmark rigid models against local models that allow different regions of the volume to drift differently, and against smooth function approximators such as radial basis function models, including spatiotemporal variants that explicitly regularize drift in both space and time. These models estimate drift directly from point cloud geometry and do not require image registration. On the identity side, we cast identity reconstruction without track labels as a frame-to-frame assignment problem. We compare Euclidean gating strategies, which can be effective when motion statistics are homogeneous, to uncertainty aware gating strategies based on innovations in a state space estimation view, where gating is performed in a normalized innovation space. We evaluate robustness in regimes that are known to break naive approaches, including heteroskedastic motion where displacement statistics vary across objects or across time, mid-sequence births and deaths, and near collision scenarios that produce intrinsic assignment ambiguity.

Taken together, the study proceeds along two closely related lines of analysis. On the geometric side, we examine whether systematic drift can be reconstructed reliably from point clouds using purely geometric models, and we characterize the regimes in which globally rigid transformations are sufficient versus those that require spatially varying drift fields. We further quantify how inaccuracies in drift correction propagate into downstream trajectory metrics, including RMS-type displacement errors and identity consistency measures, thereby linking geometric correction quality to tracking reliability. On the association side, we analyze identity reconstruction in the absence of explicit track labels in three-dimensional settings. We evaluate assignment strategies based on Euclidean gating under homoskedastic motion and compare them to uncertainty-aware gating strategies formulated in innovation space. In particular, we examine how these approaches behave under heteroskedastic motion, where displacement statistics vary across objects or across time, and where gating must remain robust to unknown or spatially varying motion scales. Finally, we investigate dynamic population scenarios with births, deaths, and near-collision ambiguity, and assess under which modeling assumptions stable trajectory reconstruction remains possible. Together, these analyses provide a mathematically explicit characterization of drift correction and identity reconstruction in 3D time-lapse point clouds, treating systematic geometric bias and missing identity information as coupled components of a single inference problem.

## Results

We first evaluate the candidate drift-correction models on synthetic benchmarks in which the ground-truth drift field is known, beginning with baseline spatially varying regimes and then moving to more structured non-rigid stress cases. We then turn to point-cloud association (PCA), reconstructing identities from unordered point clouds and comparing assignment strategies under progressively harder ambiguity, from controlled no-drift displacement sweeps to birth/death, mixed-motility, residual-drift, and density stress tests. Finally, we ask whether the synthetic rankings transfer to real biological data by applying the full PCA → DriftCorr workflow to a volumetric carcinoma-cell example with imposed rigid drift and to a planar DAPI time-lapse sequence with intrinsic experimental drift.

### Synthetic drift-correction benchmarks on baseline spatially varying drift regimes

The first synthetic benchmark block evaluates drift correction under baseline regimes that already depart from the assumption of a single globally rigid transformation. In these regimes, the synthetic generator produces spatially varying drift fields whose structure ranges from smoothly varying spatial fields to region-wise drift patterns with either gradual or abrupt spatial transitions. These baseline regimes therefore already represent non-uniform drift and serve as the first comparison setting in which globally rigid correction models are expected to degrade relative to spatially flexible approaches. The primary goals of this section are therefore to establish the initial method ranking under spatially varying drift, to examine whether the space-time RBF model admits a stable and interpretable parameter region in such regimes, and to test whether drift estimation can be improved by anchoring the fit to stationary trajectories rather than to the full mixed population.

#### Baseline suite benchmark across spatially varying drift regimes

We selected seven candidate correction models for this benchmark: the rigid stepwise model (Eq. (5), Eq. (11)), the local community model (Eq. (21), Eq. (22)), the per-timepoint RBF model (Eq. (25), Eq. (23), Eq. (36)), the joint space-time RBF model (`st_rbf`; Eq. (41), Eq. (43), Eq. (44)), and three mixture-of-experts variants of the space-time RBF model (Eq. (45), Eq. (46), Eq. (47)). These methods represent a range of modeling assumptions and spatial flexibility, from globally rigid to locally adaptive to fully nonparametric approaches. The benchmark therefore probes how well different correction strategies recover latent trajectories when drift varies across the field of view. In all baseline-suite runs, model hyperparameters were fixed as follows: rigid used trimmed pairwise fitting with inlier fraction *q* = 0.7, minimum pair count *n*_min_ = 15, and 30 inlier re-estimation iterations; local used a *k*-NN graph with *k* = 12, graph length scale *ℓ*_graph_ = 80 *µ*m, drift-field blending scale *ℓ*_field_ = 120 *µ*m, minimum edge weight *w*_min_ = 0.25, and at most 50 candidate edges per node; per-timepoint RBF used *M* = 45 centers, *λ* = 3 *×* 10^−4^, kernel width *σ* = 180 *µ*m, temporal coupling *γ* = 1.0, and trimmed fitting with *q* = 0.7; ST-RBF used *M* = 70 centers, *λ* = 8.5 *×* 10^−4^, (*ℓ*_*s*_, *ℓ*_*t*_) = (4.0, 0.4), robust keep fraction *q* = 0.83, 2 robust iterations, and 3 refinement iterations; ST-RBF-MoE reused the same ST-RBF base settings and was evaluated with *k* ∈ { 2, 3, 5} experts and blending width *σ*_blend_ = 140 *µ*m. These hyperparameters were fixed a priori from preliminary pilot runs and kept unchanged across all baseline-suite regimes to ensure that comparisons reflect method behavior rather than differences in data exposure or tuning. In subsequent analyses, we then explicitly revisited this choice in two ways: first, by performing targeted hyperparameter and robustness sweeps for the more flexible RBF-based models (Figure 3), and second, by reporting supplementary fairness sweeps for the rigid and local baselines (Supplementary Figure 12). These follow-up analyses show that moderate retuning can shift quantitative errors somewhat, but does not alter the qualitative ranking of methods established by the baseline suite.

To evaluate the general behavior of these candidate correction models once the assumption of globally uniform drift is relaxed, we first benchmarked all methods on synthetic datasets with spatially varying drift fields. In this setting the imposed drift depends on spatial position, so that the observed trajectories cannot be explained by a single global translation or rigid transform. The benchmark therefore probes how well different correction strategies recover latent trajectories when drift varies across the field of view.

Three drift regimes were considered: smooth, piecewise smooth, and piecewise abrupt. These represent progressively more structured spatial drift fields, ranging from globally smooth variation to region-wise drift with either continuous or discontinuous spatial transitions. Concretely, in smooth we generated a drift field as the sum of a temporally smoothed global translation component and a low-frequency spatially varying local component (RBF field with fixed spatial centers and time-varying smooth weights), including a mild smooth depth-dependent trend. In piecewise smooth, we added region-specific drift trajectories and mixed them with soft spatial weights (Gaussian-like blending), yielding continuous transitions between regions. In piecewise abrupt, the same region-wise trajectories were combined using hard nearest-region assignment (one-hot weights), yielding discontinuous spatial boundaries. All seven correction models were evaluated on identical synthetic realizations.

The underlying datasets were generated from the baseline spatially varying drift profile listed in Supplementary Table 1. For each regime, five datasets were generated with *n*_tracks_ = 80 tracks and *n*_times_ = 20 time points, using seed = 42, a stationary fraction of 0.45, a migrating fraction of 0.35, and a drift temporal smoothness of 0.9. This yields mixed populations of stationary and intrinsically migrating tracks, which is essential because a useful drift-correction method must separate externally imposed drift from true biological motion rather than merely reduce global offsets.

Figure 1 walks through one representative smooth realization in full detail. Panels a and b show, respectively, the ground-truth trajectories *X*^true^ = {*x*_true_(*t*)} and the corresponding observed trajectories *X*^obs^ = {*x*_obs_(*t*)} from Eq. (2). In panel b the trajectories are visibly displaced by the imposed spatially varying drift, while panel a provides the unperturbed reference geometry. Panel c then overlays the corrected trajectories *X*^corr^ obtained with the rigid model on top of *X*^true^, using the same track color code and a transparent rendering for the ground truth. This panel serves two purposes. First, it gives the reader a direct geometric impression of what the benchmarked data look like in 3D. Second, it already shows the central limitation of the task: even when a correction appears visually plausible at the global scale, small residual offsets remain because the imposed drift field is not globally rigid.

**Figure 1:**
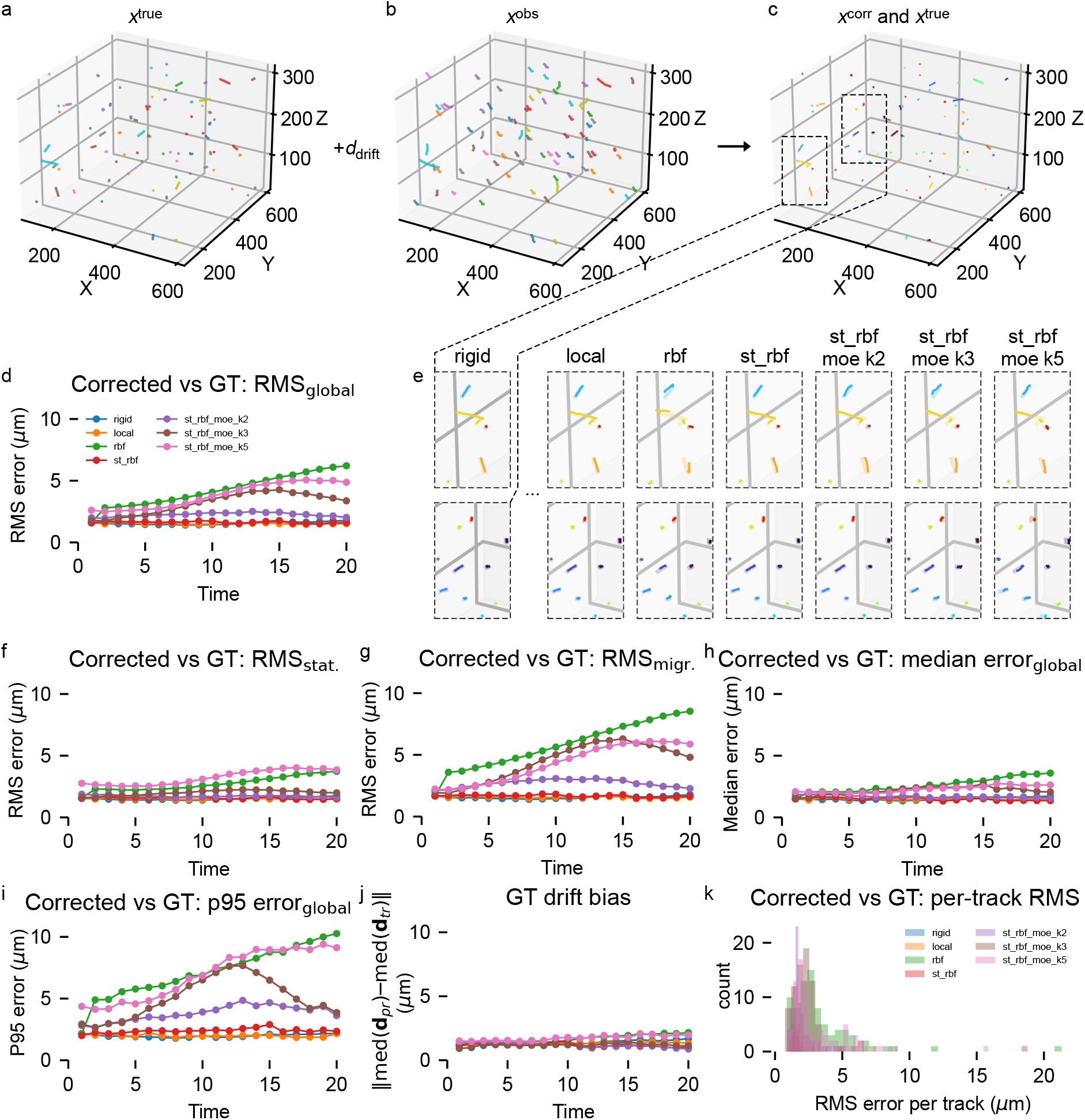
Representative example from the baseline spatially varying drift benchmark. Representative smooth realization from the baseline spatially varying drift profile (Supplementary Table 1), generated with *n*_tracks_ = 80, *n*_times_ = 20, stationary fraction 0.45, migrating fraction 0.35, smoothness 0.9, and seed = 42. (a) Ground-truth trajectories *X*^true^ = {*x*_true_(*t*)} from Eq. (2). (b) Observed trajectories *X*^obs^ = {*x*_obs_(*t*) { after addition of the synthetic spatially varying drift field. (c) Corrected trajectories *X*^corr^ for the rigid method overlaid with the ground truth using the same color code; corrected trajectories are shown opaque and ground-truth trajectories transparent. (d) Global corrected-versus-ground-truth RMS error over time for all benchmarked methods: rigid stepwise model, local community model, per-timepoint RBF model, joint space-time RBF model, and three mixture-of-experts variants of the space-time RBF model. (e) Matched zoom-ins from the highlighted regions in panel c for all seven corrected solutions, shown to visually compare local overlap between corrected and ground-truth segments. (f) RMS error over time restricted to the stationary ground-truth subset. (g) RMS error over time restricted to the migrating ground-truth subset. (h) Global median pointwise corrected-versus-ground-truth error over time. (i) Global *p*95 pointwise corrected-versus-ground-truth error over time. (j) Absolute drift-bias error over time, computed as the absolute difference between recovered and ground-truth drift magnitude. (k) Histogram of per-track RMS error between corrected and ground-truth trajectories across all tracks. All trajectory-error metrics are reported in *µ*m.

Panel d quantifies the same example across all methods using the global pointwise RMS error between corrected and ground-truth coordinates over time. The rigid and local methods have lowest RMS throughout the sequence together with the joint space-time RBF model, and the plain per-timepoint RBF as well as the more flexible mixture-of-experts variants accumulate substantially larger errors as time progresses. Importantly, the curves diverge only gradually. The dominant failure mode in this example is therefore not a sudden catastrophic breakdown at isolated time points, but a systematic accumulation of residual mismatch.

Panel e complements this numerical comparison with local visual inspection. It shows matched zoom-ins for all seven corrected solutions from the same spatial regions highlighted in panel c. These zoomed views are particularly useful because they reveal what the global metrics actually measure. For the rigid and local models, most corrected segments closely follow the ground-truth segments and are usually parallel or only slightly offset. The same holds for `st_rbf`. For `rbf`and, even more clearly, for the mixture-of-experts variants with larger *k*, the corrected segments deviate more strongly from the target geometry, indicating that additional model flexibility does not translate into better recovery in this baseline regime. In other words, the visually best solutions are not the most flexible ones, but the ones that regularize the drift estimate more strongly to avoid absorbing true motion or local noise.

Panels f and g split the global RMS into the stationary and migrating ground-truth subsets. This distinction is essential for interpretation. A method may appear successful when evaluated only on nearly stationary tracks while still failing to preserve genuinely moving trajectories. In the present example, rigid, local, and the space-time RBF model remain comparably strong on both subsets, which indicates that they are not merely aligning static anchors but are preserving moving tracks as well. The plain RBF model shows the clearest degradation on the migrating subset, where its error rises much more strongly than on the stationary subset. This is the signature expected from a model that is too weakly constrained in time and therefore more easily deformed by locally moving tracks.

Panels h and i add two complementary error summaries: the global median error and the global *p*95 error over time. These are important because the RMS alone can be misleading when dominated by a small number of extreme failures. That is not what happens here. For this representative smooth example, the methods that are best in RMS are also best in median and *p*95, and the rank ordering is essentially unchanged. This shows that the lower RMS of the rigid, local, and space-time RBF models is not an artifact of a few fortunate outliers. Rather, their advantage reflects a generally tighter error distribution. Put differently, the residual discrepancy for the best methods is distributed rather uniformly across points, with comparatively small tails, instead of being driven by isolated catastrophic mismatches. This is also why the visual impression from panel e and the numerical summaries tell the same story. The correction is usually close to the ground truth, but it is not exact, and the remaining offset is typically small rather than rare and large.

Panel j reports the absolute drift-bias metric, that is, the discrepancy between recovered and ground-truth drift magnitude over time. This metric is complementary to corrected-versus-ground-truth trajectory error because it focuses on the fidelity of the estimated drift field itself. In the present example, the drift-bias curves remain relatively compact and do not separate methods as strongly as the corrected-trajectory metrics. This is informative, as a method can recover the bulk drift magnitude reasonably well and still produce larger pointwise reconstruction errors if the spatial allocation of that drift is inaccurate or if true motion is partially absorbed into the estimated correction. Drift bias alone is therefore insufficient for model selection, but it is useful for diagnosing whether a method fails primarily by misestimating the global amount of drift or by misplacing it across the spatial field.

Finally, panel k shows the histogram of per-track RMS errors for the same example. This panel confirms the conclusions from panels d, h, and i in distributional form. The rigid, local, and space-time RBF models concentrate more mass at lower per-track errors and exhibit shorter high-error tails, whereas `rbf`and the more flexible MoE variants produce broader distributions with more tracks extending to large RMS values. This again argues against the interpretation that the global differences are caused by only a handful of pathological tracks. The performance gap is population-wide.

Taken together, Figure 1 already reveals the main trend that recurs throughout the full baseline study. In the smooth regime, rigid, local and space-time RBF models are essentially tied at the top, while `rbf`and the MoE extensions perform substantially worse. This point is worth emphasizing because, at first glance, one might expect the more expressive nonlinear models to dominate under spatially varying drift. The opposite happens here. In this baseline regime, added flexibility tends to reduce robustness rather than improve accuracy.

Figure 2 aggregates these comparisons across all five realizations for each of the three baseline drift modes. The heatmaps in panels a to d summarize the mean overall RMS, median error, *p*95 error, and absolute drift bias, respectively, while panel e plots mean RMS against mean *p*95 for all method-mode combinations. Several consistent patterns emerge.

**Figure 2:**
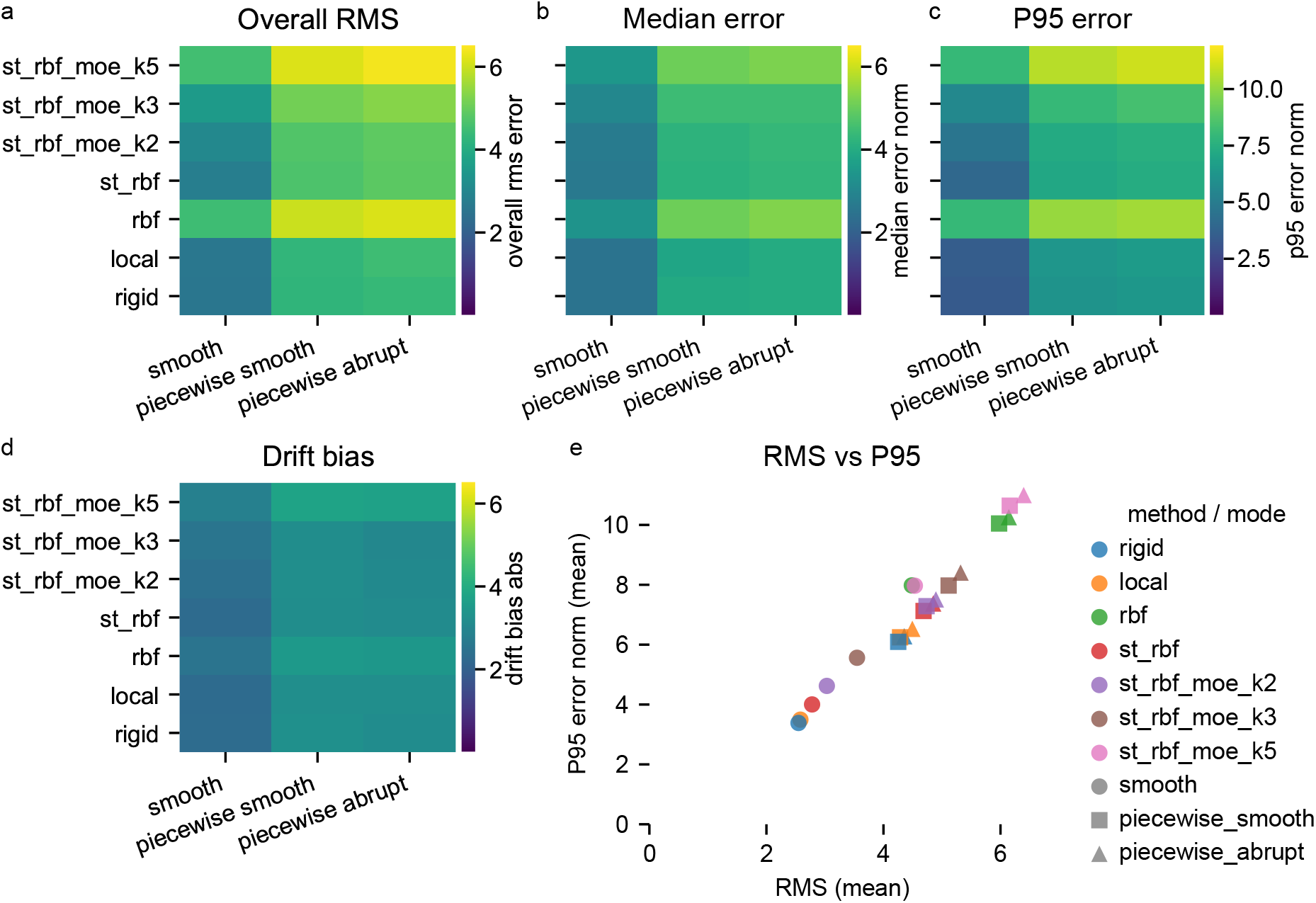
Overall results for the baseline suite across spatially varying drift regimes. Aggregate benchmark summary across the three baseline synthetic drift modes smooth, piecewise smooth, and piecewise abrupt, each evaluated on five independently generated datasets from the baseline spatially varying drift profile. Rows correspond to the seven correction methods rigid stepwise model, local community model, per-timepoint RBF model, joint space-time RBF model, and three mixture-of-experts variants of the space-time RBF model. (a) Mean overall corrected-versus-ground-truth RMS error. (b) Mean overall median pointwise error. (c) Mean overall *p*95 pointwise error. (d) Mean absolute drift-bias error. (e) Mean RMS versus mean *p*95 error for all method-mode combinations, with color indicating method and marker shape indicating drift mode: circles for smooth, squares for piecewise smooth, and triangles for piecewise abrupt. In panel e, points closer to the lower-left corner indicate better overall performance with both lower average reconstruction error and lower tail error. All values are reported in *µ*m and are averaged over the five datasets per mode. The exact numerical values underlying the heatmaps are provided in Table 5.

First, performance degrades monotonically as the drift regime becomes more structured. Across virtually all methods, the smooth mode yields the lowest errors, piecewise smooth increases all trajectory-error metrics, and piecewise abrupt is usually the most difficult or nearly the most difficult case. This confirms that the synthetic generator produces a meaningful hierarchy of difficulty: hard spatial boundaries in the drift field are more challenging to recover than continuously varying fields.

Second, the rigid and local models are the most robust methods across the baseline suite. In the smooth mode, rigid and local achieve the best overall RMS values, 2.548 *µ*m and 2.580 *µ*m, respectively, with identical median error of 2.483 *µ*m and low *p*95 errors of 3.384 *µ*m and 3.495 *µ*m. Their drift-bias values are also low at 2.311 *µ*m and 2.268 *µ*m. In piecewise smooth, they remain the best pair with RMS values of 4.252 *µ*m for rigid and 4.284 *µ*m for local, while in piecewise abrupt rigid remains best at 4.355 *µ*m and local follows closely at 4.493 *µ*m. The same near-tie is visible in median and *p*95 error. Thus, although the benchmarked drift is not globally rigid, the rigid stepwise model remains highly competitive, and in the harder regimes it even becomes marginally superior to the local community model.

Third, the joint space-time RBF model occupies a stable intermediate position. It is consistently better than the plain per-timepoint RBF and most MoE variants, but consistently worse than rigid and local in these overall results. Its overall RMS increases from 2.780 *µ*m in smooth to 4.685 *µ*m in piecewise smooth and 4.850 *µ*m in piecewise abrupt. The corresponding *p*95 errors rise from 4.003 *µ*m to 7.125 *µ*m and 7.373 *µ*m. This pattern suggests that explicit coupling across space and time is helpful relative to a purely time-local nonlinear field, but that the model is still not as robust as the simpler rigid and local estimators in this baseline setting.

Fourth, the plain per-timepoint RBF model and the more flexible MoE variants do not benefit from their added expressivity. In fact, they are often the worst-performing approaches. In the smooth regime, `rbf`and `st_rbf_moe_k5`already reach RMS values around 4.5 *µ*m and *p*95 values near 8 *µ*m, far above the rigid and local baselines. The same ordering persists and even strengthens in the piecewise regimes, where `rbf`reaches RMS values of 5.976 *µ*m and 6.142 *µ*m and *p*95 values of 10.048 *µ*m and 10.257 *µ*m in piecewise smooth and piecewise abrupt, respectively. The *k* = 5 MoE variant is similarly poor and even slightly worse in the abrupt regime. Increasing model complexity therefore does not solve the spatially varying drift problem in this benchmark. It primarily increases susceptibility to overfitting or to misattributing intrinsic track motion to the drift field.

Fifth, the drift-bias heatmap in panel d refines this picture. The ranking in drift bias is notably flatter than the ranking in corrected-versus-ground-truth trajectory error. In piecewise abrupt, for example, `st_rbf_moe_k3`attains the lowest mean drift bias at 3.019 *µ*m, slightly below rigid at 3.150 *µ*m and local at 3.213 *µ*m, yet its overall RMS and *p*95 remain markedly worse. This difference matters conceptually. Recovering the mean drift magnitude reasonably well is not equivalent to reconstructing corrected trajectories accurately. A flexible method may match the gross drift scale while still distributing correction in the wrong spatial regions, or while partially cancelling genuine migration. The trajectory-based metrics are therefore the primary criteria for model selection, whereas drift bias serves as a mechanistic diagnostic.

Panel e of Figure 2 summarizes these findings by plotting the mean RMS against the mean *p*95 error for all method–mode combinations. The points follow a near-monotonic trend from lower left to upper right, indicating that methods with lower mean RMS also consistently exhibit lower tail error. In other words, the ranking of methods is preserved across both average and high-percentile error measures. This agreement shows that the performance differences observed in the benchmark reflect broadly distributed reconstruction accuracy rather than isolated extreme failures. Reporting RMS together with median and *p*95 therefore provides a complementary view of the residual error distribution, capturing both typical reconstruction accuracy and the behavior of the upper error tail.

Overall, the baseline suite yields a clear and somewhat counterintuitive conclusion. Once spatially varying drift is introduced, one might expect highly flexible nonlinear field models to dominate. In the present synthetic benchmark, they do not. The most reliable performance is obtained from the rigid and local models, with the joint space-time RBF model forming a second tier and the plain RBF and larger MoE variants trailing substantially behind. The fact that this ordering persists from smooth to piecewise abrupt indicates that, at least for the baseline spatially varying regimes considered here, robustness to mixed stationary and migrating populations is more important than raw representational flexibility. This benchmark therefore establishes the reference ranking against which the more structured stress-test scenarios in the following subsections can be interpreted.

#### RBF model family robustness and hyperparameter sweeps

The baseline comparison showed that the two RBF-based model families did not outperform the simpler rigid and local estimators under spatially varying drift. The next question was therefore not only whether the RBF family could be improved by tuning, but also whether its behavior was stable and interpretable across reasonable hyperparameter and robustness choices. To address this, we performed a targeted set of sweeps over the principal space-time RBF lengthscales from the anisotropic Gaussian kernel in Eq. (41), robust fitting options based on the ridge objective in Eq. (43), representational extensions of the augmented design in Eq. (42), and the optional stationary-subset fitting strategy.

Figure 3a first examines the central space-time RBF lengthscale pair (*ℓ*_space_, *ℓ*_*t*_) from Eq. (41) while all other settings were held fixed. For each regime, the top row shows the aggregated mean global RMS error 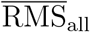 and the bottom row the corresponding mean absolute drift-bias 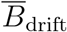. Red markers indicate the minimum of the respective plotted metric. In addition, for the lower row we marked a simple compromise point between reconstruction accuracy and drift recovery by minimizing the combined score

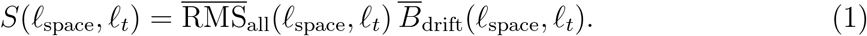

**Figure 3:**
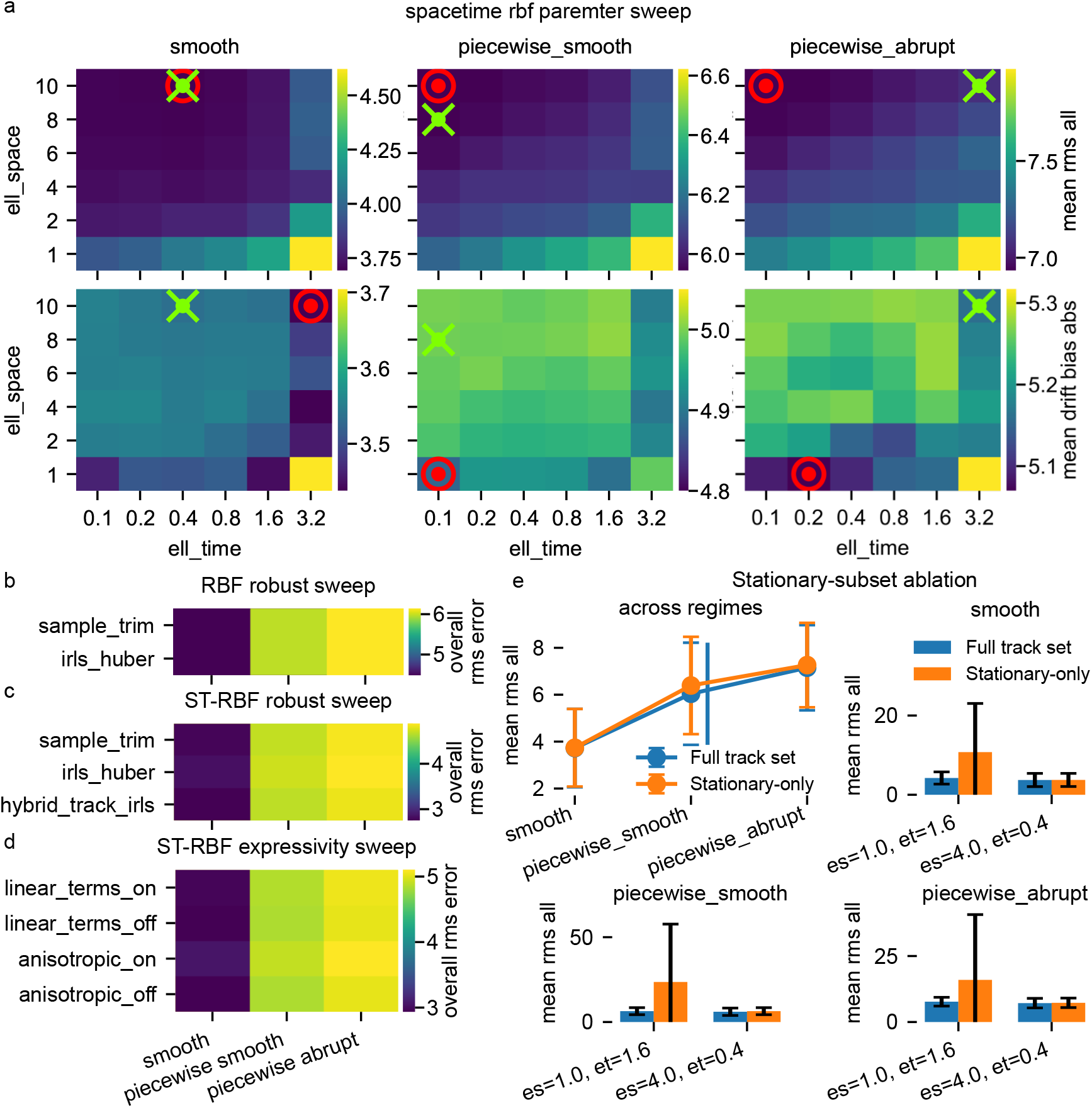
RBF-family robustness analyses and hyperparameter sweeps across baseline spatially varying drift regimes. (a) Space-time RBF lengthscale sweep over ℓ_space_ ∈ {1, 2, 4, 6, 8, 10} and ℓ_*t*_ ∈ {0.1, 0.2, 0.4, 0.8, 1.6, 3.2}, aggregated separately for `smooth`, `piecewise_smooth`, and `piecewise_abrupt`. The top row shows the mean global RMS error 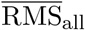 and the bottom row the mean absolute drift-bias 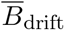, each averaged across benchmark realizations. Red markers indicate the minimum of the plotted metric. Green crosses indicate the best compromise between RMS and drift bias according to the combined score defined in Eq. (1). (b) Per-timepoint RBF robust sweep comparing trimmed ridge fitting and IRLS with Huber weights, summarized by mean overall RMS across regimes. (c) Space-time RBF robust sweep comparing trimmed ridge fitting, IRLS with Huber weights, and track-level trimming combined with IRLS refitting. (d) Space-time RBF expressivity sweep comparing isotropic versus anisotropic spatial kernels and linear-term augmentation on versus off, again summarized by mean overall RMS. (e) Stationary-subset ablation for the practical space-time RBF candidate ( ℓ_space, *t*_) = (4.0, 0.4), comparing training on the full track set versus training only on the heuristically selected stationary subset (top left). Error bars indicate standard deviation across benchmark realizations. Top right to bottom pannels: Mean RMS for two parameter candidates, (ℓ _space, *t*_) = (1.0, 1.6) and (4.0, 0.4), across the three baseline drift regimes. The candidate (1.0, 1.6) is included here to show that stationary-only fitting can become strongly unstable in less favorable parameter regions. All values are reported in *µ*m. *Overall RMS* denotes the dataset-level RMS over all corrected points, and *mean RMS* denotes the average of that quantity across realizations. The complete numerical results underlying the parameter sweeps are provided in Supplementary Tables 6–10.

This heuristic is used to identify parameter regions that are not optimal in RMS alone but remain competitive when drift bias is taken into account simultaneously.

Several trends are immediately visible. In all three regimes, increasing *ℓ*_space_ generally improves the RMS, whereas increasing *ℓ*_*t*_ beyond small values tends to worsen it. For the `smooth`regime, the best RMS occurs at (*ℓ*_space_, *ℓ*_*t*_) = (10.0, 0.4) with 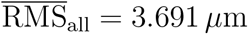 and 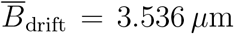. For `piecewise_smooth`, the best RMS lies at (10.0, 0.1) with 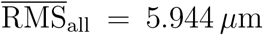 and bias 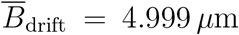, while the best compromise score is essentially tied and shifts only slightly to (8.0, 0.1). For `piecewise_abrupt`, the best RMS is again found at the largest tested spatial scale and smallest temporal scale, (10.0, 0.1), with 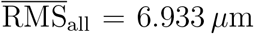, whereas the compromise score shifts toward larger temporal smoothing at (10.0, 3.2) because this slightly reduces bias at the cost of a modest RMS increase. Thus, the dominant trend across regimes is clear: within the tested range, stronger spatial smoothing is beneficial, whereas overly large temporal lengthscales are not.

This trend is also interpretable at the model level. Larger *ℓ*_space_ in the anisotropic kernel of Eq. (41) forces the learned field to vary more smoothly across space, which suppresses local overfitting to individual trajectories. Smaller *ℓ*_*t*_ in the same kernel allows more temporal freedom, which appears advantageous in these synthetic regimes because the drift does vary over time and should not be oversmoothed. Put differently, the present benchmark does not favor highly local spatial deformations that are also strongly coupled across time. It instead favors a comparatively smooth spatial field with only moderate temporal coupling. At the same time, these gains remain limited: even the best grid points do not close the gap to the top-performing rigid and local baselines from Figure 2. Hyperparameter tuning can improve the space-time RBF model within its own family, but it does not change the overall model ranking established by the baseline suite.

We then asked whether the weaker performance of the RBF family might mainly reflect insufficient robustness to outlying correspondences. Figure 3b shows the per-timepoint RBF robust sweep, comparing trimmed ridge fitting on an iteratively updated inlier set and IRLS with Huber-type weights as defined in Eq. (31) and optimized through Eq. (32). Here the result is unambiguous: both configurations yield numerically identical mean overall RMS values in all three regimes, namely 4.484 *µ*m in `smooth`, 5.976 *µ*m in `piecewise_smooth`, and 6.142 *µ*m in `piecewise_abrupt`. For the plain per-timepoint RBF model, changing between these two robustification modes therefore has no measurable practical effect in the present benchmark.

Figure 3c shows the corresponding robustness sweep for the joint space-time RBF model. Here, the outcome is slightly more favorable but still modest in magnitude. The best of the tested variants is track-level trimming combined with IRLS refitting, which reaches mean overall RMS values of 2.750 *µ*m for `smooth`, 4.666 *µ*m for `piecewise_smooth`, and 4.819 *µ*m for `piecewise_abrupt`. The competing variants, namely trimmed ridge fitting and IRLS with Huber-type weights under the objective in Eq. (43), remain very close, with differences typically well below 0.1 *µ*m. Thus, robust fitting details do matter somewhat for the space-time model, but only at the level of fine tuning. They do not alter the broader conclusion that the model remains less reliable than the rigid and local baselines.

The expressivity sweep in Figure 3d addresses a different question, namely whether the space-time RBF underperforms simply because it is not flexible enough. We therefore compared isotropic versus anisotropic spatial scales in Eq. (41) and the presence or absence of additional linear terms in the augmented design of Eq. (42). The result is again negative. The simplest configuration, with anisotropy off and linear terms off, is consistently best or tied for best across all regimes, yielding 2.934 *µ*m in `smooth`, 4.835 *µ*m in `piecewise_smooth`, and 5.023 *µ*m in `piecewise_abrupt`. Turning anisotropy on or appending linear terms slightly increases the error in every regime. In other words, the present failure mode is not lack of representational capacity. Increasing model flexibility within the tested range does not improve reconstruction accuracy and instead tends to make the fit slightly worse.

A final question concerned the optional stationary-subset strategy. Conceptually, this ablation asks whether drift estimation benefits from being anchored only to tracks that are heuristically classified as stationary, rather than using the full track population. The attraction of this idea is clear: if truly migrating tracks contaminate the drift estimate, restricting the fit to stationary anchors could produce a cleaner field. The practical risk is equally clear: if the stationary selector is imperfect, the reduced training set may become too small or too biased, leading to unstable estimation.

Figure 3e (top left) shows this comparison for the practically relevant candidate (*ℓ*_space_, *ℓ*_*t*_) = (4.0, 0.4), using all tracks versus a stationary-only training subset. Across all three regimes, the stationary-only variant is not better. In `smooth`, the mean RMS changes only marginally from 3.723 *µ*m to 3.737 *µ*m when the stationary-only subset is used. In `piecewise_smooth`, the mean RMS increases from 6.040 *µ*m to 6.393 *µ*m, and in `piecewise_abrupt`from 7.151 *µ*m to 7.263 *µ*m. The corresponding drift-bias values also remain unchanged or worsen slightly. Hence, even for the better of the two tested parameter candidates, stationary-only fitting provides no robust gain.

The full ablation results are shown in the remaining panels of Figure 3e, where the second candidate (1.0, 1.6) reveals the more severe failure mode of this strategy. For that parameter pair, switching to the stationary-only subset causes large increases in both mean error and variance. In `smooth`, the mean RMS rises from 4.230 *µ*m to 10.756 *µ*m. In `piecewise_smooth`, it rises from 6.396 *µ*m to 23.586 *µ*m, and in `piecewise_abrupt`from 7.694 *µ*m to 15.960 *µ*m, accompanied by extremely large standard deviations. This is strong evidence that the stationary-preselection idea is not robust in the present setting and can fail catastrophically in unfavorable parameter regions.

Taken together, these sweeps lead to a compact set of conclusions. First, the space-time RBF model does have an interpretable lengthscale dependence, with larger *ℓ*_space_ and comparatively small *ℓ*_*t*_ performing best within the tested grid. Second, robustification choices and additional expressivity options change the results only marginally and do not rescue the RBF family from its inferior ranking relative to rigid and local correction. Third, fitting on a heuristically selected stationary subset is not beneficial here and can be strongly destabilizing. This supports using the simpler, non-ablated space-time RBF configuration without stationary-only fitting as the default representative of the RBF family.

#### Baseline comparison across structured stress-test regimes

We next asked whether the ranking established in the baseline spatially varying suite remains stable once the synthetic drift becomes more explicitly structured. To this end, we evaluated the same seven correction models, namely the rigid stepwise model (Eq. (5), Eq. (11)), the local community model (Eq. (21), Eq. (22)), the per-timepoint RBF model (Eq. (25), Eq. (36)), the joint space-time RBF model (Eq. (41), Eq. (43), Eq. (44)), and three mixture-of-experts extensions of the space-time RBF model (Eq. (45), Eq. (46), Eq. (47)), on a second set of synthetic stress regimes. Unlike the previous suite, these regimes target specific mechanistic departures from globally smooth drift: rotation drift introduces time-varying rotational components around a reference center, piecewise spatial drift imposes region-specific drift trajectories with stronger spatial divergence, and jump events introduce temporally discontinuous offsets, optionally including rotational jumps. The purpose of this benchmark is to test whether the simpler models fail in predictable ways once the drift contains structured rotational, discontinuous, or strongly region-specific components. The underlying datasets were generated from the preset structured non-rigid drift profile listed in Supplementary Table 1, with *n*_tracks_ = 80, *n*_times_ = 20, seeds 42 to 51, stationary fraction 0.45, migrating fraction 0.35, and regime-specific preset controls for spatial transition width, drift scale, and measurement noise.

Figure 4 summarizes the aggregate results across these three stress regimes. Panels a to d report mean overall RMS, mean median error, mean *p*95 error, and mean absolute drift bias, while panel e plots mean RMS against mean *p*95 for all method–mode combinations. Several points stand out immediately.

**Figure 4:**
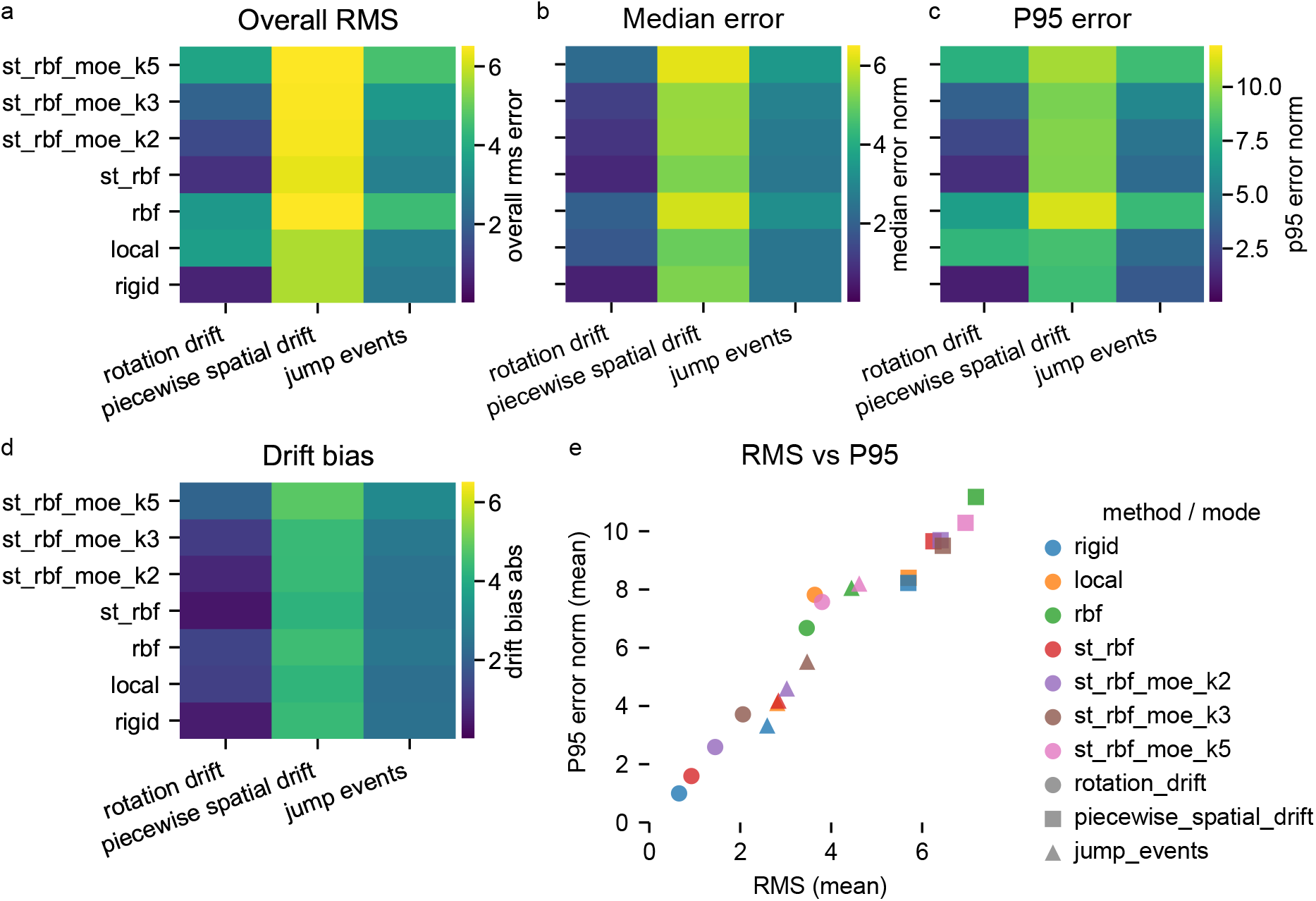
Overall results for the structured non-rigid drift stress-test suite. Aggregate benchmark summary across the three structured synthetic drift modes rotation drift, piecewise spatial drift, and jump events. Rows correspond to the seven correction methods rigid stepwise model, local community model, per-timepoint RBF model, joint space-time RBF model, and three mixture-of-experts variants of the space-time RBF model. (a) Mean overall corrected-versus-ground-truth RMS error. (b) Mean overall median pointwise error. (c) Mean overall *p*95 pointwise error. (d) Mean absolute drift-bias error. (e) Mean RMS versus mean *p*95 error for all method–mode combinations, with color indicating method and marker shape indicating drift mode: circles for rotation drift, squares for piecewise spatial drift, and triangles for jump events. In panel e, points closer to the lower-left corner indicate better overall performance with both lower average reconstruction error and lower tail error. All values are reported in *µ*m and are averaged across benchmark realizations for the respective mode.

First, the qualitative ranking from the baseline spatially varying suite is only partly preserved. In the previous benchmark, the rigid and local models formed the strongest pair across all three modes, the joint space-time RBF model occupied a stable second tier, and the per-timepoint RBF and larger mixture-of-experts variants trailed behind. Under the present stress regimes, this pattern becomes more differentiated. The rigid stepwise model remains among the top performers in all three regimes and is the clearest winner in rotation drift. By contrast, the local community model is no longer uniformly near-tied with rigid. Its mean RMS in rotation drift is visibly much larger than that of the rigid model and also clearly above the joint space-time RBF model, indicating that this regime exposes a structural weakness of the local community approach. This is plausible from the model definition in Eq. (21): the method blends locally estimated displacement traces in space, which is well matched to smoothly varying translational fields, but less naturally matched to globally coherent rotational structure.

Second, rotation drift produces the strongest ranking shift relative to the previous baseline suite. Here the rigid stepwise model performs best by a wide margin, with the joint space-time RBF model forming the second-best solution and the smaller mixture-of-experts variants following behind. The local community model, the per-timepoint RBF model, and the largest mixture-of-experts variant all show substantially larger average and tail errors. This differs qualitatively from the earlier smooth and piecewise smooth regimes, where rigid and local were nearly tied. The difference is informative: a globally coherent rotational component remains compatible with the rigid formulation in Eq. (5), whereas a spatially blended drift representation or highly flexible nonlinear field can distribute the correction less efficiently and thereby incur larger residuals.

Third, piecewise spatial drift is the most difficult regime in the present figure, and indeed appears more challenging than the earlier piecewise smooth and piecewise abrupt baseline regimes. All methods shift upward in RMS, median, and *p*95 error, and the heatmaps show a compressed but consistently poor ranking across the full model family. Even here, however, the simpler models remain strongest: rigid and local still occupy the lowest-error region, while the joint space-time RBF model and the mixture-of-experts variants remain clearly above them, and the per-timepoint RBF model is again among the weakest performers. This is an important negative result. One might expect the more flexible nonlinear field models, especially the mixture-of-experts extension in Eq. (47), to benefit from strongly region-dependent drift. In these synthetic stress tests, they do not. The same general conclusion from the first suite therefore persists: added representational flexibility does not translate into improved reconstruction accuracy.

Fourth, jump events occupies an intermediate position. It is more difficult than rotation drift, but clearly less severe than piecewise spatial drift. In this regime the rigid model again remains at the top, while the local community model, the joint space-time RBF model, and the smaller mixture-of-experts variants cluster somewhat above it. The per-timepoint RBF model and the largest mixture-of-experts variant remain worst or near-worst. This regime is especially relevant conceptually because temporal discontinuities challenge the assumption of smoothly varying drift. Yet the space-time RBF family, despite explicit temporal modeling via the four-dimensional kernel in Eq. (41), does not gain a decisive advantage. The simplest interpretation is that explicit temporal flexibility alone is insufficient when the correction model is still prone to spatial misallocation or to partial absorption of genuine object motion.

The drift-bias heatmap in panel d adds an important nuance. As in the previous baseline suite, the method ranking in drift bias is flatter than the ranking in corrected-versus-ground-truth trajectory errors. In piecewise spatial drift, for example, multiple methods attain broadly similar bias values while remaining clearly separated in RMS and *p*95. This again shows that recovering the gross drift magnitude is not equivalent to reconstructing the corrected trajectories accurately. A model may estimate a plausible average drift scale while still distributing that correction across the wrong spatial regions or overfitting intrinsically migrating tracks. Drift bias therefore remains a useful mechanistic diagnostic, but the trajectory-based metrics remain the primary basis for model comparison.

Panel e reinforces this interpretation. As in Figure 2, the points follow a near-monotonic trend from lower left to upper right, showing that methods with lower mean RMS also tend to have lower high-percentile error. The ranking is therefore consistent across central and tail-sensitive error summaries. What changes relative to the previous suite is not the internal consistency of the metric set, but the relative placement of the methods. The rigid model moves even more clearly into the lower-left corner for rotation drift, whereas the local community model shifts upward in that same regime. For piecewise spatial drift, all methods move toward the upper-right corner, confirming that this regime acts as the strongest stress test in the present block.

Compared with the earlier baseline suite, these results therefore sharpen the interpretation of model strengths and failure modes. The rigid stepwise model is not merely competitive under mildly spatially varying drift, but remains remarkably robust even under explicitly structured non-rigid regimes, particularly when the dominant distortion includes coherent rotational or jump-like components. The local community model remains strong for region-wise translational heterogeneity, but is less robust than rigid when the drift structure becomes rotational. The joint space-time RBF model continues to occupy an intermediate tier: it is usually better than the plain per-timepoint RBF model and often better than the larger mixture-of-experts variants, but it still does not surpass the simpler baselines. Finally, the per-timepoint RBF model and the more expressive mixture-of-experts variants again fail to convert flexibility into accuracy, even in the very regimes where one might have expected them to benefit most.

We also repeated the robustness and hyperparameter analyses for the RBF family under these structured stress regimes (Supplementary Figure 13). The qualitative conclusions remained unchanged: tuning, robust fitting variants, and additional expressivity produced only marginal differences and did not alter the overall ranking relative to the simpler baselines. The corresponding sweep summaries are therefore provided in Supplementary Information rather than in the main text.

Taken together, the structured non-rigid stress tests do not overturn the core conclusion of the previous benchmark block. Instead, they refine it. The rigid stepwise model emerges as the most consistently reliable method across the full range of synthetic conditions considered here, the local community model remains highly competitive but somewhat more regime-dependent, the joint space-time RBF model forms a persistent second tier, and the more flexible RBF-based variants remain limited by reduced robustness rather than insufficient representational capacity.

### Synthetic point-cloud association benchmarks

The second major benchmark block addresses the reconstruction of object identities from time-resolved point clouds. In contrast to the drift-correction benchmarks, the central question here is not how accurately a geometric distortion field can be estimated and removed, but how reliably individual detections can be linked across time into consistent pseudo-tracks. This problem remains nontrivial even in the absence of drift, because nearby candidates may compete for the same continuation, motion scales may vary across datasets, and appearance, disappearance, or intermittent ambiguity can fragment otherwise coherent trajectories.

The following benchmark block is therefore organized as a progressive increase in association difficulty. We begin with clean no-drift settings that isolate the core assignment problem and establish a broad baseline ranking across local, motion-aware, global, and tracklet-based association strategies. We then ask how these methods behave when the relevant displacement scale is no longer known a priori, and whether uncertainty-aware gating improves robustness under increasingly heterogeneous conditions. Finally, we examine compositional stress scenarios designed to test how well the strongest association strategies tolerate mixed motion statistics, density-induced ambiguity, and more realistic departures from the simplest synthetic setting. Together, these experiments identify which association principles remain reliable as the point-cloud linking problem moves from controlled to progressively more challenging regimes.

#### Baseline comparison

We first established a broad baseline ranking of association strategies under the cleanest possible synthetic conditions, namely in the absence of imaging drift and with a fixed displacement scale per dataset. The purpose of this benchmark is to isolate the identity-reconstruction problem itself before introducing additional complications such as unknown motion scales, heteroscedasticity, compositional stress, or drift-contaminated geometry. In this setting, the observed point clouds follow

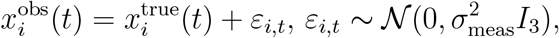

with *σ*_meas_ = 0.2 *µ*m and zero imposed drift. For each fixed displacement scale *d*_*x*_ ∈ {0.2, 0.5, 1, 2, 4} *µ*m, we generated three synthetic datasets with *N*_tracks_ = 80 and *T* = 20 time points. Motion followed a temporally smoothed stochastic process with smoothness 0.9, with mixed populations of stationary, slow, and migrating objects. All methods were evaluated on the same realizations.

The benchmark compared a broad family of association approaches. These included frame-local Hungarian assignment under the gated objective in Eqs. 55–57 with purely geometric costs from Eq. (48), variants using Huberized and Mahalanobis-type costs from Eqs. 49 and 50, constant-velocity refinement based on Eqs. 58 and 59, Kalman constant-velocity association in innovation space using Eqs. 74–77, the global minimum-cost path-cover formulation from Eqs. 81–82, and the tracklet-first stitching strategy based on Eqs. 84–86. The central evaluation metric throughout this association study is the mean link-*F*_1_ score (Eq. (108)), because it directly measures whether consecutive correspondences are correct. By contrast, geometric RMS can remain small even when the identity assignment is poor, for example if a method becomes overly conservative and leaves difficult links unmatched.

Figure 5a shows a representative blind input point cloud assembled from all frames after removing the ground-truth track identities. The underlying synthetic ground-truth trajectories are shown separately in Supplementary Figure 14. Panel b then illustrates two reconstructions from the same blind point cloud. On the left, the frame-local Hungarian assignment with constant-velocity refinement and coordinate-space gating reconstructs trajectories that almost perfectly overlap the transparent ground truth, yielding a mean link-*F*_1_ of 0.99. On the right, Kalman constant-velocity association with innovation gating performs much worse in this example, with a mean link-*F*_1_ of 0.81. These visualizations are useful because they show what the numerical metrics mean in practice. A score near 0.99 still allows small local failures, but the reconstructed tracks remain almost entirely consistent with the ground truth. A score near 0.81 already corresponds to clearly visible reconstruction damage, including broken trajectories, missing continuations, and incorrect continuation structure. Black points indicate detections that were not linked consecutively into a reconstructed trajectory, while color changes within an otherwise continuous ground-truth path indicate fragmentation or identity inconsistency. Additional 3D reconstruction examples for all methods are shown in Supplementary Figure 14.

**Figure 5:**
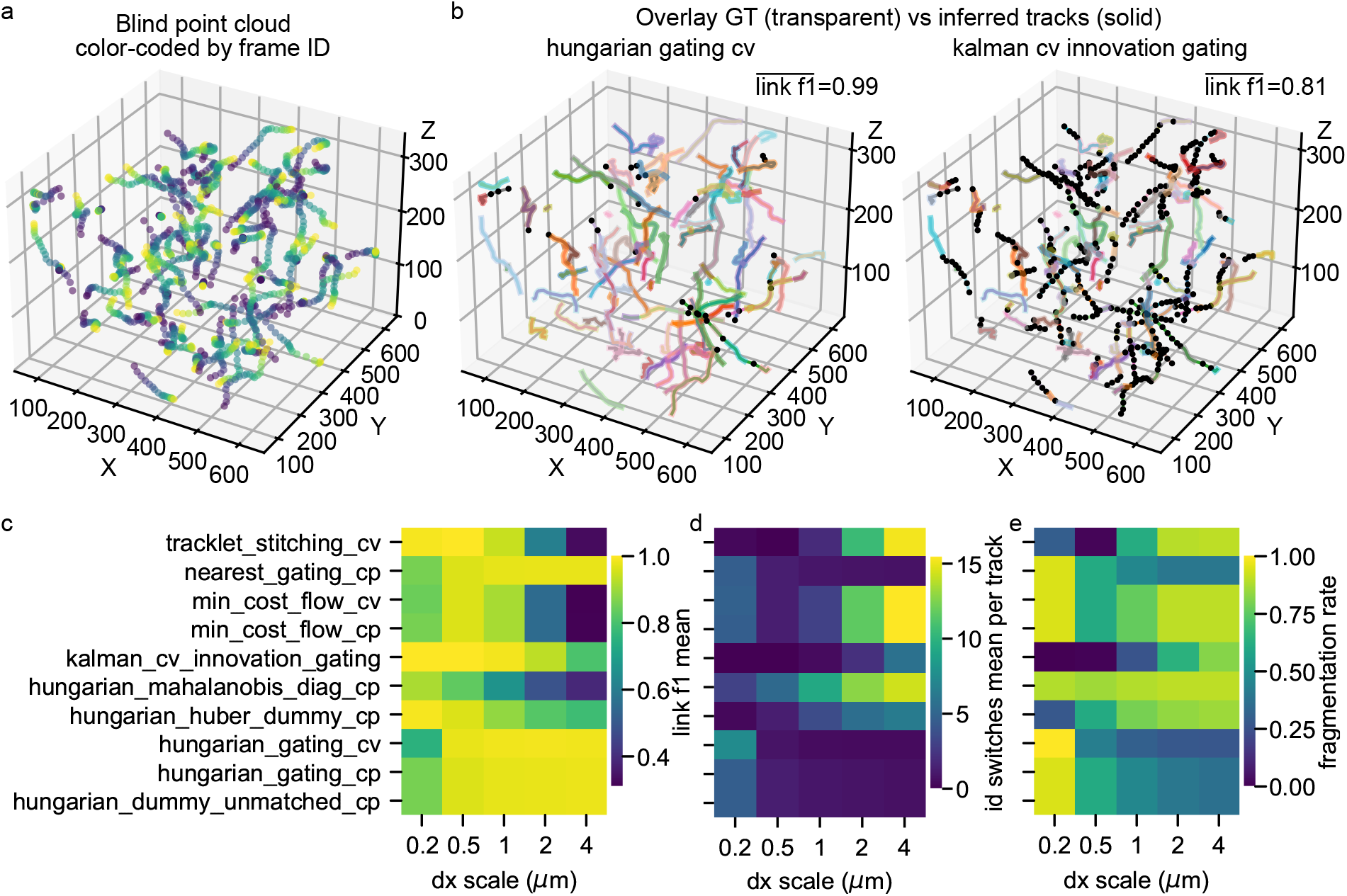
Baseline comparison of point-cloud association strategies under no-drift conditions. Synthetic benchmark for identity reconstruction from unordered 3D point clouds in the absence of imaging drift. For each fixed displacement scale *d*_*x*_ ∈ {0.2, 0.5, 1, 2, 4} *µ*m, three datasets were generated with *N*_tracks_ = 80, *T* = 20, measurement noise *σ*_meas_ = 0.2 *µ*m, temporal smoothness 0.9, and mixed stationary and migrating populations. (a) Representative blind point cloud assembled from all frames after removal of track identities, color-coded by frame index. The corresponding ground-truth trajectories are shown in Supplementary Figure 14. (b) Two representative reconstructions overlaid with the transparent ground truth. Left: frame-local Hungarian assignment with constant-velocity refinement and coordinate-space gating, illustrating near-perfect recovery with mean link-*F*_1_ = 0.99. Right: Kalman constant-velocity association with innovation gating, illustrating visibly degraded recovery with mean link-*F*_1_ = 0.81. Reconstructed tracks are shown solid; ground-truth trajectories are shown transparent and thicker. Black points indicate detections that were not consecutively linked into a reconstructed trajectory. (c) Mean link-*F*_1_ score across methods and displacement scales, where higher values indicate better identity reconstruction. (d) Mean identity switches per track, where lower values are better. (e) Fragmentation rate, where lower values indicate fewer broken trajectories. Methods include frame-local Hungarian assignment under geometric costs and gating, constant-velocity refinement, Kalman innovation gating, global minimum-cost path cover, nearest-neighbor gating, and tracklet-first stitching. All heatmap values are averaged across the three synthetic realizations per displacement scale.

The aggregate results in Figure 5c–e reveal a strikingly non-monotonic dependence on displacement scale. For the smallest displacement *d*_*x*_ = 0.2 *µ*m, Kalman constant-velocity association with innovation gating is best in mean link-*F*_1_, reaching 1.00, while the two-stage tracklet-first stitching variant and the Hungarian method with Huberized cost and explicit unmatched modeling also perform nearly perfectly with 0.991. By contrast, the purely geometric Hungarian baselines, nearest-neighbor gating, and the minimum-cost path-cover variants achieve only about 0.86 in this regime. This small-displacement case is therefore dominated by over-linking risk: when successive observations move only slightly relative to measurement noise, conservative motion-aware methods that normalize by predictive uncertainty can suppress many false links.

At intermediate displacement scales the ranking changes substantially. For *d*_*x*_ = 0.5 *µ*m and 1 *µ*m, the frame-local Hungarian assignment with constant-velocity refinement and coordinate-space gating becomes one of the strongest methods, reaching link-*F*_1_ values of 0.977 and 0.985, respectively. The plain geometric Hungarian and nearest-neighbor baselines remain close behind at these scales, each reaching about 0.962 at 0.5 *µ*m and 0.974 at 1 *µ*m. Kalman innovation gating remains competitive and is still best or tied at low to moderate displacement, but its relative advantage has largely disappeared by *d*_*x*_ = 1 *µ*m. The practical interpretation is that once the displacement becomes large enough to yield informative local motion direction, a simple constant-velocity refinement layered onto Hungarian matching is sufficient and highly effective.

For the larger displacement scales *d*_*x*_ = 2 and 4 *µ*m, the frame-local Hungarian method with constant-velocity refinement and coordinate-space gating becomes the best overall performer, with link-*F*_1_ values of 0.987 and 0.986, respectively. In this regime it clearly outperforms Kalman innovation gating, whose link-*F*_1_ drops to 0.930 and then to 0.805. The geometric Hungarian and nearest-neighbor baselines remain surprisingly robust, staying near 0.978 at 2 *µ*m and only slightly below that at 4 *µ*m. By contrast, the more elaborate global strategies deteriorate strongly: the minimum-cost path-cover methods collapse from around 0.91 at 1 *µ*m to about 0.55 at 2 *µ*m and near 0.31 at 4 *µ*m, while the tracklet-first stitching approach drops from 0.943 at 1 *µ*m to 0.607 and then 0.329. The diagonal Mahalanobis-cost variant performs worst overall, decreasing almost monotonically from 0.911 at 0.2 *µ*m to 0.388 at 4 *µ*m.

The subsidiary diagnostics in panels d and e clarify why these rankings emerge. The Kalman innovation-gating method achieves the lowest fragmentation rate in the low-displacement regime, essentially zero at *d*_*x*_ = 0.2 *µ*m and still very low at *d*_*x*_ = 0.5 *µ*m, which explains its perfect or near-perfect link-*F*_1_ there. However, its identity-switch count rises sharply with increasing displacement, from almost zero at small *d*_*x*_ to more than 5 switches per track at *d*_*x*_ = 4 *µ*m. The opposite pattern is seen for the Hungarian constant-velocity refinement with coordinate-space gating: it starts with relatively high fragmentation and identity-switch values at the smallest displacement, but these decrease rapidly as the motion scale increases, eventually reaching some of the lowest switch and fragmentation values in the benchmark. The broad message is that innovation-normalized filtering is advantageous when the displacement scale is very small and ambiguity is dominated by noise, whereas a simpler coordinate-space formulation with explicit velocity extrapolation becomes more robust once the actual inter-frame motion is sufficiently informative.

The global and tracklet-based methods reveal a second important lesson. Their poor high-*d*_*x*_ performance is not primarily a small geometric mismatch problem, but an identity-consistency failure. This is visible in the strong rise of both fragmentation and identity switches with increasing displacement. In other words, once local ambiguity grows, globally optimizing over candidate link structures or stitching conservative short tracklets does not automatically improve identity recovery. In the present synthetic setting, early wrong decisions appear to propagate rather than being repaired by the global objective. This makes these methods much less suitable as primary baselines for the subsequent stress tests.

Taken together, this baseline comparison establishes two key conclusions for the remainder of the association study. First, link-*F*_1_ indeed provides the clearest ranking criterion, because it tracks the visually apparent reconstruction quality much more faithfully than a purely geometric residual would. Second, among the tested families, frame-local Hungarian association with constant-velocity refinement and coordinate-space gating emerges as the most consistently strong and scalable baseline across the full displacement range, while Kalman innovation gating is highly competitive only in the very low-displacement regime. This is why the subsequent subsections focus more narrowly on the Hungarian constant-velocity family and its gating variants: it offers the best overall tradeoff between assignment accuracy, robustness across displacement scales, and algorithmic simplicity.

#### Association under unknown displacement scale

The baseline comparison above assumed that each synthetic dataset was generated with a fixed displacement scale *d*_*x*_, so that the association problem could be analyzed under controlled and homogeneous motion conditions. We next asked whether the strongest frame-local Hungarian constant-velocity variants remain reliable once this assumption is removed. To this end, we generated no-drift datasets in which the displacement amplitude varies across tracks and time within the same sequence,

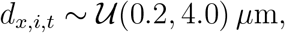

so that no single global motion scale is available at inference time. This setting is substantially more realistic than the fixed-*d*_*x*_ benchmark because the admissible displacement scale must now be inferred adaptively from the data rather than supplied implicitly by the dataset design.

The benchmark was restricted to the Hungarian constant-velocity family, motivated by the previous subsection. In particular, we compared six variants built on the iterative constant-velocity refinement scheme in Eqs. 58 and 59: coordinate-space gating with an adaptive distance threshold from Eq. (66), innovation-based gating using the normalized innovation statistics in Eqs. 61, 65, and 67, the optional directional switch penalty from Eq. (69), and the two-stage assignment strategy from Eq. (70). Across variants, we additionally swept the number of motion-refinement iterations *n*_ref_ ∈ {0, 1, 2, 5}, and for the innovation-based variants we swept the initialization scale of the innovation covariance relative to the nearest-neighbor scale in the initial frame. For each method, the final summary metrics shown in Figure 6b correspond to the parameter configuration that maximized mean link-*F*_1_ across the five synthetic realizations.

**Figure 6:**
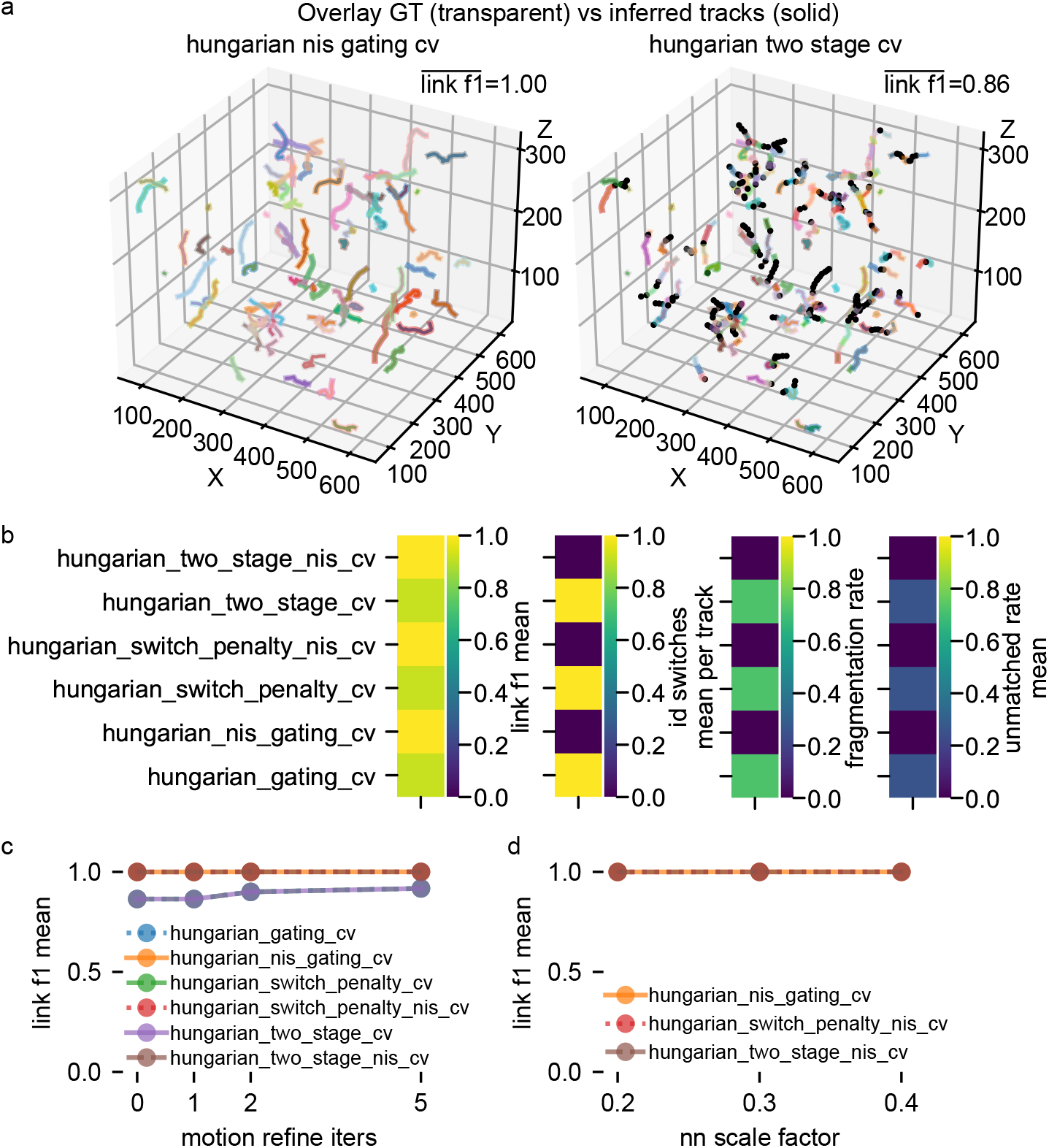
Association under unknown and heterogeneous displacement scale within the Hungarian constant-velocity family. Synthetic no-drift benchmark in which the displacement amplitude varies by track and time within the same dataset, so that no single global motion scale is available at inference time. Five datasets were generated with *N*_tracks_ = 80, *T* = 20, measurement noise *σ*_meas_ = 0.2 *µ*m, and heteroscedastic motion amplitudes *d*_*x,i,t*_ ∼ *U* (0.2, 4.0) *µ*m. All methods are variants of frame-local Hungarian assignment with constant-velocity refinement, differing in gating space and optional penalties or staging. (a) Representative qualitative reconstruction examples. Left: Hungarian constant-velocity association with innovation-based gating, illustrating near-perfect recovery with mean link-*F*_1_ = 1.00. Right: two-stage Hungarian constant-velocity association with coordinate-space gating, illustrating a substantially poorer reconstruction with mean link-*F*_1_ = 0.86. Ground-truth trajectories are shown transparent and slightly thicker; inferred tracks are shown solid. Black points indicate detections that were not linked into consecutive pseudo-tracks. (b) Final method comparison after selecting, for each method, the parameter configuration with highest mean link-*F*_1_ across benchmark realizations. Shown are mean link-*F*_1_ (higher is better), mean identity switches per track (lower is better), fragmentation rate (lower is better), and mean unmatched rate (lower is better). (c) Sensitivity of mean link-*F*_1_ to the number of motion-refinement iterations. The upper, nearly overlapping curves correspond to the innovation-based variants, whereas the lower curves correspond to the coordinate-space variants. (d) Sensitivity of mean link-*F*_1_ to the initialization scale factor of the innovation covariance for the innovation-based methods. Across the tested range, innovation-based variants remain essentially perfect. All values are averaged across the five synthetic realizations.

Figure 6a again provides a qualitative anchor. The left panel shows the best-performing reconstruction, obtained with Hungarian constant-velocity association using innovation-based gating. The inferred tracks almost perfectly coincide with the transparent ground truth, and the mean link-*F*_1_ reaches 1.00. The right panel shows one of the weakest variants, namely the two-stage Hungarian constant-velocity procedure with coordinate-space gating, for which the reconstruction is visibly fragmented and the mean link-*F*_1_ drops to 0.86. As in the previous subsection, black points denote detections that were not linked into consecutive pseudo-tracks. This side-by-side example makes clear that, under heterogeneous and unknown displacement scales, the dominant failure mode is no longer primarily a mild local mismatch but a clear loss of continuity and consistency in the reconstructed identities.

The quantitative comparison in Figure 6b shows a clean separation between two method groups. The three innovation-based variants, namely Hungarian constant-velocity association with innovation gating, the same method with an additional directional switch penalty, and the two-stage Hungarian constant-velocity strategy with innovation gating, all achieve essentially perfect mean link-*F*_1_ values of 1.00. Their identity-switch counts and fragmentation rates are simultaneously near zero. In contrast, the three coordinate-space variants remain substantially weaker. The plain coordinate-space Hungarian constant-velocity method reaches a mean link-*F*_1_ around 0.93, the switch-penalty variant remains at a similar level, and the two-stage coordinate-space variant is worst at about 0.86. These poorer scores are accompanied by markedly higher identity-switch and fragmentation rates. Thus, once the displacement scale is no longer fixed and globally known, the main benefit does not come from additional heuristic penalties or staged assignment alone. It comes from normalization of candidate residuals by an inferred innovation scale.

This shift relative to the previous subsection is conceptually important. Under fixed-*d*_*x*_ conditions, Hungarian constant-velocity association with coordinate-space gating performed extremely well across a broad displacement range and became the preferred baseline because of its simplicity and robustness. The present benchmark shows that this conclusion does not fully transfer to heterogeneous within-sequence motion scales. In that setting, a Euclidean gate, even when adapted through Eq. (66), still relies on a coordinate-space notion of plausibility whose absolute scale can vary substantially across transitions and across local motion states. By contrast, the innovation-based gate in Eqs. 65 and 67 normalizes residuals by an estimated covariance scale and therefore provides a dimensionless criterion that is much less sensitive to such heterogeneity. The improvement is not marginal but decisive: innovation-based gating eliminates almost all remaining association errors in this benchmark.

Panels c and d further clarify the parameter dependence. Panel c shows the mean link-*F*_1_ as a function of the number of motion-refinement iterations. The three innovation-based variants remain at 1.00 across the full tested range, already from zero or one refinement iteration onward. This indicates that, once the innovation-space normalization is in place, the precise number of additional refinement passes is largely irrelevant in the present synthetic setting. The coordinate-space variants behave differently. Their performance starts visibly lower and improves only moderately with more refinement iterations, remaining clearly separated from the innovation-based group. The two-stage coordinate-space method is consistently worst across all iteration counts. This suggests that extra iterative refinement cannot compensate for a mismatch in the gating space itself.

Panel d shows the sensitivity of the innovation-based methods to the initialization scale of the innovation covariance. Again, the result is strikingly stable. Across all tested nearest-neighbor scale factors, the three innovation-based variants remain at a mean link-*F*_1_ of 1.00. In other words, once gating is expressed in normalized innovation space, even a fairly coarse initialization of the innovation scale is sufficient. This stability is highly desirable for practical use, because it implies that the method does not depend on delicate manual calibration of a global displacement threshold.

The directional switch penalty and two-stage assignment deserve brief comment. In the present benchmark, neither produces a meaningful gain once innovation-based gating is already enabled. The switch-penalty innovation variant and the two-stage innovation variant are essentially tied with the simpler Hungarian constant-velocity method using innovation gating alone. Their main value is therefore not improved peak performance under the present conditions, but methodological confirmation that the strong results are not dependent on one very specific implementation detail. Conversely, when combined only with coordinate-space gating, neither modification is sufficient to recover the lost performance. The bottleneck is therefore the unknown and heterogeneous motion scale itself, not merely ambiguous local direction changes or the absence of a staged decision rule.

Taken together, this benchmark refines the candidate selection from the baseline benchmark. Under fixed and known displacement scales, coordinate-space Hungarian constant-velocity association already performed very well and was attractive as a simple default. Under unknown and heterogeneous displacement scales, however, the decisive improvement comes from innovation-based normalization. The innovation-gated Hungarian constant-velocity family remains essentially perfect across all tested parameter settings and therefore emerges as the most robust frame-local association strategy under more realistic assumptions. This result motivates the subsequent focus on statistically normalized innovation gating when the synthetic conditions are made still more heterogeneous and compositionally challenging.

#### Robustness under compositional stress

The previous benchmark showed that innovation-based Hungarian constant-velocity association remains essentially perfect when the displacement scale is unknown but no drift, no birth or death events, and no strong density stress are present. We next asked whether this conclusion survives under substantially more realistic compositional stress. To this end, we evaluated the same six Hungarian constant-velocity variants under three progressively harder conditions: first, a no-drift regime with birth and death events together with a mixed population of stationary and migrating objects; second, the same regime with additional global drift contamination; and third, a density stress test in which the number of tracks increases at fixed volume. Across all three benchmarks, the association family was again based on constant-velocity refinement via Eqs. 58 and 59, with either coordinate-space gating from Eq. (64) or innovation-based gating from Eqs. 61, 65, and 67, optionally augmented by the switch penalty in Eq. (69) or the two-stage strategy in Eq. (70).

Figures 7 and 8 summarize the first two stress regimes. In the birth/death plus stationary-mix benchmark without drift, the three innovation-based variants are again clearly superior. Hungarian constant-velocity association with innovation gating, the same method with an added switch penalty, and the two-stage innovation-gated variant all reach the same best mean link-*F*_1_ of 0.990. At the same time, their mean identity-switch counts are only 0.138 per track, fragmentation is only 0.128, and the unmatched rate is approximately 0.013. The qualitative overlay in Figure 7a is consistent with these values: the innovation-gated reconstruction remains almost perfectly aligned with the transparent ground truth, whereas the weaker coordinate-space variant exhibits visibly more black unmatched detections and more broken trajectories.

**Figure 7:**
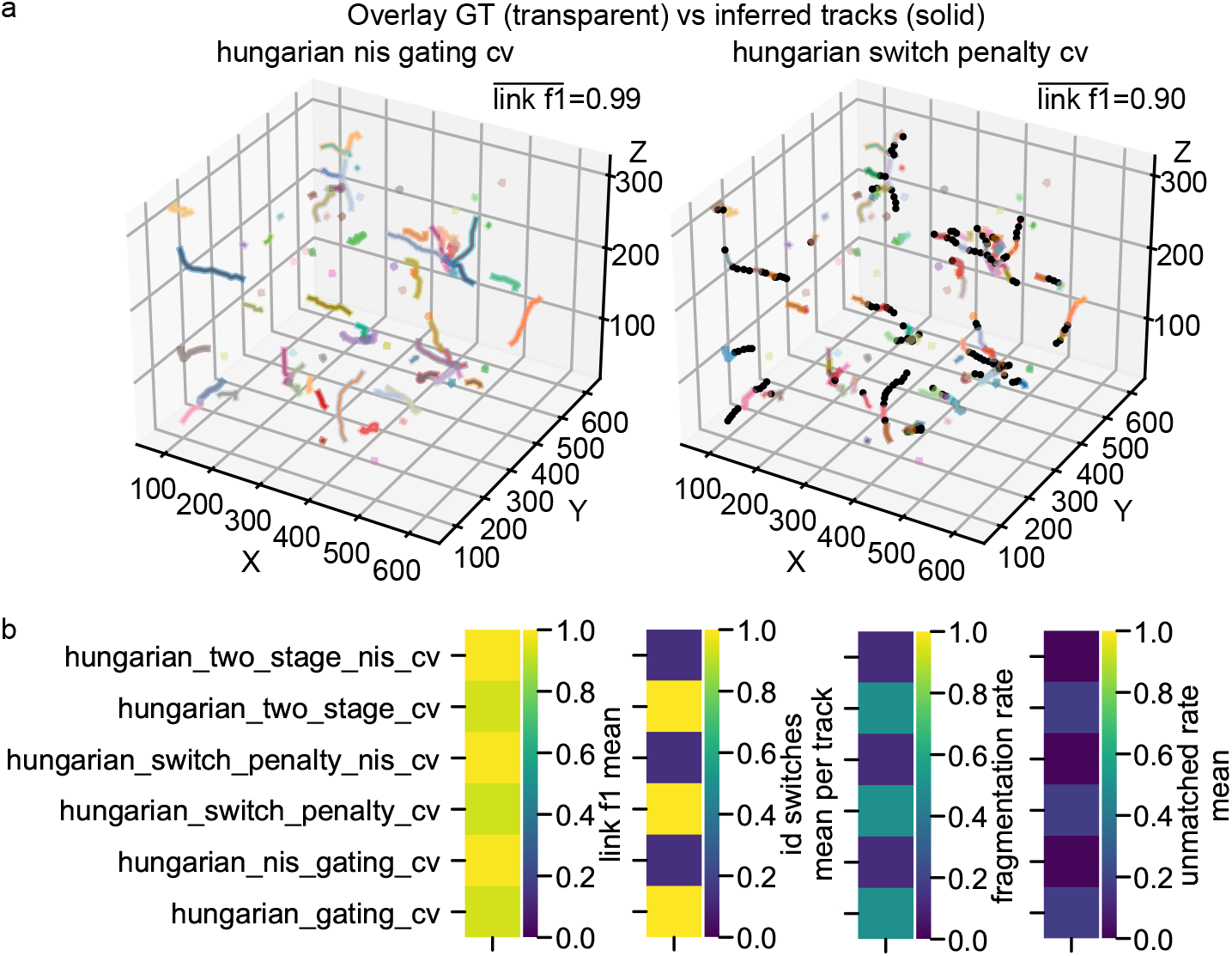
Association robustness under birth/death events and mixed stationary or migrating populations, without drift. Synthetic no-drift stress benchmark with heterogeneous displacement amplitudes, explicit appearance and disappearance events, and a mixed population containing both stationary and migrating objects. All methods belong to the Hungarian constant-velocity family and differ only in their gating strategy and optional penalties or staging. (a) Representative reconstructions overlaid with the transparent ground truth. Left: Hungarian constant-velocity association with innovation-based gating, illustrating near-perfect recovery with mean link-*F*_1_ = 0.99. Right: Hungarian constant-velocity association with coordinate-space gating and directional switch penalty, illustrating visibly degraded recovery with mean link-*F*_1_ = 0.90. Black points indicate detections that were not linked into consecutive pseudo-tracks. (b) Final method comparison after selecting, for each method, the parameter configuration with highest mean link-*F*_1_ across benchmark realizations. Shown are mean link-*F*_1_ (higher is better), mean identity switches per track (lower is better), fragmentation rate (lower is better), and mean unmatched rate (lower is better). Innovation-based variants clearly dominate the coordinate-space variants across all metrics. All values are averaged across the five synthetic realizations.

**Figure 8:**
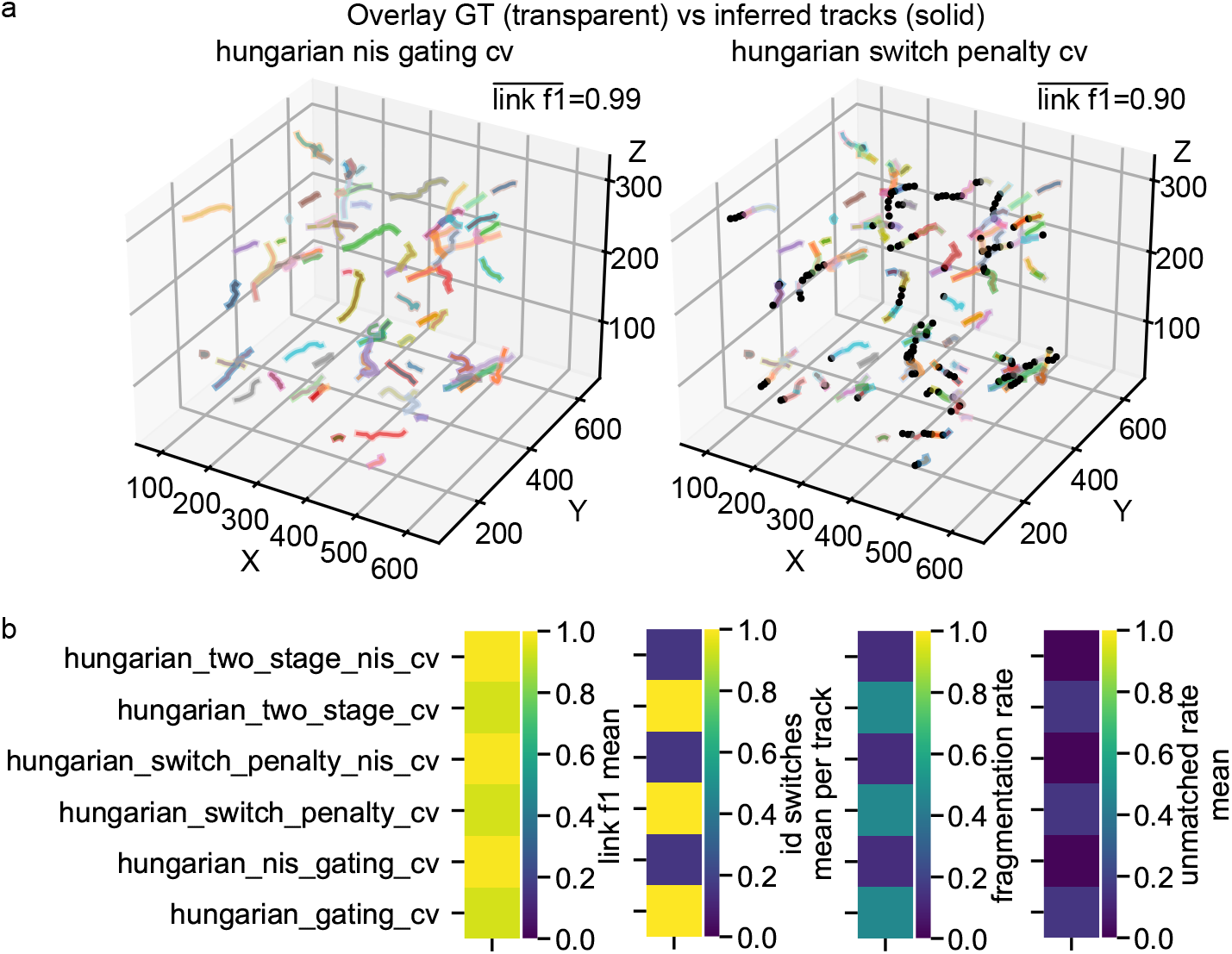
Association robustness under birth/death events, mixed motility, and additional drift contamination. Synthetic stress benchmark identical in spirit to Figure 7, but with added global drift consisting of smooth translation and rotation applied to the observed coordinates. This tests whether the stronger association variants remain stable when local linking must operate under residual geometric contamination. (a) Representative reconstructions overlaid with the transparent ground truth. Left: Hungarian constant-velocity association with innovation-based gating, again yielding near-perfect recovery with mean link-*F*_1_ = 0.99. Right: Hungarian constant-velocity association with coordinate-space gating and directional switch penalty, showing visibly more fragmentation and unmatched detections with mean link-*F*_1_ = 0.90. (b) Final method comparison after per-method parameter selection by mean link-*F*_1_. Shown are mean link-*F*_1_, mean identity switches per track, fragmentation rate, and mean unmatched rate. The ranking remains essentially unchanged relative to the no-drift birth/death benchmark: innovation-based variants remain clearly superior, indicating strong robustness to residual drift contamination. All values are averaged across the five synthetic realizations.

The coordinate-space variants remain substantially worse in the same benchmark. The plain coordinate-space Hungarian constant-velocity method reaches a mean link-*F*_1_ of 0.930, the switch-penalty version drops slightly further to 0.929, and the two-stage coordinate-space version is only marginally better at 0.930. All three are associated with about 1.62 identity switches per track and fragmentation rates around 0.50. This is a large performance gap, not a fine-tuning effect. Relative to the innovation-based group, the coordinate-space methods produce roughly an order-of-magnitude increase in identity switches and about a fourfold increase in fragmentation. Thus, once appearance and disappearance are allowed and some objects become effectively stationary, the main advantage of innovation-based gating remains intact.

Figure 8 shows that this ranking is preserved even after adding residual global drift on top of the birth/death and mixed-motility setting. Again, the three innovation-based variants are tied for best with mean link-*F*_1_ = 0.989, mean identity switches of 0.160 per track, fragmentation of 0.130, and unmatched rate near 0.012. The coordinate-space variants remain clustered around mean link-*F*_1_ ≈ 0.935, with identity switches around 1.45, fragmentation around 0.47, and unmatched rates near 0.157. The difference from the no-drift birth/death benchmark is therefore remarkably small. Adding drift causes only a mild quantitative degradation, and it affects both method families far less than the difference between gating in coordinate space and gating in normalized innovation space.

This stability under drift contamination is important because it links the present association study back to the broader pipeline. In practice, identity reconstruction may be performed on geometries that are not perfectly drift-free, or on data in which residual geometric contamination remains after correction. The present stress test indicates that the innovation-gated Hungarian constant-velocity family is robust to such residual contamination, at least within the synthetic regime considered here. In other words, the method does not require perfectly clean geometry in order to maintain high linking quality.

The density benchmark in Figure 9 provides the hardest stress test, because it directly increases local ambiguity while keeping the observation volume fixed. Here the number of tracks is scaled from the baseline density up to a twentyfold increase, with birth and death events present throughout. Three effects are immediately visible. First, all methods degrade with increasing density, as expected. Second, the degradation is much stronger for the coordinate-space variants than for the innovation-based variants. Third, despite this increasing ambiguity, the innovation-based methods remain clearly separated from the coordinate-space group across the full sweep.

**Figure 9:**
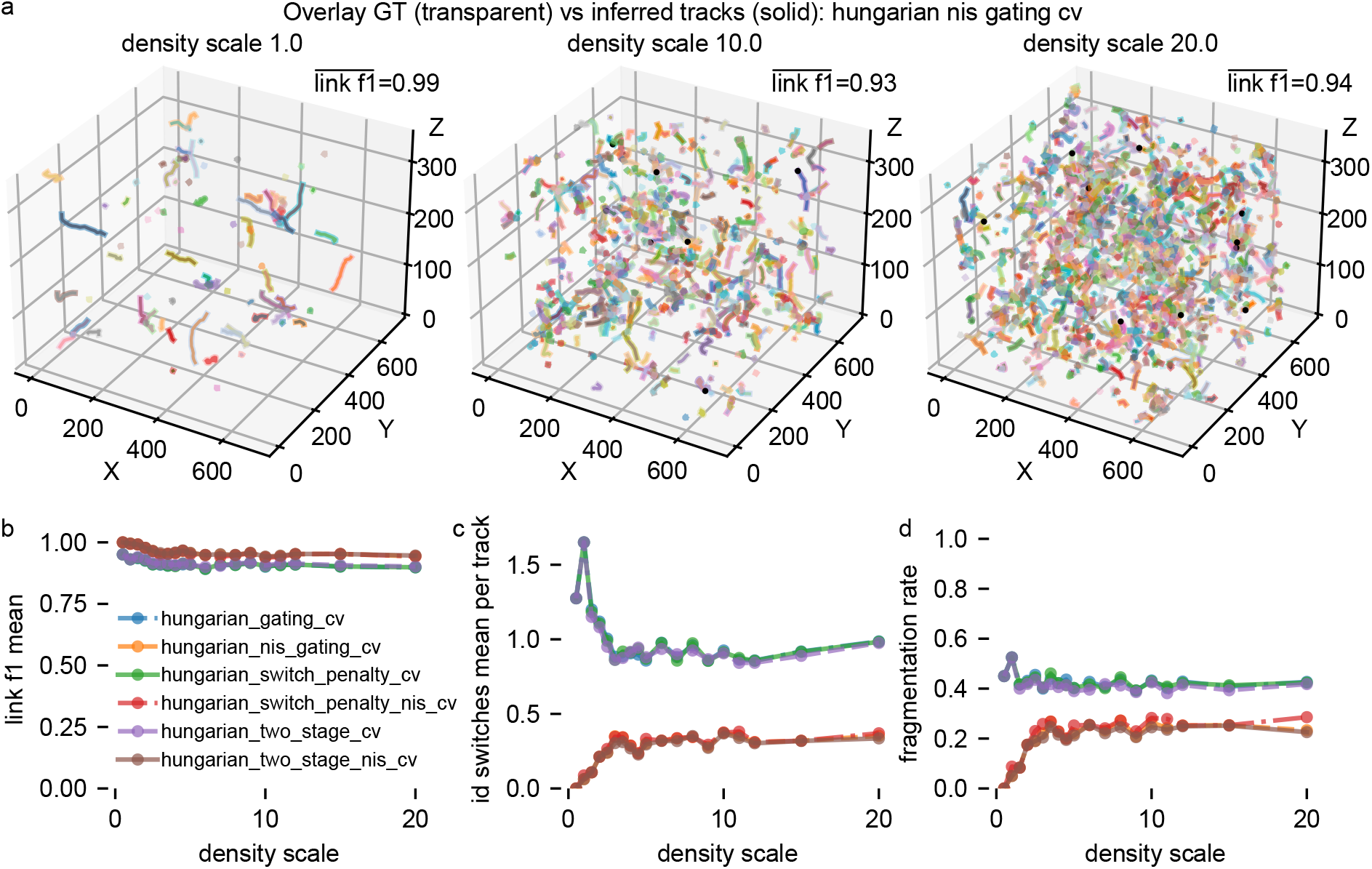
Density stress test for Hungarian constant-velocity association variants. Synthetic no-drift benchmark with birth/death events and mixed motility, in which the number of tracks is progressively increased at fixed volume to induce stronger local ambiguity and reduced nearest-neighbor separation. (a) Representative overlays of ground truth and inferred tracks for Hungarian constant-velocity association with innovation-based gating at three density scales. At baseline density (1.0), reconstruction remains almost perfect; at density scales 10.0 and 20.0, ambiguity increases substantially, but the global track structure remains largely preserved. Black points indicate unmatched detections. (b) Mean link-*F*_1_ as a function of density scale. Innovation-based variants remain consistently above the coordinate-space variants across the full sweep. (c) Mean identity switches per track as a function of density scale. (d) Fragmentation rate as a function of density scale. The density benchmark constitutes the strongest ambiguity stress test in the association study and shows that innovation-based gating degrades gracefully rather than failing abruptly under crowding. All values are averaged across the five synthetic realizations after selecting the best parameter configuration per method and density level by mean link-*F*_1_.

For mean link-*F*_1_, the innovation-based variants start near perfect at low density and remain around 0.94 even at the highest tested density scales. For example, Hungarian constant-velocity association with innovation gating decreases from 0.990 at density scale 1.0 to 0.937 at 10.0 and 0.941 at 20.0. The switch-penalty innovation variant and the two-stage innovation variant remain almost identical throughout, with differences only in the third decimal place. By contrast, the coordinate-space variants start already lower at the baseline density and then settle around 0.89 to 0.90 across the higher-density range. Thus the innovation-based group preserves an absolute link-*F*_1_ advantage of roughly four to five percentage points under heavy crowding.

The secondary diagnostics show the same separation. For identity switches per track, the coordinate-space group begins near 1.65 at baseline density and, after a transient decrease, remains close to 0.94 to 0.98 over the denser regimes. The innovation-based group remains far lower, typically between about 0.35 and 0.40 over the same range. Fragmentation follows the same pattern: the coordinate-space variants stabilize near 0.42 to 0.45, whereas the innovation-based variants stay near 0.24 to 0.28. The representative overlays in Figure 9a illustrate what this means qualitatively. At density scale 1.0, the innovation-gated reconstruction is almost indistinguishable from the ground truth. At density scales 10.0 and 20.0, local ambiguity becomes clearly visible and unmatched detections appear more frequently, but the global track structure still remains largely correct.

Two additional conclusions follow from the density sweep. First, the switch penalty and the two-stage strategy again provide little or no benefit once innovation-based gating is already present. The three innovation-based curves are nearly superimposed for link-*F*_1_, identity switches, and fragmentation across the entire density range. Second, the innovation-based family does not collapse abruptly as density increases. Instead, performance degrades gradually and remains in a regime that is still practically usable even under extreme crowding. This is exactly the type of behavior one would hope for in a robust local association rule.

Taken together, these compositional stress tests provide the strongest evidence in the association study. The innovation-based Hungarian constant-velocity variants remain highly reliable not only when the displacement scale is unknown, but also when the benchmark includes birth and death processes, mixtures of stationary and migrating objects, residual drift contamination, and severe density-induced ambiguity. The coordinate-space variants remain consistently inferior under all three stress conditions, and neither a directional switch penalty nor a two-stage assignment strategy is able to close that gap. The main determinant of robustness is therefore the use of normalized innovation-based gating itself. This identifies Hungarian constant-velocity association with innovation gating as the most reliable local association strategy across the full range of synthetic conditions considered here.

### Biological example workflows and end-to-end validation

The synthetic benchmarks above establish the ranking of association and drift-correction methods under controlled conditions in which the latent geometry and the imposed perturbations are known by construction. We next asked whether the same pipeline remains useful on biological time-lapse data, where segmentation is imperfect, intrinsic cell motion is heterogeneous, and the available trajectory references are necessarily proxy references derived from image processing rather than manually curated truth sets. As described in the Supplementary Methods, we therefore constructed two complementary biological examples: a sparse 3D+*t* carcinoma sequence (Kapoor, 2022) for which a known rigid drift could be imposed on a segmentation-derived trajectory reference, and a dense 2D+*t* DAPI sequence with strong native drift and no axial sampling (Radvansky et al., 2026), for which the association and correction methods had to operate on effectively planar tracks embedded in ℝ^3^ via *z* = 0.

#### Three-dimensional carcinoma example with imposed rigid drift

The carcinoma example (Kapoor, 2022) provides the closest biological analogue of the synthetic stress tests because it combines real cell trajectories with a known imposed rigid perturbation. As summarized in Figure 10a,b and detailed in the Supplementary Methods, we first segmented the raw low-SNR 3D+*t* volume sequence by volumetric Otsu-type thresholding with morphological regularization and then linked the segmented components over time by connected-component analysis (CCA) and centroid continuation. The resulting CCA trajectories served as a proxy ground truth for downstream evaluation. This step is important because it converts a raw fluorescence stack into a trajectory reference with persistent cell identities, which is required both for benchmarking point-cloud association (PCA) and for evaluating the combined PCA → Drift Correction pipeline. As in any segmentation-derived reference, however, this proxy ground truth is not error-free and may still contain missed detections, merges, splits, or identity inconsistencies.

**Figure 10:**
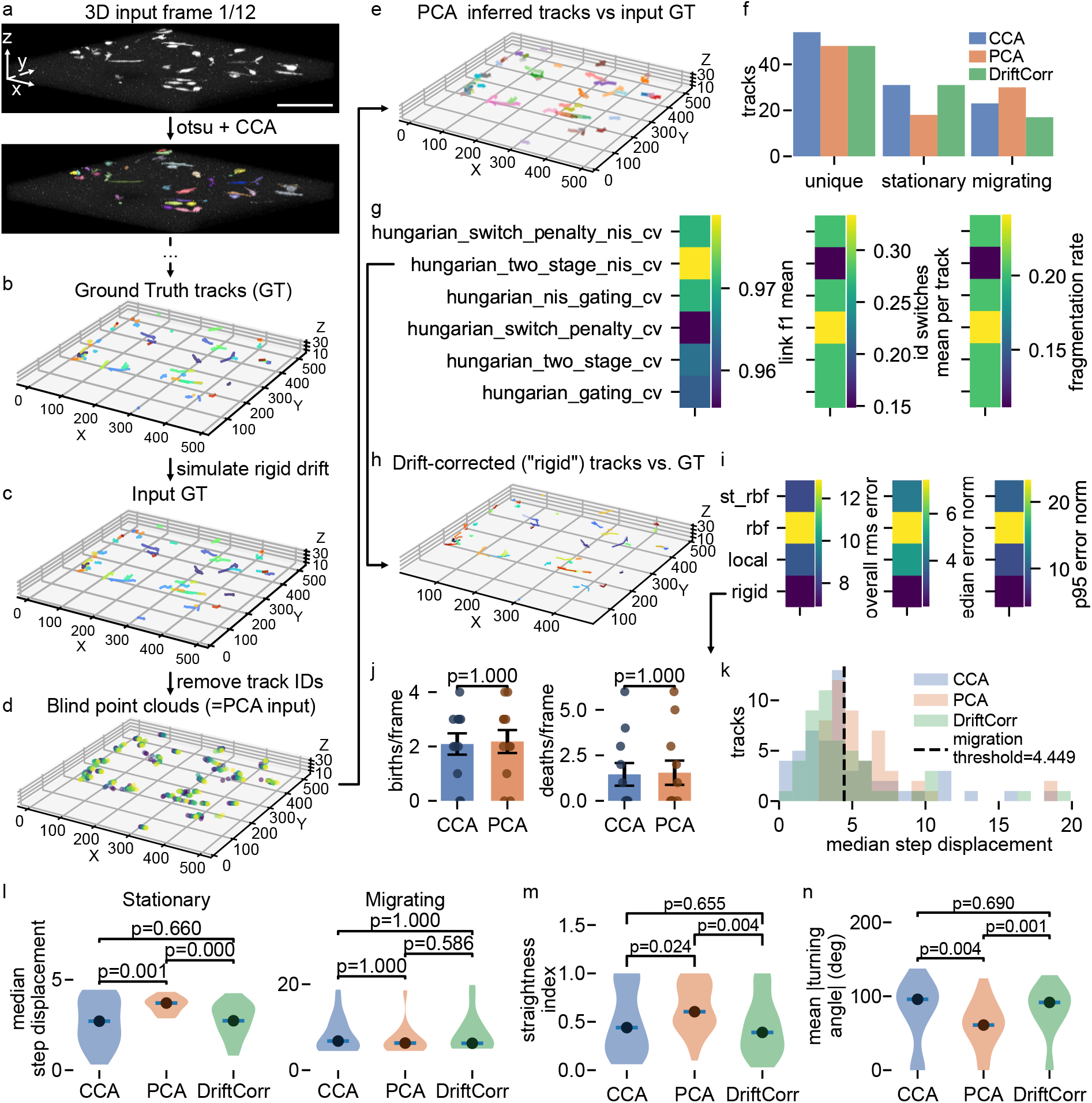
Biological 3D+*t* carcinoma example with imposed rigid drift. (a) Raw input volume at frame 1*/*12 (top; scalebar shows 100 *µ*m) and corresponding Otsu-plus-CCA proxy reference extraction (bottom). (b) CCA-inferred proxy ground-truth tracks (GT). (c) Rigidly drifted input GT obtained by adding a known global translation to the proxy GT. (d) Blind point clouds after removal of trajectory identities (input for the point cloud association, PCA). (e) Best PCA reconstruction (here: two-stage Hungarian constant-velocity strategy with innovation gating; thin, solid tracks) overlaid with the drifted input GT (thick, transparent tracks). (f) Unique, stationary, and migrating track counts for CCA, PCA, and drift correction (DriftCorr). (g) PCA quality metrics: mean link-*F*_1_, mean identity switches per track, and fragmentation rate. (h) Best drift-corrected solution (here: rigid; thin, solid tracks) overlaid with the undrifted GT (thick, transparent tracks). (i) GT-referenced drift-correction metrics: overall RMS error, median error norm, and *p*95 error norm. (j) Birth and death counts per frame for CCA and PCA. (k) Histogram of per-track median step displacement with the stationary–migrating threshold. (l) Median step displacement for stationary and migrating tracks. (m) Straightness index. (n) Mean absolute turning angle.

Because the raw carcinoma stack exhibited little intrinsic drift, we then imposed a known uniform translation on all proxy-GT trajectories to construct the observed input GT (Figure 10c). This choice was made deliberately to create a biologically realistic but geometrically controlled end-to-end stress test: the latent trajectories remain those of real cells, yet the observed coordinates are contaminated by a known rigid acquisition-like bias. After removing the trajectory identities, we obtained the blind point-cloud input for association (Figure 10d). Among the six HCV variants, the two-stage innovation-gated variant was best in this example, with mean link-*F*_1_ = 0.979, mean identity switches of 0.148 per track, and fragmentation rate 0.111 (Figure 10g). The qualitative overlay in Figure 10e is consistent with these values: the inferred pseudo-tracks recover the global identity structure well, although some CCA trajectories are fragmented into shorter inferred segments. This is precisely the failure mode expected from the synthetic association study, where ambiguity primarily manifests as fragmentation rather than grossly incorrect global reassignment.

The inferred trajectories from this best PCA method were then used as input to the drift-correction stage (DriftCorr). Among the four candidate correction models, the rigid stepwise model again performed best in GT-referenced evaluation, with overall RMS error 6.86 *µ*m, median error norm 2.02 *µ*m, and *p*95 error norm 4.18 *µ*m (Figure 10i). The overlay in Figure 10h shows that the corrected trajectories visually collapse back onto the undrifted proxy GT to a good approximation. At the same time, some mismatches remain visible because the input for the drift correction is the actual PCA output rather than the input GT itself. As described in the Supplementary Methods, GT-referenced drift-correction metrics were computed after deterministic one-to-one remapping between inferred and proxy-GT identities. Consequently, the reported RMS-type errors summarize the end-to-end PCA → DriftCorr pipeline rather than the geometric correction stage in isolation: fragmentation or merge errors introduced by PCA propagate directly into the final GT-referenced residuals.

The downstream track-count and motion summaries reinforce this interpretation. CCA yielded 54 unique proxy-reference tracks, of which 31 were classified as stationary and 23 as migrating using a Gaussian-mixture threshold on the per-track median step displacement (threshold 4.45 *µ*m; Figure 10k). PCA reduced the number of unique tracks to 48, decreased the stationary count to 18, and increased the migrating count to 30 (Figure 10f). This shift is exactly what one would expect after adding rigid drift to a mixed stationary-plus-migrating population: true stationary tracks are artificially displaced into the migrating regime, while PCA also loses some identity continuity by fragmenting or merging reference trajectories. After drift correction, the stationary count returned to 31, matching the CCA reference, whereas the migrating count decreased to 17. The most plausible interpretation is that rigid correction successfully removed the globally induced false migration component, but could not fully restore the original migrating count because the PCA stage had already reduced the total number of recoverable identities.

Several additional metrics support the same conclusion. First, birth and death counts per frame were unchanged between CCA and PCA (both Wilcoxon *p* = 1.000; Figure 10j), indicating that the association stage did not systematically invent or suppress track onsets or offsets. Instead, the main discrepancy lies in how detections are partitioned into continuous identities. Second, the median step displacement of stationary tracks increased from 2.64 *µ*m in the CCA reference to 3.63 *µ*m after PCA, but returned to 2.76 *µ*m after drift correction; the difference between CCA and DriftCorr was not significant (*p* = 0.660), whereas both contrasts involving PCA were significant (Figure 10l). By contrast, the migrating subset showed no significant change across groups, consistent with the idea that the rigid perturbation mainly inflates the apparent motion of otherwise weakly moving cells. Third, PCA made trajectories markedly straighter and less tortuous: the straightness index increased from 0.495 for CCA to 0.644 for PCA and then returned to 0.450 after correction, with no significant difference between CCA and DriftCorr (*p* = 0.655; Figure 10m). The complementary turning-angle measure behaved inversely, decreasing from 78.9° for CCA to 61.8° for PCA and recovering to 81.4° after correction, again with no significant difference between CCA and DriftCorr (*p* = 0.690; Figure 10n). Together, these shape descriptors show that the added rigid drift does not merely displace trajectories globally, but also makes them appear artificially straighter; the rigid correction step largely restores the original irregularity and directional variability of the biological motion pattern.

Overall, the carcinoma example therefore provides a realistic biological validation of the synthetic conclusions. Even when the latent motion is not generated by our simulation model but arises from real cells, the best HCV association variant still provides a sufficiently reliable identity backbone for downstream correction, and the rigid drift-correction model is able to recover the original motion statistics of the proxy reference with high fidelity under imposed global drift.

#### Two-dimensional DAPI example as a planar stress test

The synoviocyte DAPI example (Radvansky et al., 2026) addresses a different and in some ways harder question: can methods designed for 3D point clouds still behave sensibly on dense, long, effectively planar data with strong intrinsic drift? As described in the Supplementary Methods, we therefore embedded the 2D+*t* Cellpose detections (Stringer et al., 2021) into ℝ^3^ by fixing *z* = 0 and linked them over time by a post hoc Hungarian linker to obtain a proxy trajectory reference (denoted HL). Figure 11a,b introduces the dataset and the reference-building step. The analysis itself was always performed on the full field of view; the zoomed panel in Figure 11b is shown only to make the segmentation and identity labeling visually interpretable. The resulting HL trajectories already look highly irregular and tortuous (Figure 11c), which is expected because the raw sequence contains both genuine cell migration and substantial frame-to-frame geometric contamination.

**Figure 11:**
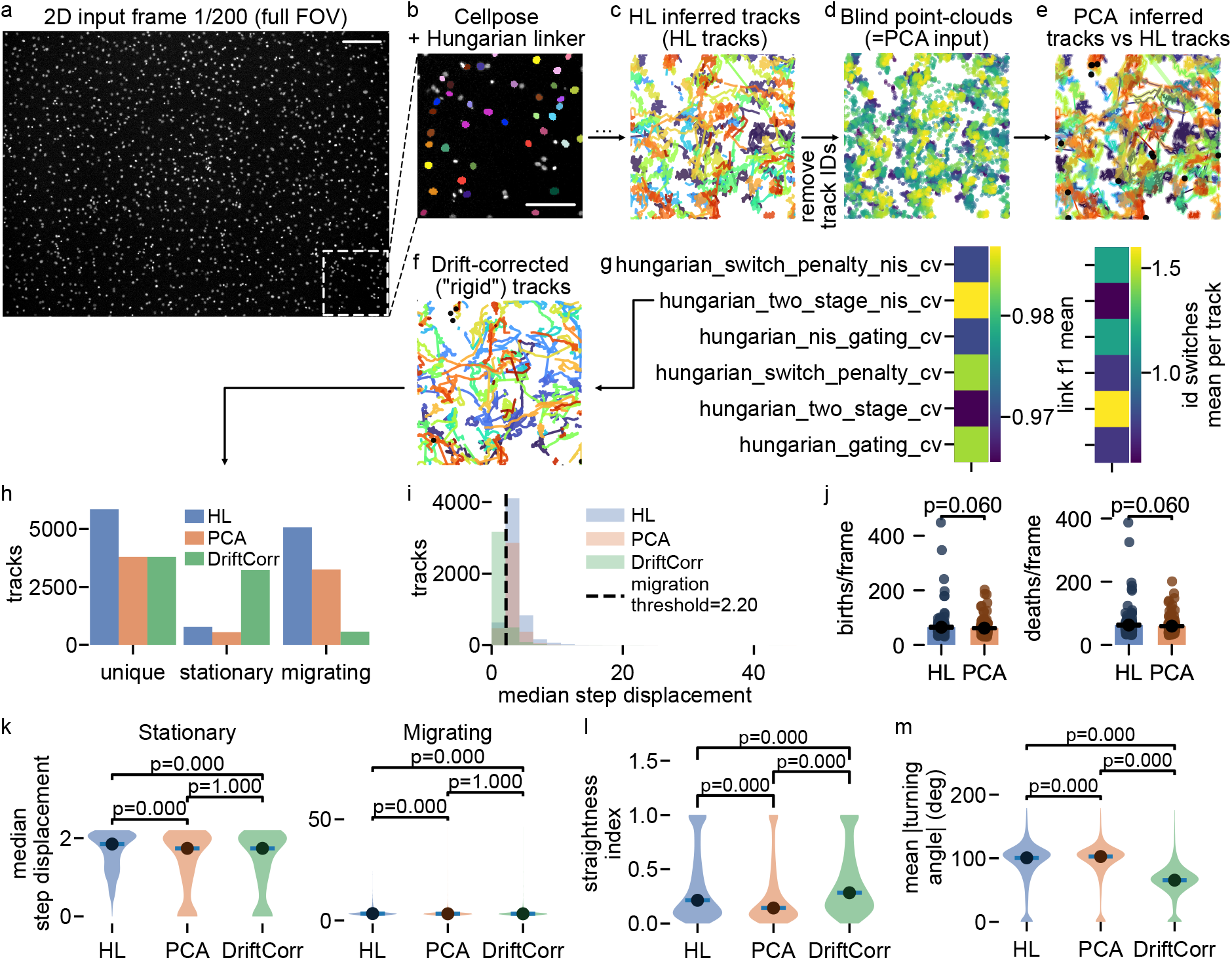
Biological 2D+*t* DAPI example with intrinsic drift and planar embedding. (a) Full-field overview of frame 1*/*200 with the zoom region indicated (scalebar shows 200 *µ*m). (b) Zoom-in showing Cellpose segmentation and Hungarian-linked proxy reference (scalebar shows 100 *µ*m). (c) HL-inferred tracks over the full sequence. (d) Blind point clouds after removal of trajectory identities. (e) Best PCA reconstruction (here again: two-stage Hungarian constant-velocity strategy with innovation gating; thin, solid tracks) overlaid with the HL tracks (thick, transparent tracks). (f) Drift-corrected trajectories for the `rigid`model. (g) PCA quality metrics: mean link-*F*_1_ and mean identity switches per track. (h) Unique, stationary, and migrating track counts for HL, PCA, and DriftCorr. (i) Histogram of per-track median step displacement with the manually chosen stationary–migrating threshold. (j) Birth and death counts per frame for HL and PCA. (k) Median step displacement for stationary and migrating tracks. (l) Straightness index. (m) Mean absolute turning angle.

**Figure 12:**
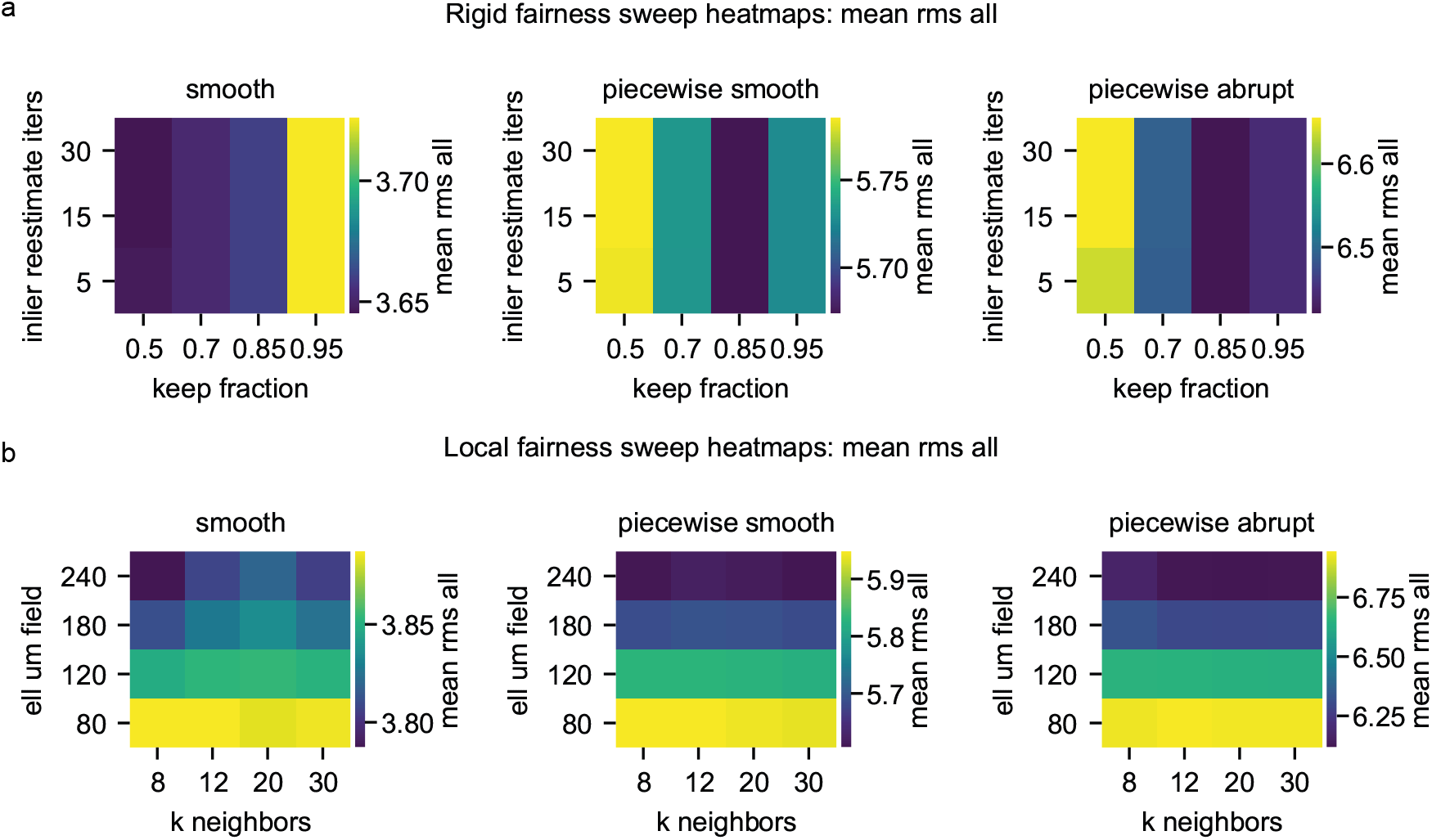
Fairness sweeps for rigid and local baseline models. Parameter-sensitivity analysis for the two baseline correction models “rigid” and “local” evaluated in the main benchmark suite. Results are aggregated across the three spatially varying drift regimes (smooth, piecewise smooth, piecewise abrupt). Colors indicate the mean overall corrected-versus-ground-truth RMS error across synthetic realizations, reported in *µ*m. (a) Rigid stepwise model sweep. The trimmed robust estimation parameters in Eq. (8) are varied by sweeping the inlier keep fraction *ρ* ∈ {0.5, 0.7, 0.85, 0.95} and the number of inlier re-estimation iterations {5, 15, 30} . Performance varies only weakly across this range, with the best region near *ρ* ≈ 0.85 and minimal dependence on iteration count. (b) Local community drift model sweep. The neighborhood size used for graph construction and the spatial blending scale of the drift field are varied by sweeping *k* ∈ {8, 12, 20, 30} neighbors and field length scale ℓ_field_ ∈ {80, 120, 180, 240} *µ*m in Eq. (19). Larger blending scales generally yield lower RMS, indicating that stronger spatial smoothing improves robustness in this benchmark. Overall performance differences across the explored parameter ranges remain modest, confirming that the baseline-suite results are not driven by a narrow choice of hyperparameters.

**Figure 13:**
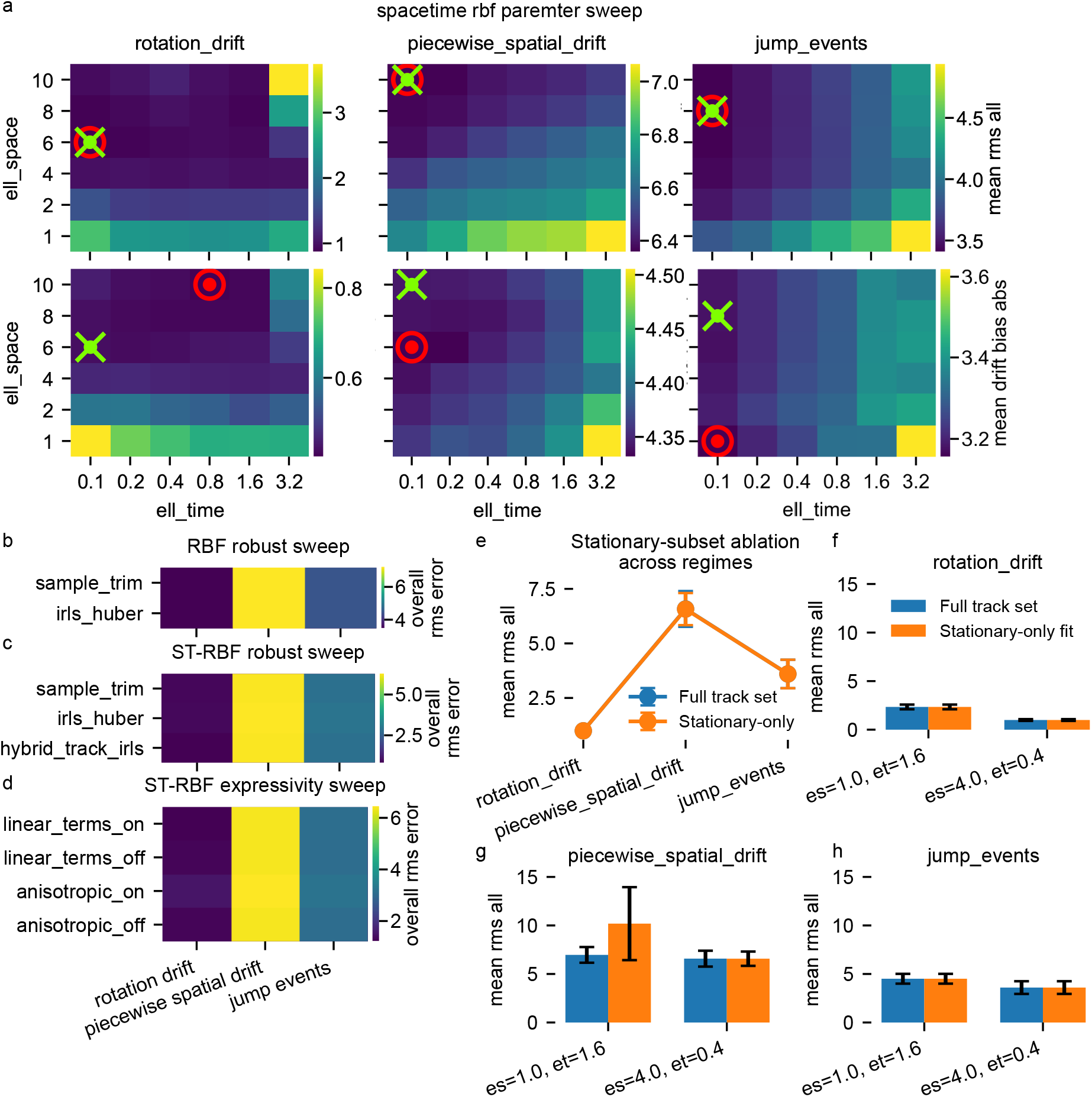
RBF-family robustness analyses and hyperparameter sweeps across baseline stress regimes. (a) Space-time RBF lengthscale sweep over ℓ_space_ ∈ {1, 2, 4, 6, 8, 10} and ℓ_*t*_∈ { 0.1, 0.2, 0.4, 0.8, 1.6, 3.2}, aggregated separately for rotation drift, piecewise spatial drift, and jump events. The top row shows the mean global RMS error 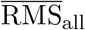 and the bottom row the mean absolute drift-bias 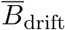, each averaged across benchmark realizations. Red markers indicate the minimum of the plotted metric. Green crosses indicate the best compromise between RMS and drift bias according to the combined score defined in Eq. (1). (b) Per-timepoint RBF robust sweep comparing trimmed ridge fitting and IRLS with Huber weights, summarized by mean overall RMS across regimes. (c) Space-time RBF robust sweep comparing trimmed ridge fitting, IRLS with Huber weights, and track-level trimming combined with IRLS refitting. (d) Space-time RBF expressivity sweep comparing isotropic versus anisotropic spatial kernels and linear-term augmentation on versus off, again summarized by mean overall RMS. (e) Stationary-subset ablation for the practical space-time RBF candidate (ℓ _space, *t*_) = (4.0, 0.4), comparing training on the full track set versus training only on the heuristically selected stationary subset. Error bars indicate standard deviation across benchmark realizations. Here, *overall RMS* denotes the dataset-level RMS over all corrected points, and *mean RMS* denotes the average of that quantity across realizations. All values are reported in *µ*m.

**Figure 14:**
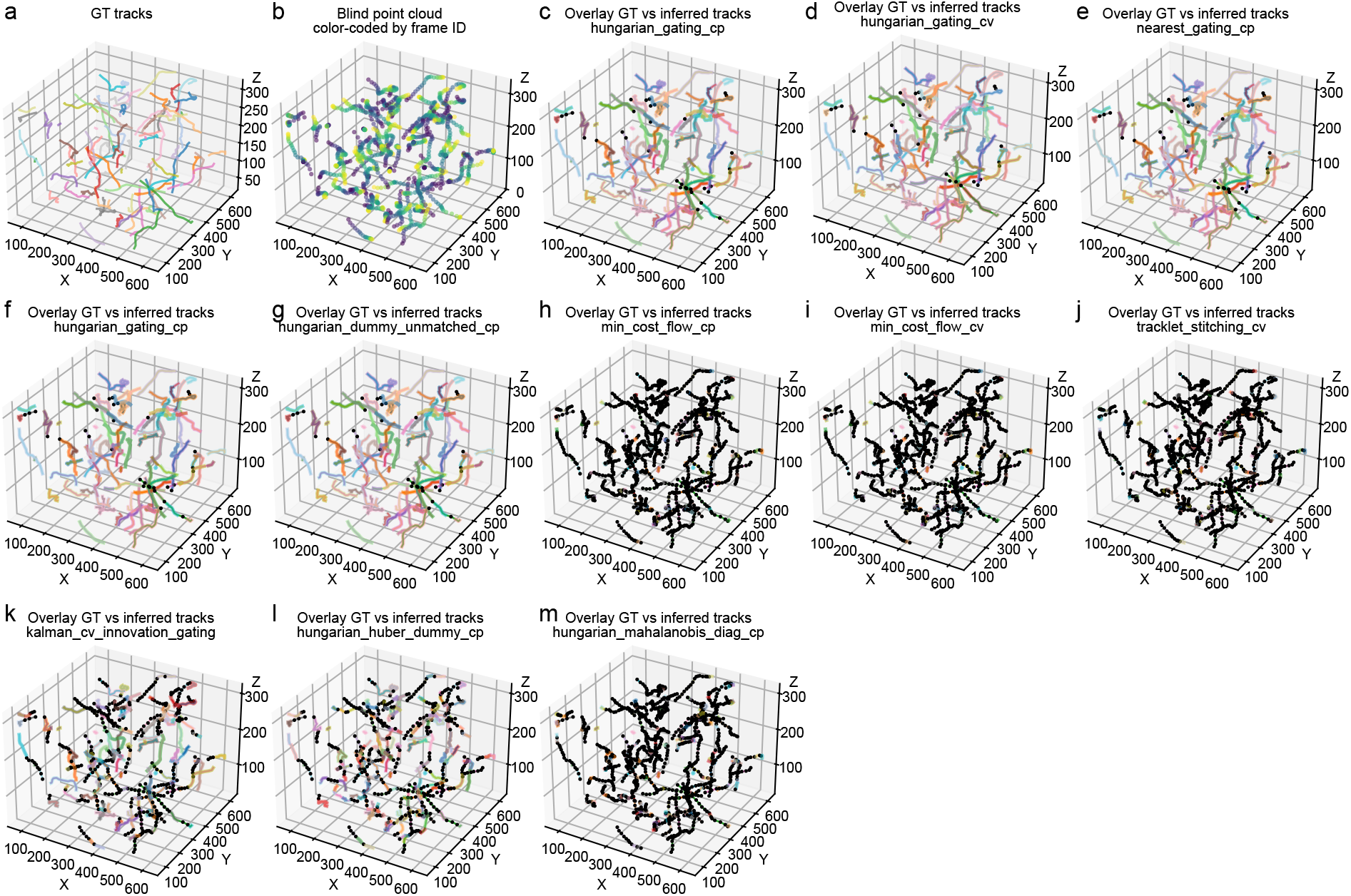
Qualitative reconstruction examples for all association methods in the baseline benchmark. Visualization of ground-truth trajectories and reconstructed tracks for a representative synthetic dataset from the no-drift displacement benchmark. (a) Ground-truth trajectories used to generate the synthetic observations. (b) Corresponding blind point cloud obtained by removing track identities and pooling detections across frames, color-coded by frame index. (c–m) Overlay of ground-truth trajectories (transparent, slightly thicker lines) with reconstructed tracks (solid colored lines) for each tested association method. Black points indicate detections that were not assigned to a consecutive track continuation by the respective method. Differences in reconstruction quality are visible as fragmented trajectories, incorrect linkages, or large numbers of unmatched detections. Panels correspond to the following methods: Hungarian assignment with coordinate-space gating (CP) and with constant-velocity refinement (CV), nearest-neighbor gating, explicit dummy-unmatched handling, minimum-cost-flow linking, tracklet-first stitching, Kalman constant-velocity association with innovation gating, Huberized assignment cost with dummy unmatched modeling, and Mahalanobis-scaled assignment cost. These examples complement the quantitative comparison in Figure 5 by illustrating typical reconstruction failure modes across the tested association families.

After removing the HL identities, we again benchmarked the six HCV association variants on the resulting blind point clouds. The same method that was strongest in the carcinoma example, the two-stage Hungarian constant-velocity strategy with innovation gating, was also best here, with mean link-*F*_1_ = 0.987, mean identity switches of 0.569 per track, and fragmentation rate 0.341 (Figure 11g). The overlay in Figure 11e shows that the inferred tracks recover the gross continuation structure of the HL reference well, despite the far greater density and temporal length of this dataset. At the same time, the comparison between PCA and HL should not be overinterpreted as a comparison against absolute truth. HL itself is only a proxy reference derived from Cellpose segmentation and a post hoc linker, and thus may already contain fragmented or imperfect continuations. In this setting, agreement between PCA and HL should be interpreted as consistency with a strong proxy reference rather than with an externally validated cell-tracking ground truth.

For the drift-correction stage, no drift-free GT was available. We therefore selected a representative correction model from the no-GT summaries and carried forward the rigid stepwise model, because it produced the strongest reduction of short-lag MSD (median MSD(*τ*_1_) from 14.93 to 3.39 *µ*m^2^) together with the largest increase in the near-stationary fraction (from 0.066 to 0.153). The corrected trajectories in Figure 11f are qualitatively much smoother and less zig-zag-like than the corresponding PCA tracks, already suggesting that a substantial part of the apparent motion in the raw linked trajectories was due to common geometric contamination rather than to true cell migration.

The track-count summaries in Figure 11h make this point quantitatively clearer. HL yielded 5849 unique trajectories, PCA yielded 3796, and DriftCorr of course retained the same number of identities as its PCA input. The drop from HL to PCA indicates that the HCV association stage consolidates many short or interrupted HL track fragments into longer continuations. Given the known limitations of the HL proxy reference, this reduction should not be read simply as identity loss; it may equally reflect recovery of continuity across brief segmentation failures. The stationary/migrating partition changes even more strongly. Using a fixed manual threshold of 2.2 *µ*m on the per-track median step displacement (Figure 11i), HL classified 774 tracks as stationary and 5075 as migrating. PCA preserved roughly the same overall balance but with fewer total tracks (547 stationary, 3249 migrating). After drift correction, however, the stationary count increased dramatically to 3225, while the migrating count dropped to 571. The natural interpretation is that a large fraction of the trajectories that initially appeared migratory were in fact dominated by common drift and should not be interpreted as independently migrating cells once that shared geometric component is removed.

The birth/death analysis again indicates that PCA mainly redistributes identity continuity rather than changing event counts. Birth and death rates per frame were not significantly different between HL and PCA (Wilcoxon *p* = 0.060 for both; Figure 11j). Biologically, these events are expected because cells can enter or leave the field of view across the image boundaries over a long experiment. Technically, additional apparent births and deaths can arise when segmentation intermittently fails for a nucleus and the corresponding trajectory is temporarily interrupted. The lack of a significant shift between HL and PCA therefore suggests that the association step largely preserves the global event budget even though it reassigns many individual continuations.

The motion summaries show that PCA and DriftCorr affect different aspects of the trajectory geometry. For stationary-class tracks, the median step displacement decreased from 1.67 *µ*m in HL to 1.39 *µ*m after PCA and further to 1.11 *µ*m after drift correction; all pairwise differences were significant (Figure 11k). Thus, even before explicit drift correction, the HCV association stage already suppresses some high-frequency local irregularity relative to the HL reference. For the migrating class, the median step displacement changed more subtly in absolute terms (3.79 *µ*m for HL, 3.57 *µ*m for PCA, 3.91 *µ*m for DriftCorr), but again all pairwise contrasts were significant. This pattern can be explained by the fact that after correction, many weakly moving trajectories are reclassified out of the migrating set, so that the remaining migrating subset becomes smaller but more strongly motile on average. In such a dense dataset, statistical significance is expected even for modest absolute differences because the number of trajectories is very large.

The path-shape metrics provide the clearest evidence that drift correction alters the geometry in the expected direction. PCA alone reduced the straightness index from 0.354 in HL to 0.266, whereas DriftCorr increased it to 0.379; all pairwise differences were highly significant (all *p <* 10^−18^; Figure 11l). In parallel, the mean absolute turning angle changed from 89.7° in HL to 92.4° after PCA and then dropped sharply to 62.0° after drift correction, again with all pairwise differences highly significant (Figure 11m). Together with the visual impression in Figure 11f, these two metrics show that the correction stage makes trajectories globally smoother and more persistent in direction, exactly as one would expect if an important part of the original zig-zag structure were caused by frame-to-frame geometric drift rather than by true stochastic cell motility.

This DAPI example is therefore a genuine planar stress test for the full workflow. The data are dense, long, intrinsically drift-contaminated, and strictly two-dimensional, yet the same HCV association family and the same downstream correction framework still produce coherent and interpretable results when applied to coordinates embedded in ℝ^3^ with *z* = 0. The association stage again provides a strong identity backbone, and the drift-correction stage reveals that much of the apparent migration in the uncorrected proxy tracks is better explained as shared geometric contamination.

#### Summary of biological examples

Taken together, the two biological examples extend the synthetic benchmark conclusions in complementary directions. The carcinoma example shows that, on real volumetric cell trajectories with an imposed known perturbation, the PCA → DriftCorr pipeline can recover both geometry and motion statistics close to a segmentation-derived trajectory reference. The DAPI example shows that the same pipeline remains usable in a much harder regime with intrinsic drift, dense populations, and effectively planar data. In both cases, the innovation-gated HCV association family provides the most reliable identity reconstruction backbone, and the rigid correction model either remains quantitatively best (carcinoma) or produces the clearest geometric regularization of the inferred tracks (DAPI). Thus the synthetic benchmarks do not merely optimize methods for artificial data; they correctly anticipate which method families transfer most robustly to biological time-lapse imaging.

## Discussion

Structural time-lapse microscopy frequently leads to a coupled inference problem that is often only partially addressed in practice: object identities must be reconstructed from unlabeled point clouds, and systematic geometric drift must be corrected after the image data have already been reduced to centroid coordinates. These two tasks are not independent. Drift estimation requires sufficiently consistent trajectories, because identity errors bias any fitted drift field. Conversely, identity reconstruction depends on the observed geometry and is therefore perturbed by drift itself. The main motivation of this study was to make this coupling explicit and to benchmark both components within a common point-cloud framework.

A first central result is that, on the drift-correction side, greater model flexibility did not automatically improve performance. Across both the baseline spatially varying regimes and the more structured non-rigid stress tests, the rigid stepwise model was the most consistently reliable method, and the local community model was the strongest alternative when region-wise spatial heterogeneity became more important. The space-time RBF model often formed an intermediate tier, while the per-timepoint RBF model and the mixture-of-experts extensions were generally less robust. This is an important negative result. It suggests that, in the present point-cloud setting, the limiting factor is not simply the representational capacity of the drift model. Instead, robustness of estimation under mixed stationary and migrating populations appears at least as important as nominal expressivity. A model that can represent highly structured drift fields is not necessarily the model that best recovers latent trajectories if the fitting procedure becomes unstable or if genuine biological motion is partially absorbed into the inferred correction.

This also explains why the rigid model remained surprisingly competitive even in regimes that explicitly violated the assumption of a single global transform. Many synthetic drift fields, and also the biological examples, still contained a substantial coherent component that could be captured by a robust rigid estimate. Once that dominant shared component is removed, the remaining spatial heterogeneity often contributes less to the final trajectory error than the nominal complexity of the ground-truth field might suggest. The local community model benefited in those regimes where coherent region-wise differences were large enough to justify local flexibility without requiring a fully nonparametric global fit. By contrast, the more expressive RBF-family models paid a robustness penalty that was not compensated by commensurate gains in geometric fidelity. In practical terms, the results argue against the common intuition that increasingly flexible drift fields should automatically dominate once spatial nonuniformity is present.

An important practical implication of these findings is that drift correction can be meaningfully shifted from the image domain to the point-cloud domain. In many experimental workflows, motion correction is performed directly on image stacks prior to segmentation, typically using rigid or affine registration against a reference template. However, such approaches are not always applicable or reliable. Suitable templates may be unavailable or difficult to construct in long recordings with evolving structure, and global transformations may fail to capture complex motion patterns such as in-frame motion artifacts, abrupt shifts, rotational drift, or tissue deformation. In these situations, image-based registration may partially correct global misalignment while simultaneously distorting or blurring biologically meaningful structures and temporal dynamics.

The present framework addresses these limitations by operating directly on segmentation-derived point clouds. This enables drift correction to be performed post hoc, without requiring access to or modification of the original image data. In particular, it allows correction in regimes where image-based methods are ill-posed or introduce undesirable side effects. By decoupling geometric correction from image intensity information, the approach provides a complementary strategy that remains robust under heterogeneous and partially nonrigid motion, while preserving the discrete structure of tracked objects. This is especially relevant in settings where segmentation is reliable but image-level registration is challenging or ambiguous.

A second central result concerns identity reconstruction. Here the benchmarks yielded an even sharper conclusion. Within the Hungarian constant-velocity family, the decisive factor was not the addition of auxiliary heuristics such as directional switch penalties or staged assignment alone, but the choice of gating space. Once displacement scales became heterogeneous, uncertain, or embedded in more realistic settings with births, deaths, density stress, and residual drift contamination, innovation-based gating consistently outperformed coordinate-space gating. The innovation-gated Hungarian constant-velocity family remained close to perfect in the unknown-displacement benchmark and degraded gracefully under increasingly difficult stress conditions, whereas coordinate-space variants showed markedly higher fragmentation and identity-switch rates. The main methodological lesson is therefore that candidate correspondences should be judged relative to an inferred uncertainty scale rather than by a fixed, or only weakly adaptive, Euclidean gate.

This finding is broader than the specific synthetic scenarios used here. Real time-lapse data rarely admit a single globally valid displacement scale. Motion statistics can vary across cells, over time, with depth, and across experimental conditions. Under such heterogeneity, a dimensionless innovation criterion is methodologically preferable to a gate defined purely in coordinate space. At the same time, the results also show that once innovation-based gating is in place, further elaborated variants often produce only marginal gains. Directional switch penalties and two-stage assignment were useful robustness variants and in some settings slightly improved the final ranking, but they did not change the primary conclusion. The essential ingredient was the innovation-normalized gating itself.

The biological examples support these conclusions, but they also clarify how they should be interpreted outside a fully synthetic setting. In the carcinoma example, a known rigid perturbation was imposed on segmentation-derived volumetric trajectories. This provided a realistic end-to-end test in which the latent motion was biological but the geometric contamination was controlled. Here, the best association method was again an innovation-based Hungarian constant-velocity variant, and the best correction model was again the rigid stepwise model. The correction stage restored stationary versus migrating structure and improved trajectory-shape statistics such as straightness and turning-angle distributions relative to the reference. This is important because it shows that the benchmark rankings are not merely artifacts of an artificial simulator, but remain meaningful in trajectories extracted from real imaging data.

The DAPI example answered a complementary question: whether the same framework remains meaningful in dense, long, intrinsically drift-contaminated, effectively planar data when supplied with coordinates embedded into ℝ^3^ via *z* = 0. Again, the same association family produced the strongest identity backbone, and the rigid correction model yielded the clearest geometric regularization of trajectories. More importantly, this example illustrates a point that is easy to miss in purely synthetic studies. A substantial fraction of what appears as cell migration in long uncorrected recordings may in fact reflect common geometric contamination. The pronounced increase in stationary-class trajectories after correction, together with reduced turning angles and increased straightness, is consistent with exactly that interpretation. The biological examples therefore do more than reproduce the synthetic rankings. They show why those rankings matter for downstream biological interpretation.

A recurrent theme in both biological examples is the role of proxy references. In the synthetic benchmarks, drift and identity errors can be separated by construction because ground-truth trajectories and ground-truth drift fields are independently known. In the biological examples, by contrast, the available references are segmentation-derived or linker-derived proxy references and may themselves contain fragmentation, merges, missed detections, or identity inconsistencies. Accordingly, a reported RMS-type disagreement in the biological analyses reflects the quality of the entire PCA → DriftCorr pipeline relative to that proxy reference, not an isolated property of the correction model. This is not a weakness of the present framework, but a realistic description of the practical problem. In real applications, drift correction is rarely applied to perfect trajectories. It is applied to whatever trajectory set an upstream tracking or association stage can provide.

The biological results therefore support two complementary interpretations. First, they validate the pipeline as a practical workflow: the strongest association family produces trajectories that are sufficiently consistent to support useful drift correction, and the strongest correction model improves downstream trajectory geometry in biologically interpretable ways. Second, they caution against reading all disagreement with a proxy reference as direct algorithmic failure. In dense or low signal-to-noise settings, disagreement may reflect imperfections in the reference as well as in the evaluated method. This is one reason why the combination of controlled synthetic benchmarks and biological examples is methodologically stronger than relying on either source of evidence alone.

Taken together, the results support a concrete default workflow for point-cloud-based structural time-lapse analysis. When stable identities are not available, identity reconstruction should be performed with a Hungarian constant-velocity scheme using innovation-based gating. When the downstream goal is geometric correction of centroid trajectories, the rigid stepwise model should be considered the first-line correction strategy, with the local community model as the main alternative when region-wise spatial heterogeneity is expected to be substantial. More flexible RBF-based models may still be useful as exploratory tools or in very specific regimes, but the present benchmarks do not support them as default choices. This recommendation is attractive precisely because it is conservative: it achieves strong performance with comparatively simple models, fewer hyperparameters, and greater interpretability than the more expressive alternatives.

The conclusions should, however, be interpreted within the scope of the benchmark design. The synthetic regimes span a broad range of drift structures, motion heterogeneity, birth and death processes, and density-induced ambiguity, but they still derive from a specific stochastic simulator and cannot represent the full diversity of biological time-lapse data. Strongly deformable tissue motion, structured anisotropic noise, severe segmentation dropout, or systematic morphology changes may introduce coupled failure modes not fully captured here. Likewise, the biological examples do not provide manually curated absolute cell-tracking truth sets. They are sufficient to test whether the pipeline behaves coherently and improves trajectory geometry in biologically interpretable ways, but they do not allow a definitive decomposition of total error into reference error, association error, and correction error.

A second limitation is architectural rather than empirical. Identity reconstruction and drift correction were treated sequentially rather than jointly. This separation keeps both components transparent and benchmarkable in their own right, but it also leaves open the possibility that tighter coupling could further improve performance in especially difficult regimes. Similarly, the present framework operates deliberately on geometric summaries alone. This is a practical advantage when only centroid exports are available, but it also means that appearance features, segmentation confidence, and image-level context remain unused. These are natural directions for future work, especially if the goal is to move from a robust, interpretable baseline workflow toward a more tightly integrated probabilistic inference framework.

In summary, the main contribution of this study is not a single new drift model or a single new association heuristic in isolation, but a unified point-cloud perspective on structural time-lapse analysis in which identity reconstruction and drift correction are treated as coupled, benchmarkable inference problems. The synthetic results identify a clear practical ranking: robust rigid or local correction on the geometric side, and innovation-gated Hungarian constant-velocity association on the identity side. The biological examples show that these conclusions remain meaningful beyond synthetic data and can materially improve the interpretation of real cell trajectories. Overall, the results argue for a methodological middle ground between unconstrained image registration and naive point matching: a geometry-aware, uncertainty-aware point-cloud workflow that remains simple enough to be interpretable, yet strong enough to handle realistic drift and tracking ambiguity.

## Materials and Methods

To formalize geometric drift correction and identity reconstruction from temporally sampled three-dimensional point sets, we introduce a general observation model linking the measured coordinates to latent object positions. We then specify parameterized models for systematic geometric drift and formulate identity reconstruction, in the absence of explicit track labels, as an assignment problem under uncertainty.

Consider a sequence of volumetric observations acquired at discrete time points *t* ∈ {1, …, *T*} . After segmentation, each time point is represented by a finite set of three-dimensional object coordinates. We denote the observed point set at time *t* by

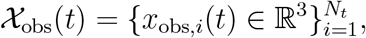

where *N*_*t*_ may vary over time due to object appearances and disappearances.

Each observed coordinate is assumed to arise from an underlying latent object position that is affected by systematic geometric distortion and measurement noise. We model this relationship as

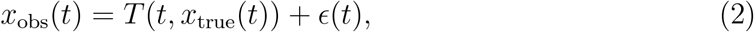

where *x*_true_(*t*) ∈ ℝ^3^ denotes the latent true position of an object at time *t, T* (*t*, ·) is a time-dependent transformation describing systematic geometric bias, and *E*(*t*) represents measurement noise and residual modeling error. The transformation *T* (*t*, · ) may encode globally rigid motion, spatially varying drift, or smooth deformation fields.

To make the role of drift explicit, we parameterize the transformation *T* (*t*, · ) as an identity mapping plus an additive displacement field. In its simplest form, this yields an additive drift model

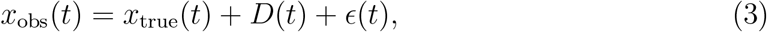

where *D*(*t*) ∈ ℝ^3^ denotes a global, time-dependent drift vector shared by all objects at time *t*. This model corresponds to a purely translational drift component and serves as a baseline within the drift model hierarchy considered here.

In volumetric recordings, however, residual misalignment can vary across the field of view or along depth. To capture such spatially non-uniform effects, we generalize the drift term to a displacement field *D*(*t, x*) : {1, …, *T* } *×* ℝ^3^ → ℝ^3^ and write

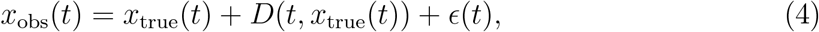

that is, *T* (*t, x*) = *x* + *D*(*t, x*). Different parameterizations of *D*(*t*) and *D*(*t, x*) define the drift models analyzed in this study.

### Trajectory representation

If object identities are known, the temporal evolution of object *i* can be represented as a trajectory 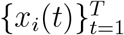. In practice, however, explicit identity labels may be unavailable, and the number of detected objects *N*_*t*_ can change across time. Trajectory reconstruction must therefore be inferred from the observed point sets.

### Coupling between drift and identity

Two structural dependencies follow from the model Eq. (2). First, association procedures commonly rely on geometric proximity between successive frames, for example through distances of the form

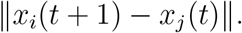

When observations are affected by drift, the apparent displacement between two time points satisfies

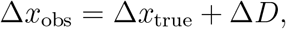

so that systematic drift perturbs inter-point distances and apparent velocities. This modification of the geometric structure directly influences assignment decisions and propagates into trajectory-level metrics such as displacement errors, track consistency, and identity switch rates.

Second, the estimation of *T* (*t*, ·) from trajectories presupposes trajectory continuity. In the models considered below, drift parameters are inferred from collections of temporally linked positions 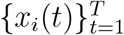. Identity inconsistencies such as ID switches or track fragmentation therefore bias the estimated drift field, since the inferred motion no longer corresponds to a single physical object.

Drift estimation and identity reconstruction thus form a coupled inference problem. Reliable estimation of the transformation *T* (*t*, · ) requires consistent trajectories, while reliable trajectory reconstruction benefits from geometrically corrected observations. The following sections specify parameterized drift models under the assumption of known trajectories and subsequently formulate identity reconstruction as an assignment problem when explicit track labels are unavailable.

### Geometric drift field models

The drift models considered here estimate, for each pair of consecutive time points (*t*_0_, *t*_1_), a geometric transformation that maps coordinates at time *t*_0_ to coordinates at time *t*_1_ using matched object trajectories. Let *i* index objects with known identity across time (known tracks). For each consecutive pair (*t*_0_, *t*_1_) we form the matched coordinate sets

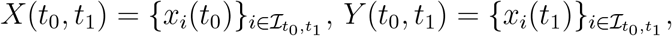

where 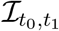 is the index set of objects present at both times. Each drift model estimates parameters 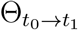 of a transformation 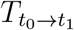 by minimizing a robust discrepancy between 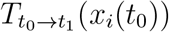 and *x*_*i*_(*t*_1_) over 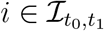. The per step transformations are then composed over time and inverted to map all observations into a common reference frame.

#### Rigid stepwise model

##### Rigid transformation between consecutive frames

In the global rigid model, the transformation between consecutive time points is assumed to be a rigid motion composed of a rotation 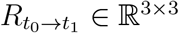 and a translation 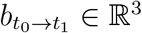, such that

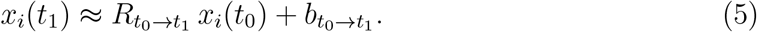

Given matched point sets 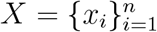 and 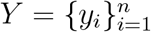 with *x*_*i*_ = *x*_*i*_(*t*_0_) and *y*_*i*_ = *x*_*i*_(*t*_1_), the rigid parameters are estimated by the Kabsch method (Kabsch, 1976; Kabsch, 1978).

Defining centered coordinates 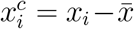 and 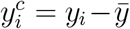 with 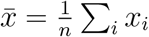 and 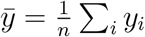, we form the cross covariance

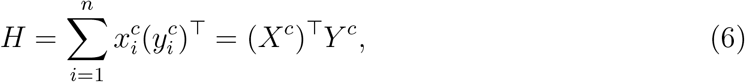

compute an SVD *H* = *U* Σ*V* ^*T*^, and set

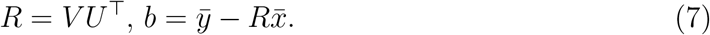

If det(*R*) *<* 0, the sign of the last singular vector is flipped to enforce det(*R*) = 1.

##### Trimmed robust estimation

To reduce sensitivity to outliers among the matched pairs, we use a trimmed iterative re-estimation procedure. Starting from an initial index set *S* containing all *n* pairs, we alternate between (i) estimating (*R, b*) from the current subset and (ii) retaining only the *k* pairs with smallest residual, where *k* = max(6, ⌊ *ρn*⌋ ) and *ρ* ∈ (0, 1] is a user specified keep fraction. For a current estimate (*R, b*), residuals are computed as

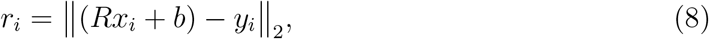

and the inlier subset is updated to the indices of the *k* smallest residuals. The procedure is repeated for a fixed number of re-estimation iterations or until the inlier set no longer changes. The final output consists of (*R, b*) and an inlier mask.

##### Minimum match requirement

A step transform is estimated only if the number of matched pairs *n* exceeds a minimum threshold *n*_min_. If *n < n*_min_, the step is marked as unavailable. This prevents unstable estimates in sparse regimes.

##### Composition and mapping to a reference frame

Let (*R*_*a*→*b*_, *b*_*a*→*b*_) denote the estimated rigid transformation from time *a* to time *b*. Rigid transforms compose as

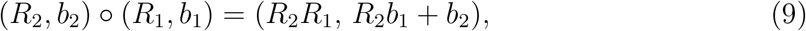

and invert as

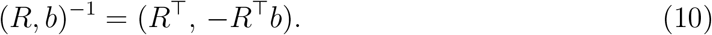

Choosing a reference time *t*_ref_, we build the cumulative forward transform from *t*_ref_ to each time *t* by composing the available per step transforms along the temporal chain. The correction transform that maps coordinates at time *t* into the reference frame is then the inverse of this cumulative forward transform. The corrected coordinates are computed as

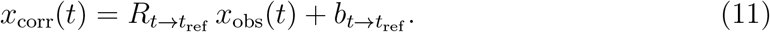

#### Local community based drift model with spatially smooth mixing

The rigid model estimates a single parametric transform between consecutive time points. As an alternative, we consider a local drift model that is formulated directly in terms of track displacement traces relative to a reference time and that allows different spatial neighborhoods to exhibit distinct drift dynamics. The method proceeds in three stages: (i) extract per track displacement time series, (ii) build a spatially local graph whose edge weights encode motion similarity and spatial proximity, and (iii) infer a small number of track communities and estimate one drift trace per community, which is then blended into a continuous drift field in space.

##### Displacements relative to a reference time

Let *t*_ref_ ∈ {1, …, *T*} be a chosen reference time and let *x*_*i*_(*t*) ∈ ℝ^3^ denote the observed coordinate of track *i* at time *t* when it exists. For tracks that are present at *t*_ref_ and at least one additional time point, we define the displacement trace relative to the reference as

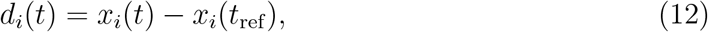

interpreted componentwise in *x, y, z*. In addition, we associate each track with a representative spatial anchor position *p*_*i*_ ∈ ℝ^3^, defined as the temporal mean of its observed positions,

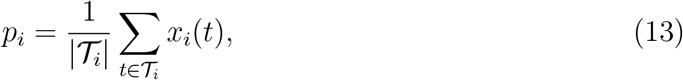

where *T*_*i*_ denotes the set of times at which track *i* exists.

##### Motion similarity and spatially local graph construction

We construct an undirected weighted graph *G* = (*V, E*) whose nodes correspond to tracks. Two nodes *i* and *j* are connected only if the corresponding anchors *p*_*i*_ and *p*_*j*_ are spatial neighbors, defined either by *k* nearest neighbors or by a fixed radius criterion.

To weight candidate edges, we combine motion similarity with spatial proximity. Motion similarity is computed from the displacement traces *d*_*i*_(*t*) and *d*_*j*_(*t*) using axiswise Pearson correlation. Let 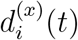 denote the *x* component of *d*_*i*_(*t*), and analogously for *y* and *z*. We define

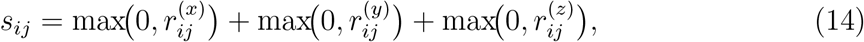

where 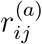 is the Pearson correlation coefficient between the component traces 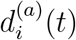 and 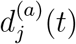 computed over time points at which both traces are finite. If fewer than three shared time points are available, or if either trace is effectively constant, the corresponding correlation is set to zero. By construction, *s*_*ij*_ ∈ [0, 3] and negative correlations are discarded.

Spatial proximity is encoded by a Gaussian kernel with length scale *ℓ*_graph_ *>* 0,

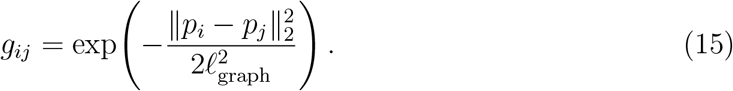

The final edge weight is then defined as

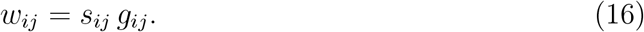

Edges with *w*_*ij*_ below a user defined threshold *w*_min_ are discarded to sparsify the graph and suppress weakly supported connections.

##### Community detection and community drift traces

Given the weighted graph, we partition the nodes into communities 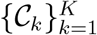 using a modularity based community detection algorithm on *G* (Louvain (Blondel et al., 2008) if available, otherwise greedy modularity (Clauset, Newman and Moore, 2004)). Each community defines a set of tracks that are both spatially close and exhibit similar displacement dynamics.

For each community *C*_*k*_, we define its spatial center as

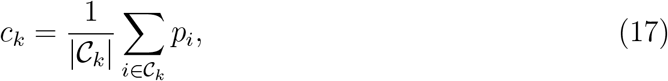

and estimate a community drift trace *d*_*k*_(*t*) ∈ ℝ^3^ as a robust componentwise median across member displacement traces,

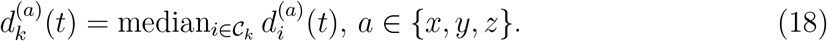

This median is computed using only finite values at each time point. The collection 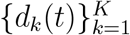 provides a discrete set of drift traces anchored at centers {*c*_*k*_}.

##### Spatial blending into a continuous drift field

To obtain a spatially continuous drift predictor, we blend the community drift traces with Gaussian weights. For a spatial position *p* ∈ ℝ^3^, define unnormalized weights

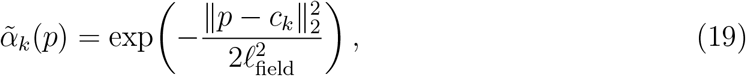

with blending length scale *ℓ*_field_ *>* 0, and normalize them as

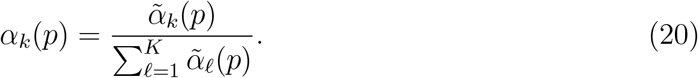

The predicted drift field at position *p* and time *t* is then

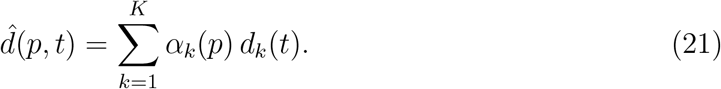

If *ℓ*_field_ ≤ 0, the predictor reduces to a spatial average over communities.

##### Drift correction

For each observation *x*_obs_(*t*), we evaluate the drift predictor at an associated position *p*, and subtract it to obtain corrected coordinates,

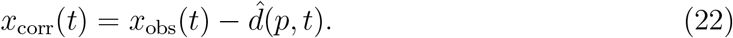

The position *p* used for prediction can be chosen as either the instantaneous coordinate *x*_obs_(*t*) or the track anchor *p*_*i*_ associated with the observation, with the latter yielding time independent spatial weights per track.

##### Model parameters

The local model is controlled by the graph neighborhood definition (either *k* nearest neighbors or a radius threshold), the graph length scale *ℓ*_graph_, the edge sparsification threshold *w*_min_, and the field blending scale *ℓ*_field_. The role of these parameters is conceptually separated: *ℓ*_graph_ affects which motion similarities are considered local when constructing the graph, whereas *ℓ*_field_ controls the spatial smoothness of the final drift field 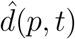.

#### Smooth drift fields via Gaussian radial basis functions (RBF)

The local community model constructs a drift field by first discretizing the volume into motion coherent communities and then blending their median drift traces in space. As a complementary approach, we directly parameterize the drift as a smooth displacement field using Gaussian radial basis functions (RBFs) (Broomhead and Lowe, 1988). The goal is to estimate, for each non reference time point, a function 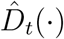 such that subtracting 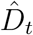 maps observations at time *t* into the reference coordinate system at time *t*_ref_ .

##### Reference pairing and drift targets

Fix a reference time *t*_ref_ ∈ {1, …, *T*} . For any time *t* & *t*_ref_, we form correspondences using tracks that exist at both *t* and *t*_ref_ . For each matched track instance we denote the observed coordinates by *x*_*i*_(*t*) and *x*_*i*_(*t*_ref_ ). The per point target drift vector is defined as

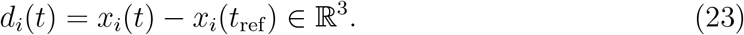

Given a set of *n*_*t*_ such correspondences at time *t*, we stack input positions into 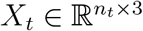 and target drifts into 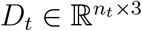.

A key implementation detail is the choice of spatial input coordinates used to evaluate the field. We denote the input position for correspondence *i* at time *t* by *ξ*_*i*_(*t*) ∈ ℝ^3^. Depending on the chosen mode, *ξ*_*i*_(*t*) is taken as the instantaneous observed coordinate *x*_*i*_(*t*) (row position mode), a time averaged track anchor, or the track coordinate at the reference time. In all cases, the regression target remains *d*_*i*_(*t*) as defined in Eq. (23).

##### Gaussian RBF parametrization

Let 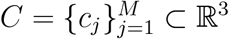 be a set of RBF centers and let *σ >* 0 be a kernel width. The Gaussian kernel matrix between inputs *X*_*t*_ and centers *C* is

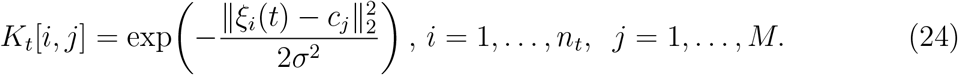

We model the drift field at time *t* as a linear combination of these kernels,

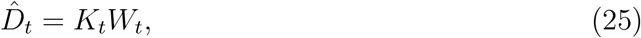

where *W*_*t*_ ∈ ℝ^*M ×*3^ contains the RBF weights for the three displacement components.

##### Kernel augmentation

To allow the model to represent a global translation even if center coverage is sparse or uneven, we augment the kernel with a constant bias column.

Denote by 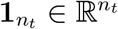 the all ones vector. We define the augmented design matrix

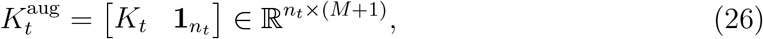

and the corresponding augmented weights 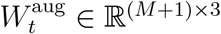 such that

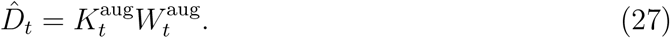

Optionally, additional linear terms in space and time can be appended to 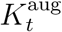, yielding basis functions *ξ*_*x*_, *ξ*_*y*_, *ξ*_*z*_ and *t* alongside the constant, in order to increase representational flexibility in regimes where the drift contains a substantial low order component.

##### Per time point ridge regression fit

For a fixed time *t* & *t*_ref_, the RBF weights are estimated by ridge regression (Hoerl and Kennard, 1970). With regularization strength *λ >* 0, the objective is

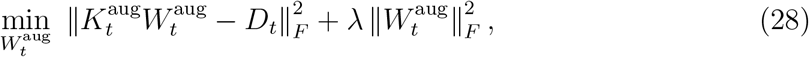

where ‖ · ‖_*F*_ is the Frobenius norm. The corresponding normal equations are

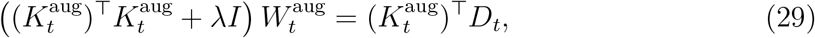

with *I* the identity matrix of size (*M* + 1) *×* (*M* + 1). In practice we also allow a distinct regularization for the bias column, *λ*_bias_ ≥ 0, by using a diagonal regularizer with the last entry set to *λ*_bias_.

##### Robust variants via trimming or iteratively reweighted least squares (IRLS)

To reduce sensitivity to mismatches or outlier correspondences, we use robust variants of Eq. (28). In the trimmed variant, we iterate between (i) fitting weights by ridge regression on a current inlier set and (ii) recomputing residuals

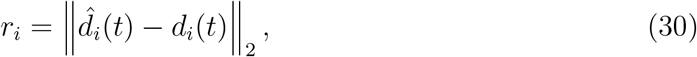

followed by retaining the fraction *ρ* ∈ (0, 1) of correspondences with smallest *r*_*i*_. The regression in the next iteration is then restricted to this reduced inlier set.

Alternatively, we use *iteratively reweighted least squares* (IRLS) (Holland and Welsch, 1977), a classical robust regression scheme. In this formulation, each correspondence is assigned a weight *w*_*i*_ ∈ [0, 1] derived from its residual norm *r*_*i*_. For a Huber-type weighting rule (Huber, 1964), we define a scale parameter *δ >* 0 and set

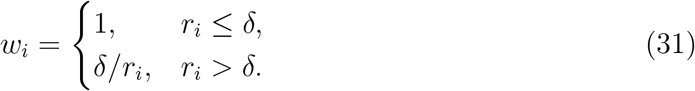

The weighted objective becomes

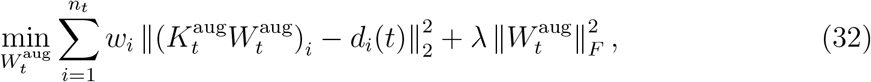

which is solved by ridge regression on the row-wise weighted design matrix and targets. The weights are recomputed from updated residuals and the procedure is iterated until convergence or until a maximum number of iterations is reached.

A second robust option operates at the track level rather than at the correspondence level. In this mode, one first fits fields using an initial track set, then computes a per-track residual summary across time points, for example the median of *r*_*i*_ over all timepoints where the track is present, and finally retains only the fraction of tracks with smallest residual summary before refitting the model.

##### Two stage baseline plus residual field

A simple decomposition used in our experiments is a two stage scheme separating global translation from residual spatial structure. First, the global translation component is estimated as the componentwise median drift across correspondences,

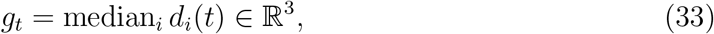

and removed to yield residual targets *d*^res^(*t*) = *d*_*i*_(*t*) − *g*_*t*_. Second, the RBF model is fit to the residual targets. The predicted drift then becomes

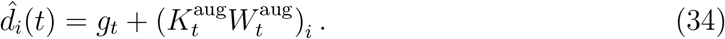

This improves numerical stability when the dominant drift component is global and the residual field is comparatively small.

##### Temporal regularization across time points

The per time point fits above treat each time independently. To encourage temporal consistency, we additionally consider a coupled formulation in which the RBF centers and kernel width are shared across time and the weights *W* ^aug^(*t*) evolve smoothly. Let *t*_1_ *<* … *< t*_*S*_ denote the set of times (excluding *t*_ref_ ) for which sufficient correspondences exist. We solve

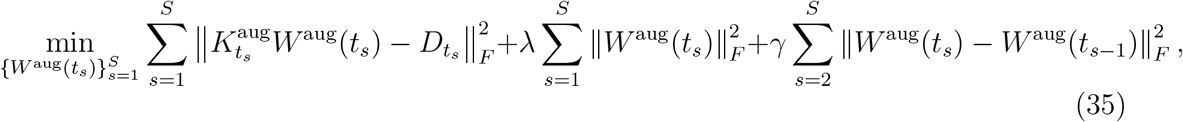

with temporal coupling strength *γ* ≥ 0. The resulting normal equations form a block tridiagonal linear system in the unknown weight matrices, which can be solved efficiently by forward elimination and backward substitution.

##### Applying the correction

After fitting 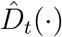, corrected coordinates at time *t* are obtained by subtracting the predicted drift:

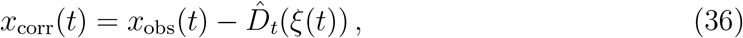

where *ξ*(*t*) denotes the chosen spatial input coordinate for field evaluation.

When the row position mode is used, the input coordinate depends on the corrected coordinate itself. In that case we apply a small fixed point iteration: initialize *ξ*^(0)^(*t*) = *x*_obs_(*t*) and update

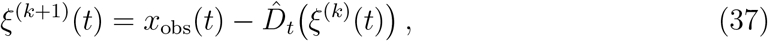

until the median update magnitude falls below a tolerance or a maximum number of iterations is reached.

##### Model parameters and hyperparameters

The RBF drift model is controlled by a small set of parameters that determine (i) representational capacity, (ii) spatial smoothness, (iii) numerical stability, and (iv) robustness to mismatched correspondences.

##### Centers *C* **and the number of centers** *M*

The field uses *M* Gaussian basis functions centered at 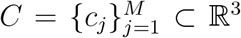; increasing *M* increases flexibility but raises variance and conditioning risk. In our implementation, centers are selected by one of three strategies: (i) *reference sampling*, where *C* is chosen by sampling from the point cloud at *t*_ref_ ; (ii) *pooled sampling*, where *C* is sampled from pooled observations across multiple time points; (iii) *pooled k-means*, where *C* is obtained by a k-means clustering of pooled observations. The pooled modes can reduce center sparsity when the reference frame contains limited spatial support.

##### Kernel width *σ*

The Gaussian kernel width *σ* in Eq. (24) controls spatial smoothness. Small *σ* yields highly localized influence of centers and can fit sharp spatial variation, but it increases the risk of overfitting and ill conditioning of (*K*^aug^)^*T*^*K*^aug^. Large *σ* enforces smooth fields that behave closer to a low frequency deformation. When *σ* is not fixed explicitly, we set it by a data driven heuristic based on center spacing, namely the median nearest neighbor distance among centers.

##### Ridge regularization *λ* and bias regularization *λ*_bias_

The ridge strength *λ* in Eq. (28) stabilizes the regression by penalizing large weights and is therefore critical for controlling conditioning and extrapolation artifacts. Optionally, the constant bias column in Eq. (26) can be regularized separately with *λ*_bias_, allowing a practically unpenalized global translation component while still regularizing the spatial weights.

##### Temporal coupling *γ*

In the temporally regularized formulation Eq. (35), the coupling parameter *γ* controls smoothness of the weight trajectories *W* ^aug^(*t*_*s*_). Setting *γ* = 0 reduces the model to independent per time point ridge fits. Larger *γ* enforces gradual temporal variation and reduces frame to frame noise in the inferred drift, at the cost of potentially smoothing true discontinuities such as stage jumps.

##### Robustness controls

Robust fitting can be enabled either at the correspondence level or at the track level.

At the correspondence level, we use a trimming fraction *ρ* ∈ (0, 1) that retains the *ρn*_*t*_ smallest residual correspondences per fit iteration, and a maximum number of iterations for convergence. Alternatively, an IRLS scheme replaces trimming by residual dependent weights derived from a Huber type rule, which reduces the influence of large residual correspondences without discarding them entirely.

At the track level, we iterate between fitting fields on a current set of tracks and computing trackwise residual summaries across time, retaining only the fraction with smallest residual summary before refitting. This mode is designed to suppress globally inconsistent tracks that would otherwise contaminate the drift estimate across many time points.

##### Two stage decomposition

The optional two stage scheme is controlled by a boolean switch. When enabled, the global translation component *g*_*t*_ is estimated by Eq. (33) and removed before fitting the RBF residual. This typically improves stability when translation dominates and the spatially varying component is small.

##### Choice of spatial input coordinates

The evaluation coordinate *ξ*(*t*) in Eq. (36) is chosen by a categorical parameter. In the row position mode, *ξ*(*t*) = *x*_obs_(*t*) and the correction becomes implicit, requiring the fixed point iteration in Eq. (37). In the track anchor and reference position modes, *ξ*(*t*) is fixed per track, which yields an explicit correction in a single pass.

#### Joint space-time RBF drift model

The space-time RBF model represents drift as a smooth vector field over a four-dimensional input consisting of spatial coordinates and time. For each correspondence between a track position at time *t* and its reference position at *t*_ref_, we define the drift target

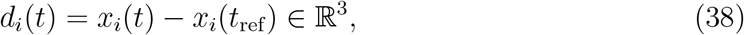

where *x*_*i*_(*t*) ∈ ℝ^3^ denotes the observed track position.

##### Spatial input modes

The model can be driven by different spatial input definitions to balance stability and expressivity. In *row position* mode, the instantaneous position is used as input, *s*_*i*_(*t*) = *x*_*i*_(*t*). In *track anchor* mode, the input is a track-specific anchor, for example the temporal mean position *s*_*i*_(*t*) = ⟨*x*_*i*_(*τ* ) ⟩ _*τ*_, which is constant in *t* for each track. In *reference position* mode, the input is the reference-time position *s*_*i*_(*t*) = *x*_*i*_(*t*_ref_ ) whenever available, with a fallback to the track anchor otherwise. These three modes affect the geometry on which smoothness is imposed, and thereby the tradeoff between locality and numerical stability.

##### Input normalization and 4D kernel

We normalize space per axis by a robust scale estimate. Let *c* ∈ ℝ^3^ be the componentwise median of all spatial inputs and *s* ℝ^3^ a robust per-axis scale, such as a median absolute deviation based estimate. The normalized spatial coordinates are

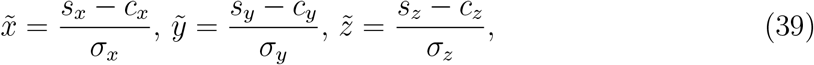

where (*s*_*x*_, *s*_*y*_, *s*_*z*_) = *s*_*i*_(*t*) denotes the chosen spatial input for the sample and (*σ*_*x*_, *σ*_*y*_, *σ*_*z*_) are the robust scales. Time is normalized to [0, 1] via

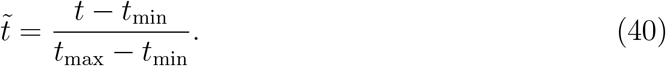

The resulting 4D input is 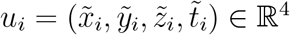.

Given *M* centers *c*_*m*_ ∈ ℝ^4^ in this normalized space, we define an anisotropic Gaussian kernel

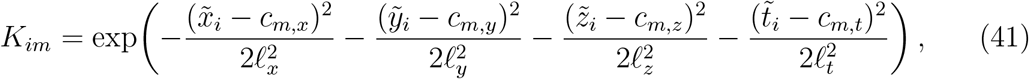

with spatial lengthscales *ℓ*_*x*_, *ℓ*_*y*_, *ℓ*_*z*_ and temporal lengthscale *ℓ*_*t*_ measured in normalized units. In the common isotropic spatial setting we set *ℓ*_*x*_ = *ℓ*_*y*_ = *ℓ*_*z*_ = *ℓ*_space_.

##### Augmented design and ridge objective

To represent global translation and, optionally, linear trends, we augment the kernel with additional columns. The basic augmentation appends a bias term,

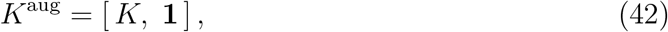

where **1** is the all-ones column. Optionally, we further append linear terms in 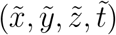 and interactions 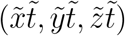 to capture affine drift components and space-dependent temporal trends.

Let 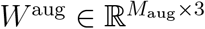 denote the weight matrix for the augmented design. We fit the model by ridge regression,

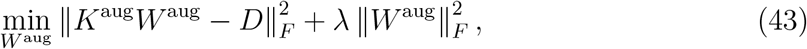

where *D* ∈ ℝ^*n×*3^ stacks the drift targets *d*_*i*_(*t*) for all *n* correspondences.

##### Two-stage baseline plus residual field

As in the per-timepoint RBF variant, we optionally separate a dominant global translation from residual spatial structure. A global translation *g*(*t*) ℝ^3^ is estimated as the componentwise median drift per timepoint and interpolated over time. The space-time RBF model is then fit to residual targets 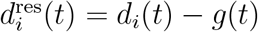, and predictions are combined as

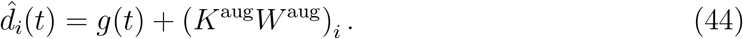

This improves conditioning when global translation dominates the drift.

##### Robust fitting

To reduce sensitivity to mismatches and outliers, robust variants of Eq. (43) are used. In trimmed fitting, we iteratively refit the ridge solution on the fraction *ρ* ∈ (0, 1) of samples with smallest residual norms 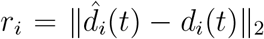. In iteratively reweighted least squares, samples are reweighted using a Huber-type rule based on *r*_*i*_, resulting in a weighted ridge regression at each iteration. A further option trims at the track level by aggregating residual statistics per track across all timepoints and retaining only the best-performing fraction of tracks before refitting.

##### Sequential refinement stages

We implement an optional refinement procedure consisting of multiple stages. Each stage fits a space-time RBF field to the current positions and applies the predicted drift correction to update the positions (excluding the reference timepoint unless row position mode is used). Repeating this procedure can reduce bias introduced by initially imperfect correspondences and can improve fits in row position mode, where predicted drift depends on the position being corrected.

##### Model parameters

The model is controlled by the following parameters. The number of centers *M* sets the expressivity of the field: larger *M* increases spatial and temporal resolution but raises computational cost and the risk of ill-conditioned normal equations. The normalized lengthscales *ℓ*_space_ and *ℓ*_*t*_ determine smoothness in space and time: larger values enforce smoother fields and reduce overfitting, while smaller values allow more local variation but require sufficient center coverage. The ridge parameter *λ* controls the bias-variance tradeoff and stabilizes the linear system. Robustness is governed by the kept fraction *ρ* and the number of robust iterations, or by the corresponding parameters of the reweighting rule. The choice of `space_input_mode`defines the spatial geometry used by the kernel and can strongly influence stability. Finally, the number of refinement stages *N*_refine_ determines how often the correction is recomputed and reapplied, which is mainly relevant in row position mode and in challenging drift regimes.

#### Mixture-of-experts extension of the space-time RBF model

A limitation of a single smooth space-time RBF field is that it enforces one globally consistent set of lengthscales and one global smoothness profile across the full field of view. To increase expressivity while maintaining smooth predictions, we use a mixture-of-experts (MoE) (Jacobs et al., 1991; Jordan and Jacobs, 1994) extension that fits multiple space-time RBF experts on spatially clustered track groups and blends their predictions smoothly in space.

##### Track anchors and clustering

Each track is represented by a 3D anchor position *p*_*j*_ ∈ ℝ^3^. Depending on the chosen mode, *p*_*j*_ is either the temporal mean position of the track (`track_anchor`) or the reference-time position *x*_*j*_(*t*_ref_ ) (`ref_position`), with a fallback to the mean if the reference position is missing. Anchors are partitioned into *K* clusters by *k*-means (Lloyd, 1982), yielding expert centers *c*_*k*_ ∈ ℝ^3^ and a hard assignment *z*_*j*_ ∈ {1, …, *K*} for each track.

##### Shared global translation baseline

Before fitting the experts, we estimate a shared global translation *g*(*t*) ∈ ℝ^3^ using robust per-time medians of the total drift targets,

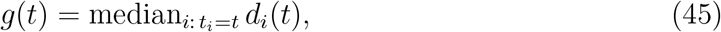

and subtract it from observed positions. Expert models are trained on these globally detrended positions, so that each expert captures primarily residual spatial structure beyond global translation. At inference, the final prediction re-adds the shared global translation.

##### Expert training

For expert *k*, we fit a space-time RBF model *f*_*k*_(*p, t*) ∈ ℝ^3^ using only correspondences originating from tracks assigned to cluster *k*. The expert model is identical to the base space-time RBF formulation, including optional robust fitting such as trimming or iteratively reweighted least squares (IRLS), and optional multi-stage refinement. In practice, the number of RBF centers per expert is capped relative to the available training samples to reduce ill-conditioning.

##### Smooth spatial blending at inference

Given a query position with anchor *p* ∈ ℝ^3^ and time *t*, each expert produces a residual drift prediction *f*_*k*_(*p, t*). We blend these predictions with spatial Gaussian gating weights

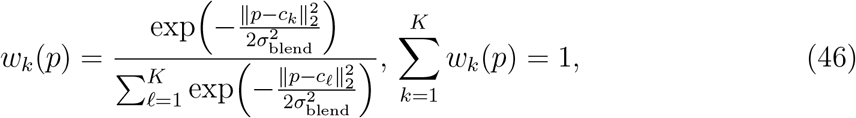

where *σ*_blend_ is specified in physical units (e.g. *µ*m), matching the anchor coordinates.

The final drift prediction is

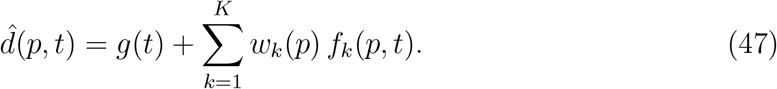

If anchors are constant per track, the weights *w*_*k*_ are constant in time for that track, which stabilizes the correction and avoids temporal switching between experts.

##### Model parameters

The MoE extension introduces parameters in addition to the base space-time RBF hyperparameters. The number of experts *K* controls the spatial granularity of specialization. Larger *K* increases local adaptivity but reduces the sample size per expert and may lead to ill-conditioned fits if clusters become too small. The blending scale *σ*_blend_ sets how smoothly experts transition: small values yield near-hard switching, while large values approach a single global model. The anchor definition (`cluster_mode`) determines whether expert selection is tied to reference position or a track-average anchor. The coupling strength *α* controls optional neighbor regularization. Finally, conditioning thresholds and center caps act as numerical stability parameters by aborting or constraining expert fits when the normal equations become ill-conditioned.

### Identity reconstruction as an assignment problem

The drift models introduced above address geometric consistency across time by mapping observations into a common coordinate system. Even under perfect drift correction, however, the identity of individual objects remains ambiguous whenever multiple spatially proximate candidates exist at a given time point. Identity reconstruction therefore constitutes a second, conceptually distinct inference problem: rather than estimating a continuous deformation field, we now infer a discrete correspondence structure across successive frames.

We formulate identity reconstruction as a sequence of constrained assignment problems. Let *t*_1_ *<* …*< t*_*T*_ denote discrete acquisition times. At each time *t* we observe an unordered point cloud

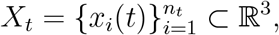

where *x*_*i*_(*t*) represents the spatial coordinate of detection *i* at time *t*, and *n*_*t*_ may vary with *t* due to appearance, disappearance, or missed detections. Our objective is to assign consistent track identities across time. Operationally, for each consecutive transition *t* → *t*^+^ with *t*^+^ = *t*_*s*+1_, we seek a one to one matching between a subset of points in *X*_*t*_ and a subset of points in 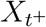. These local correspondences are subsequently composed into global pseudo tracks that represent the inferred identities.

Crucially, the matching between 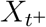 is generally *partial*. Objects may enter or leave the field of view, segmentation may fail at isolated time points, and detections may be spurious. The association problem must therefore permit unmatched points on either side of the transition. The resulting formulation is a discrete optimization problem with inequality constraints rather than a perfect bipartite matching.

This discrete assignment layer is intentionally separated from drift modeling. While drift correction modifies the geometry of the point clouds, the assignment formulation itself does not assume a particular correction method and can operate on raw or corrected coordinates alike. In the following, we describe the cost construction, gating constraints, optimization procedure, and linking strategy that together define our identity reconstruction pipeline.

#### Pairwise cost matrix

For each transition *t* → *t*^+^ we build a pairwise cost matrix 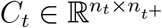, where *C*_*t*_[*i, j*] measures how plausible it is that point *x*_*i*_(*t*) corresponds to point *x*_*j*_(*t*^+^). Our baseline cost is the squared Euclidean distance

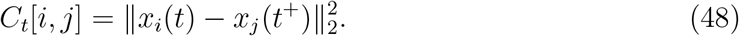

Two optional cost variants are implemented for robustness and anisotropic uncertainty. First, we support the Euclidean distance *C*_*t*_[*i, j*] = ‖*x*_*i*_(*t*) − *x*_*j*_(*t*^+^) ‖_2_. Second, we support a Huberized distance cost (Huber, 1964), defined via *d*_*ij*_ = ‖*x*_*i*_(*t*) − *x*_*j*_(*t*^+^) ‖_2_ and a scale *δ >* 0 as

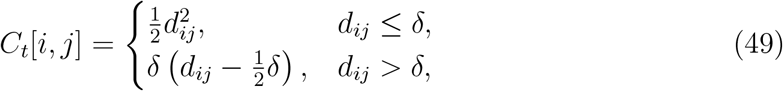

which reduces sensitivity to large displacements. Third, we support a Mahalanobis type cost (Mahalanobis, 1936) with covariance Σ ∈ ℝ^3*×*3^,

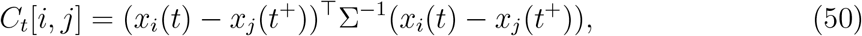

either with a full Σ or a diagonal scaling 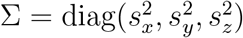.

#### Gating as hard feasibility constraints

Before solving the assignment, implausible correspondences are removed by gating (Bar-Shalom, X.-R. Li and Kirubarajan, 2001). We represent gating by a boolean mask 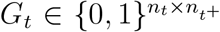, where *G*_*t*_[*i, j*] = 1 indicates that edge (*i, j*) is admissible. We combine a finite value constraint with one or more of the following gates.

A maximum distance gate with threshold *τ*_*D*_ *>* 0,

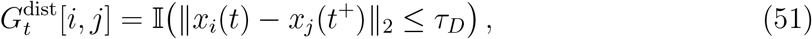

a maximum cost gate with threshold *τ*_*C*_ *>* 0,

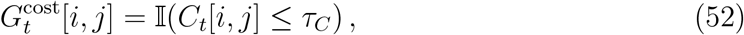

and a percentile gate with percentile *p* ∈ [0, 100], defined on the finite cost entries by

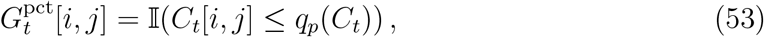

where *q*_*p*_(*C*_*t*_) is the *p*th percentile of {*C*_*t*_[*i, j*] : *C*_*t*_[*i, j*] finite }. The final gate mask is the logical conjunction of the enabled gates and the finite constraint,

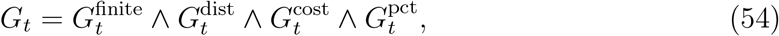

where disabled gates are treated as all ones.

#### Hungarian assignment

For each transition *t* → *t*^+^, we solve a gated linear assignment problem using the Hungarian algorithm (Kuhn, 1955; Munkres, 1957). We introduce binary decision variables *A*_*ij*_ ∈ {0, 1} indicating whether *x*_*i*_(*t*) is matched to *x*_*j*_(*t*^+^). The objective is

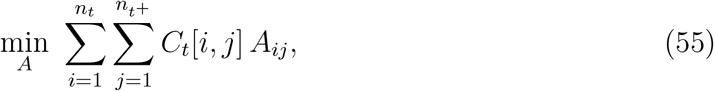

subject to one to one constraints

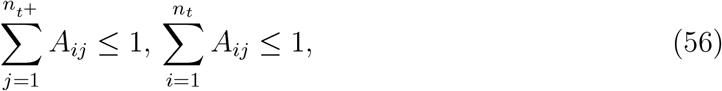

and feasibility constraints

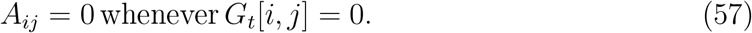

The inequality constraints in Eq. (56) allow partial matchings, so that unmatched points are permitted on either side of the transition.

In practice, the feasibility constraints Eq. (57) are enforced by assigning a large penalty cost to forbidden edges and applying the standard Hungarian solver to the modified cost matrix. After optimization, only assignments corresponding to admissible edges with finite original costs are retained; all remaining points are considered unmatched for that transition.

#### Optional explicit unmatched modeling via dummy nodes

As an alternative formulation, unmatched handling can be modeled explicitly by augmenting the assignment with dummy nodes. This converts the partial matching problem into a square assignment problem of size (*n*_*t*_ +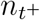 ), in which unmatched source points incur a death penalty *d* and unmatched destination points incur a birth penalty *b*. Concretely, the augmented cost matrix consists of the real gated costs for real to real edges, a diagonal block assigning each source to a dedicated dummy node with cost *d*, and a diagonal block assigning each destination to a dedicated dummy node with cost *b*, while dummy to dummy assignments carry zero cost. Solving the Hungarian algorithm on this augmented system yields both correspondences and unmatched decisions in a single optimization step.

#### Nearest neighbor baseline

As a deterministic baseline, we also consider greedy nearest neighbor matching under the same gate mask. For each source point in row order, we select the admissible unused destination point with minimum cost. Optionally, a miss threshold can be used to leave a source unmatched if its best admissible cost exceeds the threshold.

#### Linking frame to frame assignments into pseudo tracks

The per transition assignment outputs are converted into global pseudo tracks by propagating integer track identifiers through time. At the first frame, each point is assigned a unique track id. For each transition *t* → *t*^+^, if point *i* at time *t* is matched to point *j* at time *t*^+^, then the track id of *i* is assigned to *j*. Unmatched destination points at time *t*^+^ initiate new track ids (birth events). Unmatched source points at time *t* terminate their tracks (death events) in the sense that no further observations are linked to that id. This linking procedure yields a tracks table containing, for each observation, its inferred track id, time, and coordinates. In addition, we retain a link table containing the matched index pairs per transition together with their costs, which supports downstream diagnostics and benchmark metrics. By default, this linking step uses only consecutive frame assignments. Optionally, however, a short-gap post hoc stitching step can be enabled to merge track fragments separated by at most a user specified number of missing intermediate frames. This optional extension operates only at the track-linking stage and does not modify the underlying pairwise assignment, gating, or cost construction.

#### Constant velocity refinement and innovation based gating variants

In addition to the purely geometric frame to frame assignment in Eq. (55) to Eq. (57), we consider a family of focused variants that incorporate a simple constant velocity prediction and, optionally, gating in innovation space. The common structure is iterative. Given a current set of pseudo tracks from a previous association pass, we estimate a per observation velocity *v*_*i*_(*t*) ∈ ℝ^3^ from the last available displacement along the inferred trajectory,

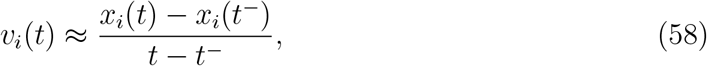

where *t*^−^ denotes the previous time point at which the same identity was observed.

For the next transition *t* → *t*^+^, the predicted position is then

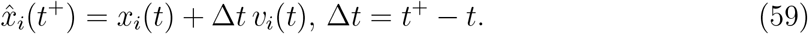

Association is performed between predicted positions 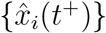 and detected points {*x*_*j*_(*t*^+^)}, after which the resulting assignments are linked into updated pseudo tracks. This refinement is repeated for a small fixed number of iterations.

#### Innovation and normalized innovation squared (NIS)

Define the residual vector for candidate correspondence (*i, j*) as

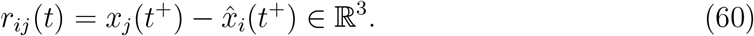

To obtain a dimensionless measure that accounts for anisotropic and time dependent uncertainty, we use the normalized innovation squared (NIS) cost (Bar-Shalom, X.-R. Li and Kirubarajan, 2001)

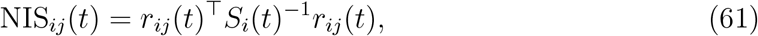

where *S*_*i*_(*t*) ∈ ℝ^3*×*3^ is an innovation covariance. In our implementation *S*_*i*_(*t*) is approximated as diagonal and depends on an estimated per axis innovation scale, measurement noise, process noise, and the time step. Concretely, with per axis innovation standard deviations *σ*(*t*) = (*σ*_*x*_(*t*), *σ*_*y*_(*t*), *σ*_*z*_(*t*)), diagonal measurement noise variances *R* = diag(*r*_*x*_, *r*_*y*_, *r*_*z*_), and diagonal process noise variances *Q* = diag(*q*_*x*_, *q*_*y*_, *q*_*z*_), we use

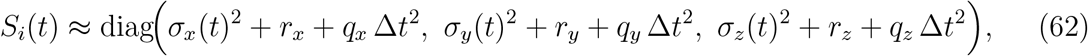

optionally scaled by a mild speed dependent factor to reflect larger prediction uncertainty for faster sources.

#### Robust adaptation of the innovation scale

After solving assignment for a transition *t* → *t*^+^, we update the per axis innovation scale from the residuals of accepted matches. Let 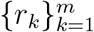 denote the set of residual vectors of matched pairs in that transition. For each axis *a* ∈ {*x, y, z* }, we compute a robust scale estimate using the median absolute deviation (MAD) (Huber, 1964),

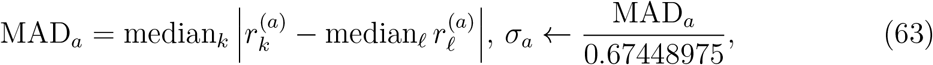

with clipping to a fixed multiplicative window around the initial scale to prevent unstable updates in small sample regimes. The scalar 0.67448975 maps MAD to the standard deviation under a Gaussian reference distribution.

#### Gating in coordinate space and in innovation space

To reduce combinatorial ambiguity and to exclude physically implausible correspondences before solving Eq. (55), we impose a *gating* constraint in one of two conceptually distinct spaces.

In *coordinate space gating*, admissible edges are determined directly from geometric proximity. Using the residual norm ‖*r*_*ij*_(*t*) ‖_2_ from Eq. (60), we define a distance gate

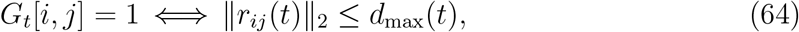

where *d*_max_(*t*) is a user specified threshold or an adaptive bound derived from an estimated innovation scale. Optionally, an additional percentile gate can be applied to the residual norms (or to the underlying costs) to keep only the lowest cost fraction of candidate edges when absolute distance scales vary across experiments.

In *innovation space gating*, admissible edges are determined by the normalized innovation squared (NIS) statistic Eq. (61), which measures residual magnitude relative to an innovation covariance. Given a confidence level *α* ∈ (0, 1), we use a *χ*^2^ threshold with three degrees of freedom,

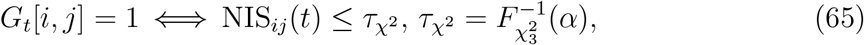

where 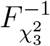 is the inverse cumulative distribution function of 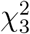. Under the standard Gaussian innovation assumptions, this gate corresponds to retaining candidate matches whose residuals are consistent with the assumed uncertainty model at confidence level *α* (Bar-Shalom, X.-R. Li and Kirubarajan, 2001). As in coordinate space, an optional percentile gate may be applied to NIS_*ij*_(*t*) to suppress weakly supported edges in dense regimes.

#### Variant: Adaptive maximum distance from innovation scale

A limitation of the fixed distance gate Eq. (64) is that the admissible displacement radius *d*_max_ must be chosen a priori and may not reflect time dependent changes in prediction uncertainty. To account for varying innovation scale across transitions, we consider an adaptive distance gate derived from the current innovation standard deviations introduced in Eq. (63). Specifically, in coordinate space we replace the fixed bound *d*_max_ in Eq. (64) by a scale dependent threshold

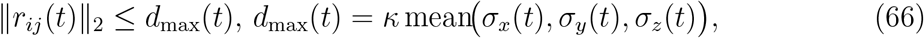

where *κ >* 0 is a user controlled factor.

This construction ties the geometric admissibility region directly to the empirically estimated innovation dispersion of the current transition. If the residual variability increases, the gate expands proportionally; if residuals concentrate, the gate contracts accordingly. Pairwise costs are then computed using the selected metric, and Hungarian assignment is performed under this adaptive gating constraint.

#### Variant: Innovation percentile gating

While the adaptive gate Eq. (66) rescales a Euclidean threshold according to the innovation dispersion, it still operates in raw coordinate space. To obtain a scale invariant decision criterion that normalizes residuals per axis, we consider gating based on a normalized innovation magnitude. Using the residual definition Eq. (60) and the per-axis innovation scales *σ*_*x*_(*t*), *σ*_*y*_(*t*), *σ*_*z*_(*t*), we define the dimensionless quantity

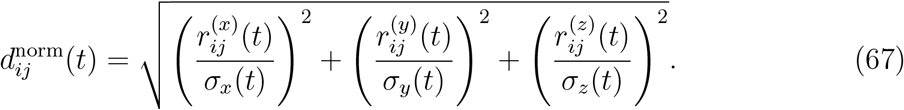

This corresponds to the square root of a diagonal approximation of the NIS statistic in Eq. (61). Admissible edges are selected either by imposing a fixed threshold on 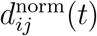 or by retaining only a specified lower percentile of candidate edges. Assignment costs are then taken as 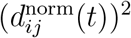.

When full innovation covariance gating is enabled as in Eq. (65), the present construction reduces to operating directly on NIS_*ij*_(*t*), so that gating and costs coincide with the quadratic form induced by *S*_*i*_(*t*)^−1^. In this sense, normalized innovation percentile gating interpolates between purely geometric Euclidean gating and statistically grounded NIS-based gating.

#### Variant: Directional switch penalty

To discourage assignments that reverse motion direction relative to the current velocity estimate, we add a soft penalty to the pairwise costs. Let *v*_*i*_(*t*) be the source velocity and *r*_*ij*_(*t*) the residual. Define the cosine of the angle between them,

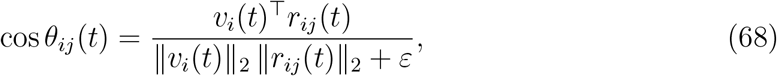

with *ε >* 0 for numerical stability. The modified cost is

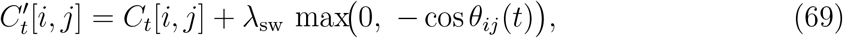

where *λ*_sw_ ≥ 0 controls penalty strength. The gate can be Euclidean or NIS based; the penalty is applied after gating.

#### Variant: Two stage assignment

In dense configurations, a single global assignment under a moderately loose gate may admit several nearly equivalent correspondences, increasing the risk that early mismatches propagate through subsequent linking. To reduce this sensitivity, we consider a two stage association strategy that separates high confidence matches from ambiguous ones.

The procedure decomposes the transition *t* → *t*^+^ into a strict first pass followed by a second pass on the remaining unmatched points. In stage one, a conservative gate is imposed by tightening the admissibility threshold, for example by reducing the Euclidean distance bound in Eq. (64) or by lowering the NIS threshold in Eq. (65). Denoting the strict and relaxed thresholds by *τ* ^(1)^ and *τ* ^(2)^, respectively, we require

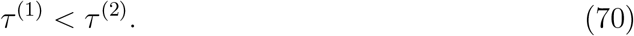

Hungarian assignment is first solved under the strict gate, optionally with explicit unmatched handling. This yields a subset of high confidence matches. In stage two, only the previously unmatched sources and destinations are considered, and assignment is repeated under the relaxed gate *τ* ^(2)^, again optionally allowing explicit unmatched decisions.

The final transition assignment is defined as the union of stage one and stage two matches, while unmatched sets are updated accordingly. This decomposition enforces that the most reliable correspondences are fixed before resolving more ambiguous pairings.

#### Algorithmic note on the Hungarian solver

All variants that solve a one to one assignment rely on the Hungarian algorithm for the underlying linear assignment problem. We refer to the classical formulations and implementations in Kuhn (1955) and Munkres (1957).

#### Notes on probabilistic gating

The NIS gating rule Eq. (65) corresponds to the standard Kalman filtering view in which, under Gaussian assumptions and a correct innovation covariance, the statistic NIS_*ij*_(*t*) follows a *χ*^2^ distribution with degrees of freedom equal to the observation dimension. We use this as a principled way to map a confidence level *α* to a gating threshold in three dimensional space (Bar-Shalom, X.-R. Li and Kirubarajan, 2001).

### Kalman constant velocity association with innovation gating

As a model based alternative to purely geometric gating, we implement a constant velocity (CV) Kalman filter per active track and perform frame to frame association in innovation space. The method maintains a dynamical state for each track and converts candidate correspondences into a normalized innovation squared (NIS) cost, which is then minimized under one to one constraints by Hungarian assignment.

#### State space model

For each track, we use the 6 dimensional CV state

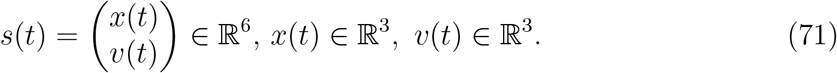

For a time increment Δ*t* = *t*^+^ − *t*, the linear transition and observation models are

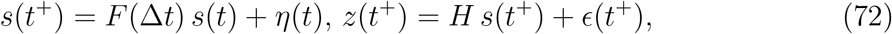

with

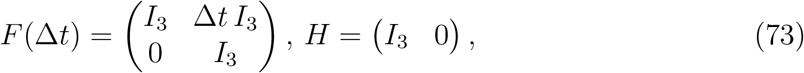

and zero mean Gaussian noises *η*(*t*) ∼ *N* (0, *Q*) and *E*(*t*) ∼ *N* (0, *R*) (Kalman, 1960; Bar-Shalom, X.-R. Li and Kirubarajan, 2001). In our implementation, *Q* and *R* are taken as diagonal and scaled by user parameters, with the position block of *Q* increasing with Δ*t*^2^ to reflect larger prediction uncertainty at larger frame intervals.

#### Prediction and innovation

Let (*s*_*i*_(*t*), *P*_*i*_(*t*)) denote the posterior mean and covariance of track *i* at time *t*. The CV prediction to *t*^+^ is

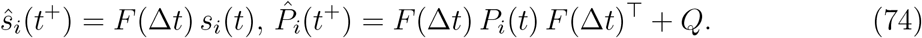

Given a candidate detection *z*_*j*_(*t*^+^) ∈ ℝ^3^, the innovation and its covariance are

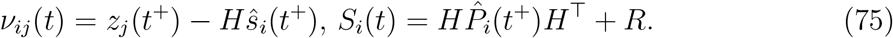

#### NIS cost and innovation gating

We score candidate correspondences by the normalized innovation squared

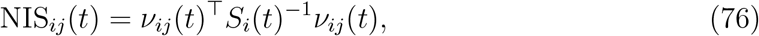

which is a Mahalanobis distance in innovation space (Bar-Shalom, X.-R. Li and Kirubarajan, 2001). Edges are gated by an innovation threshold *τ*,

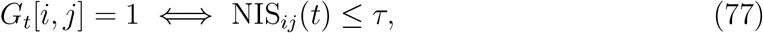

where *τ* is a user parameter. Conceptually, one may also select *τ* via a *χ*^2^ quantile with three degrees of freedom, consistent with Eq. (65), but in the present Kalman implementation *τ* is treated as an explicit hyperparameter.

#### One to one assignment in innovation space

For each transition *t* → *t*^+^, we build the cost matrix

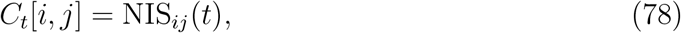

apply the gate Eq. (77) and solve the one to one assignment problem Eq. (55) to Eq. (57) with Hungarian optimization (Kuhn, 1955; Munkres, 1957). In contrast to the geometric baselines where partial matching is obtained by allowing ≤ 1 constraints, the Kalman association optionally activates explicit unmatched modeling via dummy nodes with configurable birth and death penalties, analogously to the construction described above.

#### Kalman update for accepted matches

For each accepted match (*i, j*), we perform the standard linear Kalman update (Kalman, 1960)

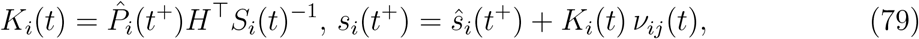

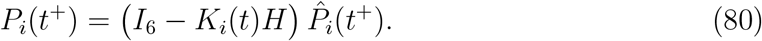

Tracks that are not matched at a transition increase a missed counter. Detections at *t*^+^ that are not assigned to any active track spawn new track states initialized at the observed position with zero velocity and identity covariance. Tracks missed for more than a user specified maximum are pruned from the active set.

#### Practical interpretation

Relative to Euclidean gating, Kalman innovation gating adapts association to locally varying uncertainty induced by the dynamical model and noise scales. The resulting NIS costs incorporate both distance and directional consistency via the predicted state, while retaining a globally optimal one to one matching per transition through Hungarian optimization.

### Global multi frame association via minimum cost path cover

The frame local assignment strategies above decide matches independently for each transition *t* → *t*^+^ and then link these decisions. As a complementary strategy, we implement a global linker that optimizes associations across the entire sequence at once. The central idea is to treat every observed detection as a node in a time ordered directed acyclic graph, to create a set of admissible directed edges between detections at later times, and to select a subset of these edges that forms a collection of disjoint paths. Each path corresponds to one reconstructed pseudo track. This formulation is closely related to classical network flow and path cover approaches for multi object tracking (Zhang, Y. Li and Nevatia, 2008; Pirsiavash, Ramanan and Fowlkes, 2011; Berclaz et al., 2011).

#### Detections and candidate links

Let *D* = {*d*} be the set of all detections across all frames. Each detection *d* is identified by a global index and has an acquisition time *t*(*d*) and a coordinate *x*(*d*) ∈ ℝ^3^. Candidate links connect a detection *u* to a later detection *v* only if *t*(*v*) *> t*(*u*) and if the pair satisfies a geometric gate. In the basic mode, links are created only between consecutive frames; optionally, skip links spanning two frames are allowed to model one missed detection.

For each admissible candidate (*u, v*) we define a link cost *c*(*u, v*) ≥ 0 computed from a selected metric (default squared Euclidean distance) and optionally multiplied by a gap penalty when skip links are used. Birth and death penalties *b >* 0 and *d >* 0 encode the cost of starting or terminating a track.

#### Objective as selecting beneficial links

A convenient way to express the global objective is to compare the cost of linking two detections with the alternative of terminating a track at *u* and starting a new track at *v*. For a candidate edge (*u, v*) with cost *c*(*u, v*), the net gain of using this edge relative to separate death and birth events is

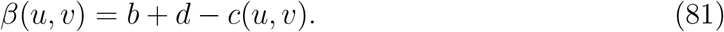

Edges with *β*(*u, v*) ≤ 0 are never beneficial and are therefore discarded.

#### Maximum weight bipartite matching formulation

We now construct a bipartite graph where each detection appears once as a potential predecessor and once as a potential successor. Denote by *D*_*L*_ and *D*_*R*_ copies of, and for every admissible candidate (*u, v*) add an undirected bipartite edge between *u*∈ *D* _*L*_ and *v* ∈ *D*_*R*_ with weight *β*(*u, v*). Selecting a set of edges with no shared endpoints enforces that each detection has at most one successor and at most one predecessor, which is exactly the disjoint path constraint for track reconstruction. We therefore solve

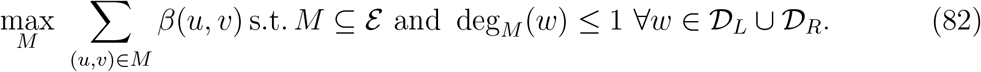

The selected matching induces successor and predecessor maps, and the resulting directed edges form a directed acyclic graph in which every node has in degree at most one and out degree at most one. The connected components are disjoint directed paths and define the reconstructed tracks. This matching based formulation is equivalent to a minimum cost path cover with birth and death penalties under the restriction that candidate links are time forward and that each node can be used at most once as predecessor and successor (Zhang, Y. Li and Nevatia, 2008; Pirsiavash, Ramanan and Fowlkes, 2011; Berclaz et al., 2011).

#### Skip links and gap penalties

To reduce fragmentation when detections are intermittently missing, we optionally allow skip links from frame *t* to *t* + 2. Skip links are penalized by a multiplicative factor

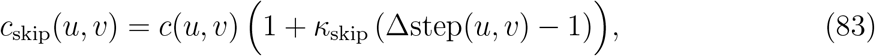

where Δstep(*u, v*) ∈ {1, 2} is the number of frame steps spanned and *κ*_skip_ *>* 0 is a user parameter. This discourages skip links unless they are strongly supported geometrically.

#### Solver choice and deterministic fallback

When a maximum weight matching solver is available, we use it to solve Eq. (82). If such a solver is unavailable, we use a deterministic greedy fallback that sorts candidate edges by decreasing *β*(*u, v*) and selects them if neither endpoint has been used before. The greedy fallback is not guaranteed to be optimal but preserves reproducibility and provides a baseline for environments without graph optimization dependencies.

#### Output as tracks and links

The selected global links are converted into a track ID labeling by assigning a new track ID to each path start node and propagating this ID along successor edges. As in the frame local methods, we output a table of per detection track assignments and a link table listing selected edges and their costs.

### Tracklet first association with stitching

As an additional variant, we implement a two stage strategy that separates identity reconstruction into a conservative local linking stage that aims to minimize false associations, followed by a global stitching stage that merges compatible short track segments into longer trajectories. The motivation is that the most harmful failure mode for identity reconstruction is an early false link that propagates identity errors forward in time, whereas fragmentation can often be repaired if a reliable motion prior is available.

#### Stage one: conservative tracklet construction

In the first stage, we perform frame to frame association with strict gating to obtain short, high confidence tracklets. Concretely, for each transition *t* → *t*^+^ we build a pairwise cost matrix *C*_*t*_ (default squared Euclidean) and an admissibility mask *G*_*t*_ as described above, but with a tightened geometric bound. If a base distance threshold *d*_max_ is provided, the strict gate uses

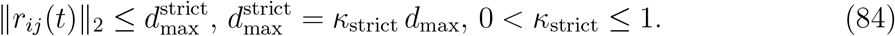

Assignment is then solved either by Hungarian optimization or by a deterministic nearest neighbor baseline. Optionally, explicit unmatched handling via dummy nodes is enabled already in this stage so that ambiguous detections are left unmatched instead of being forced into unreliable links. Linking the accepted transition assignments yields an intermediate set of short tracklets.

#### Tracklet summaries and motion priors

To stitch tracklets, we summarize each tracklet *ℓ* by its start time 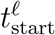, end time 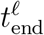, start position 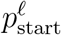, end position 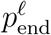, and an end velocity estimate 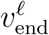. The velocity is computed from the last *K* ≥ 2 tracklet points by finite differences and averaging, providing a simple constant velocity prior that is robust to single frame noise.

#### Stage two: stitching tracklets by endpoint matching

In the second stage, we consider pairs of distinct tracklets (*ℓ*_*a*_, *ℓ*_*b*_) such that *ℓ*_*b*_ starts after *ℓ*_*a*_ ends. Let 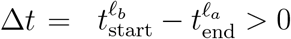. Under a constant velocity prior, the predicted start position of *ℓ*_*b*_ given the end state of *ℓ*_*a*_ is

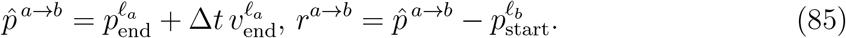

A candidate stitch edge is considered only if Δ*t* does not exceed a prescribed temporal gap tolerance. If gaps are allowed up to *g*_max_ missing frames, the implementation restricts to Δ*t* ≤ *g*_max_ + 1 in units of frame steps.

We then apply a looser gate in this endpoint space, for example a distance bound

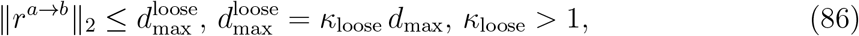

and compute a stitching cost *c*(*ℓ*_*a*_, *ℓ*_*b*_) from the residual *r*^*a*→*b*^. Several options are supported, including squared Euclidean cost, Euclidean cost, a Huber robustification, and diagonal Mahalanobis scaling.

#### Stitching as an assignment problem on tracklets

Let *E* denote the set of tracklet ends and the set of tracklet starts, both indexed by tracklet identity. We construct a cost matrix *C*^stitch^ over admissible end start pairs and solve a one to one assignment using the Hungarian algorithm. Dummy nodes are always enabled in stitching so that tracklets can remain unmerged when no compatible partner exists, controlled by a miss penalty. The resulting matched pairs define stitch edges *ℓ*_*a*_ → *ℓ*_*b*_, which are then filtered deterministically to enforce at most one successor per predecessor and at most one predecessor per successor. Chaining the stitch edges yields a final relabeling of track IDs: each stitched chain is assigned a new global identity, and all detections within the chain inherit this identity.

#### Diagnostics and comparison

The method produces three related quantities: the stage one tracklets, the selected stitch edges, and the final stitched tracks. In evaluation, the stage one link count serves as a proxy for how strict the initial gating was, while the number of stitches quantifies how much fragmentation was repaired in the second stage. The overall identity reconstruction quality is then assessed using the same global metrics as for the other association strategies.

### Benchmarking

Benchmarking serves two purposes in this study. First, it provides a controlled setting in which the true trajectories, drift components, and identity assignments are known, enabling direct quantitative evaluation rather than reliance on proxy measures. Second, it supports a reproducible comparison across heterogeneous operating conditions that mirror common failure modes in microscopy tracking, including spatially varying drift, abrupt regime changes, and ambiguous frame-to-frame associations. All benchmarks are therefore defined as families of synthetic regimes with fixed generative parameters, repeated over multiple independent realizations, and evaluated with task-specific metrics that decompose performance into geometric accuracy, systematic bias, and temporal and identity consistency.

#### Benchmark protocol

Benchmarking was designed to (i) compare methods under controlled, regime specific synthetic conditions, (ii) quantify robustness with respect to stochastic variability, and (iii) support a fair selection of representative hyperparameter settings without embedding extensive sweep descriptions into the Results section. All benchmark experiments follow a common structure: a synthetic regime family defines the data generating process, a method family defines the estimator or association mechanism, and a sweep specification defines which hyperparameters are varied while all remaining settings are kept fixed.

Two benchmark families are considered. The first family evaluates drift correction methods on synthetic trajectory collections that exhibit distinct drift regimes, including globally smooth drift, piecewise smooth drift, abrupt regime changes, rotational components, spatially heterogeneous drift fields, and jump like events. The second family evaluates identity reconstruction and association methods on synthetic point cloud sequences with controlled displacement scales, regimes with heterogeneous displacements varying across tracks and time, appearance and disappearance events, mixtures of migrating and stationary subsets, density variations within a fixed field of view, and an optional setting that couples drift and association in a single experiment.

For each regime family, datasets are generated as multiple independent realizations. Independence is ensured by varying the random seed, while all regime defining parameters are held fixed. Where applicable, multiple files per regime are generated to reduce variance of per regime estimates, and larger benchmark suites additionally sweep over a seed range to obtain regime level distributions rather than single point estimates. The default number of tracks and frames is kept constant across most suites to isolate the effect of regime changes and hyperparameter sweeps.

Within a benchmark suite, all methods are evaluated on exactly the same set of dataset realizations. Hyperparameter sweeps are performed by enumerating predefined grids that target the main sensitivity axes of each method family, for example robustification modes, neighborhood and smoothness scales, kernel length scales, gating thresholds or percentiles, and the number of refinement iterations in motion models. This ensures that comparisons reflect method behavior rather than differences in data exposure.

Outputs are aggregated hierarchically. First, evaluation metrics are computed per dataset realization. Second, results are summarized within each regime by averaging across files and seeds, yielding regime level performance estimates with associated variability. Third, where a single scalar summary is required, regime level results are aggregated across regimes using the same aggregation rule for all methods. Hyperparameter selection for subsequent experiments is performed exclusively from these aggregated benchmark scores, using a fixed selection criterion shared across all methods to avoid implicit tuning advantages.

A compact, reproducible specification of all dataset regimes and executed benchmark suites is provided in the Supplementary Information (Tables 1 to 4), which enable exact regeneration of the reported benchmark results.

#### Synthetic data generation

All benchmarking experiments are performed on controlled synthetic datasets in which the underlying trajectories and the injected drift fields are known exactly. Datasets are generated with a dedicated simulator that produces ground truth trajectories, applies spatially and temporally structured drift fields, and adds measurement noise. This design allows direct evaluation of drift correction and association algorithms against known reference trajectories.

Let *i* ∈ {1, …, *N*} index tracks and *t* ∈ {1, …, *T*} discrete time points. The generator produces ground truth positions

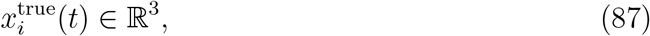

which represent the physical motion of individual objects (e.g. cells). Observed coordinates are then obtained through an additive observation model

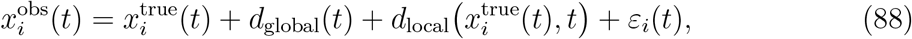

where *d*_global_(*t*) is a global translational drift component, *d*_local_(*x, t*) is a spatially varying drift field, and *ε*_*i*_(*t*) ∼ *N* (0, *σ*^2^*I*_3_) models measurement noise. The synthetic datasets therefore explicitly separate intrinsic motion from imaging drift.

##### Ground truth cell motion

Intrinsic trajectories are generated as a mixture of motion types representing different biological behaviors. Each track is assigned a motion class (stationary, migrating, or slow-moving) with user-defined fractions. Trajectories evolve according to a velocity process with correlated random increments,

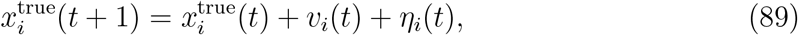

where *v*_*i*_(*t*) represents a persistent component of motion and *η*_*i*_(*t*) is a stochastic displacement term. Migrating tracks exhibit persistent velocity components, while stationary tracks are constrained to near-zero displacement. Additional oscillatory and zigzag perturbations introduce curvature and non-linear motion patterns. Motion amplitudes can be controlled through a displacement scale parameter *dx*, allowing regimes with homogeneous or heterogeneous motion magnitudes.

Optional birth–death processes introduce track appearance and disappearance events. Each track may start at time *t*_birth_ and terminate at time *t*_death_, subject to a minimum lifetime constraint. Birth events may occur randomly or in spatial proximity to existing tracks, allowing simulation of cell division–like scenarios.

##### Global drift component

The global drift component represents stage motion or sample translation. It is generated as a temporally correlated random walk,

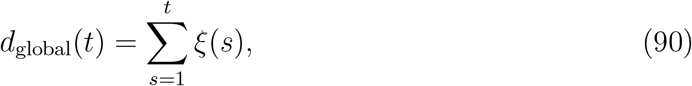

where the increments *ξ*(*s*) follow a temporally smoothed Gaussian process whose variance controls the drift amplitude along each spatial axis.

##### Spatially varying drift fields

Local drift is modeled using smooth spatial basis functions. Let 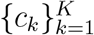 denote spatial centers and let

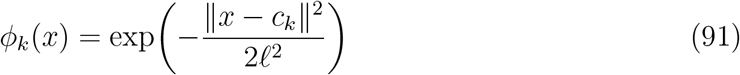

be Gaussian radial basis functions with length scale *ℓ*. The spatial drift field is then expressed as

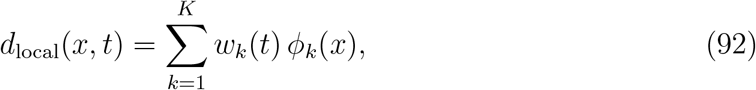

where the coefficients *w*_*k*_(*t*) evolve smoothly in time. This construction produces low-frequency spatial deformation fields similar to microscope stage distortions or sample deformation.

Several alternative drift regimes are implemented for robustness testing:

- smooth spatial drift fields generated by the RBF model,
- piecewise spatial drift fields with region-dependent motion,
- global rotational drift around a fixed center,
- abrupt jump events that introduce discontinuous stage shifts.

These regimes allow evaluation under both smooth and non-smooth drift conditions.

##### Noise model

Measurement noise is applied independently to each coordinate,

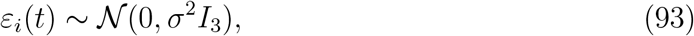

with *σ* typically on the order of 0.1–0.2 *µ*m. This noise level approximates typical localization uncertainty in microscopy tracking experiments.

##### Dataset profiles and regimes

Benchmark datasets are organized into predefined profiles that vary motion statistics, drift regimes, and experimental scale. Each profile specifies a fixed parameter configuration and replication scheme. The drift correction experiments use the profiles listed in Table 1, while identity reconstruction and association experiments use the profiles in Table 2. Across these profiles, the number of tracks is typically on the order of 80, the sequence length is *T* = 20 frames, and multiple seeds are used to generate independent realizations.

For each generated dataset the simulator outputs

- the observed trajectories 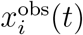,
- the drift-free ground truth trajectories 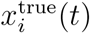,
- and the exact drift field components *d*_global_(*t*) and *d*_local_(*x, t*).

This information enables exact quantitative evaluation of both drift correction accuracy and identity reconstruction performance.

#### Evaluation metrics

Evaluation metrics are defined on synthetic benchmarks where ground truth is available for both drift correction and identity reconstruction tasks. The metric set was chosen to quantify complementary aspects of performance: geometric fidelity of reconstructed trajectories relative to ground truth, systematic drift bias, temporal stability of errors, robustness across heterogeneous subsets such as stationary versus migrating objects, and identity consistency plus link correctness in association problems. All metrics are computed after restricting evaluation to matched records, using an inner join on the common index (TrackID, *t*), so that every term is defined on pairs that exist in both inferred and ground truth outputs.

##### Drift-correction benchmarks

Let 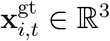 denote the ground truth position of track *i* at time *t*, and 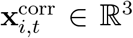 the drift-corrected estimate produced by a method. After inner-joining on (*i, t*), the per-sample Euclidean error norm is

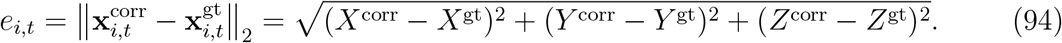

The primary accuracy metric is the overall root-mean-square error

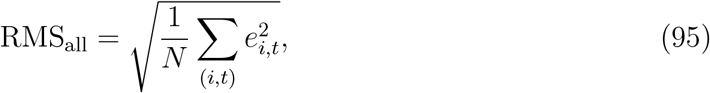

where *N* is the number of matched (*i, t*) pairs. This metric directly measures the geometric fidelity of corrected trajectories and is the principal selector for hyperparameter sweeps and baseline comparisons across drift regimes.

To complement RMS_all_ with robust distributional summaries of pointwise error magnitudes, we additionally report median error across (Med) all matched samples,

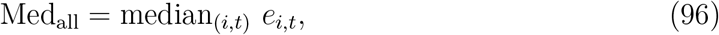

and the upper-tail quantile

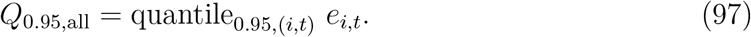

These summaries distinguish typical error levels (median) from tail behavior (p95), which is important when a minority of difficult trajectories dominates RMS.

Per-time metrics are computed analogously. Let *N*_*t*_ be the number of matched samples at time *t*. Then

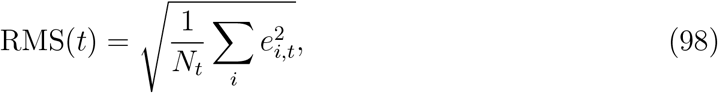

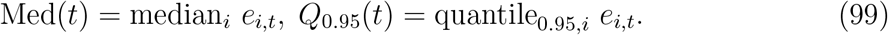

These temporal summaries are used in targeted sweeps to separate central-tendency improvements from heavy-tail failure modes over time.

Several synthetic regimes label a subset of trajectories by a categorical type CellType ∈ {stationary, migrating}. Conditional RMS metrics are then computed as

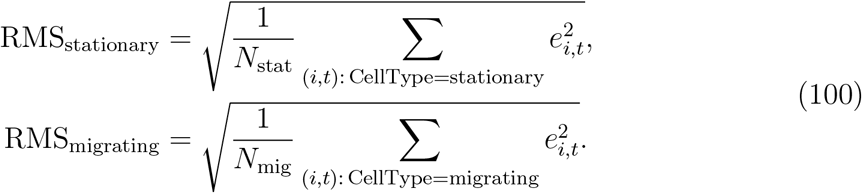

These quantities are used in large-regime benchmark summaries and in fairness studies to verify that drift correction does not preferentially overfit one subset, for example by anchoring to slowly moving trajectories at the expense of migrating ones.

In addition to geometric accuracy, drift benchmarks quantify drift bias. Let 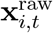 denote the uncorrected observed position and define the method-implied predicted drift vector per sample as

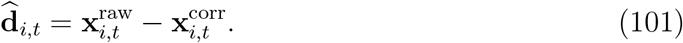

Let 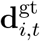 denote the ground truth total drift vector. A per-time robust drift estimate is obtained by taking the component-wise median over tracks, yielding 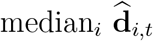 and 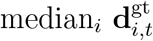. The per-time drift bias magnitude is then

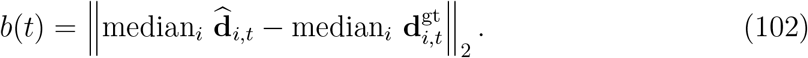

A scalar drift-bias summary is reported as the mean over time,

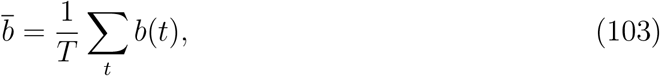

and is used alongside RMS_all_ in summary plots and aggregated sweep tables. Drift bias is intentionally computed using medians to reduce the influence of outlier trajectories or occasional mismatches, thereby measuring systematic drift misestimation rather than local tracking noise.

Finally, benchmark outputs include count statistics that contextualize metric values: the number of matched points *n*_points_ = *N* after the inner join, and the number of tracks *n*_tracks_ contributing to per-track summaries. These are not used as sweep selectors but serve as quality control indicators in aggregated reports.

##### Association and identity reconstruction benchmarks

Association benchmarks evaluate methods that reconstruct track identities from per-frame point clouds. Let *G* denote the set of ground truth detections and *G* the inferred detections linked into inferred tracks. Evaluation uses the merged inferred-ground truth table produced by matching detections at each time step and associating them with both a ground truth track identity GT_TrackID and an inferred track identity TrackID. Metrics quantify identity consistency and link correctness across time.

Fragmentation quantifies how often a ground truth track is split into multiple inferred tracks. Let *f* (*g*) be the number of distinct inferred track IDs associated with ground truth track *g* in the merged table. Then

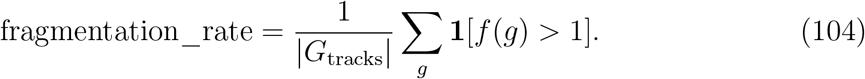

Identity switches quantify temporal instability of inferred identities along a ground truth track. For each ground truth track *g*, order matched points by time and count transitions where the inferred track ID changes:

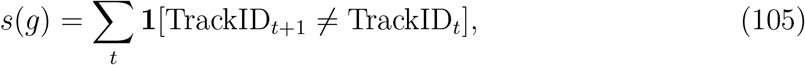

and summarize as a mean per track,

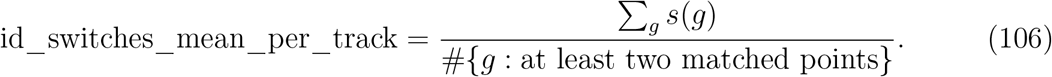

Link-based metrics assess whether inferred temporal links are correct relative to ground truth identity continuity. For each transition *t* → *t* + 1, define inferred links between detections, and define a link as correct if it connects two detections that share the same ground truth identity and are also linked within the same inferred track. Denote by *n*_inf_ (*t*) the number of inferred links at transition *t*, by *n*_corr_(*t*) the number of correct links, and by *n*_gt_(*t*) the number of expected ground truth links. Then

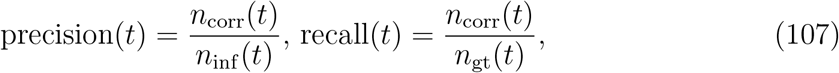

with the convention that the ratio is set to zero if the denominator is zero. The per-time F1 score is

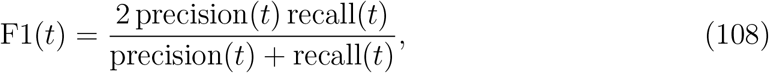

again set to zero if the denominator is zero. The reported scalar metrics are means over transition times with nontrivial link counts,

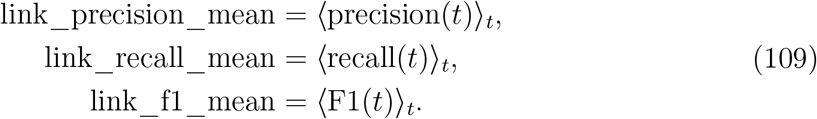

These metrics are central for association sweeps because they directly reflect whether an algorithm reconstructs correct temporal continuity, independent of longer-range track structure.

Unmatched rates quantify incompleteness in reconstructed tracks. For each time *t*, let *n*_gt_(*t*) be the number of ground truth tracks present and *n*_unmatched_(*t*) the number not matched by inferred tracks. The mean unmatched rate is reported as

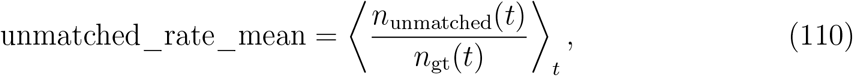

using a NaN-aware mean over times where *n*_gt_(*t*) *>* 0.

##### Auxiliary drift metrics

Additional exploratory summaries are exported during drift postprocessing, but they are used only for qualitative inspection and not as primary sweep-selection criteria.

## Data and code availability

The code used for synthetic benchmarking, biological-example processing, and figure generation is available at https://github.com/FabrizioMusacchio/driftcorr and archived on Zenodo (Musacchio and Fuhrmann, 2026). The biological datasets analyzed in the present study are publicly available from Zenodo: the carcinoma-cell segmentation benchmark dataset by Kapoor (Kapoor, 2022) and the paired bright-field/DAPI synoviocyte time-lapse dataset by Radvansky *et al*. (Radvansky et al., 2026).

## Acknowledgements

We thank Varun Kapoor for making the carcinoma-cell benchmark dataset publicly available, and Martin Radvansky, Markéta Vašinková, Miloš Kudělka, Eva Kriegová, and Petr Gajdoš for making the paired bright-field/DAPI synoviocyte time-lapse dataset publicly available. Their open data resources made the biological validation analyses possible.

## Author contributions

F.M. and M.F. conceived the study. F.M. developed the mathematical framework, implemented the analysis software, designed and performed the synthetic benchmarks, processed and analyzed the biological example datasets, generated the figures, and wrote the original manuscript draft. M.F. supervised the project, acquired funding, contributed to study design and interpretation of the results, and reviewed and edited the manuscript. Both authors discussed the results and approved the final manuscript.

## Use of AI-assisted tools

The authors used OpenAI ChatGPT, GPT-5, to assist with language editing and wording refinement. The tools were not used to generate primary research data, perform unsupervised interpretation of results, or make authorship-level scientific decisions. All AI-assisted text changes were reviewed, verified, and edited by the authors, who take full responsibility for the accuracy, integrity, and content of the submitted article.

## Supplementary Information: Benchmarking overview

### Benchmark scripts and scope

Benchmark suites were executed via three driver scripts that instantiate synthetic regimes, enumerate hyperparameter sweeps, execute methods, and aggregate outputs:

- `user_scripts/parameter_sweep_on_synthetic_data.py`
- `user_scripts/parameter_sweep_on_synthetic_non_rigid_data.py`
- `user_scripts/benchmark_point_cloud_association.py`

### Synthetic dataset profiles used in benchmarking

**Table 1.** Synthetic dataset profiles for drift correction benchmarks. Regime labels are internal identifiers used for reproducible execution.

| Profile | Regimes | Core parameters and replication |
| --- | --- | --- |
| DS-DRIFT-1 | smooth, piecewise<br>smooth, piecewise<br>abrupt | Target: 5 files per regime. Generation: $n\_tracks = 80$, $n\_times = 20$, seed = 42. Motion mix: stationary 0.45, migrating 0.35, smoothness 0.9. Used in Figures 1, 2, and 3, and in Supplementary Figure 12. |
| DS-DRIFT-2 | preset non rigid regimes | Per regime, per seed folders with presets: rotation drift: piecewise regions 2, transition $120.0\ \mu\text{m}$, lengthscale $220.0\ \mu\text{m}$, drift scale $2.0\ \mu\text{m}$, measurement noise $0.10\ \mu\text{m}$. Piecewise spatial drift: piecewise regions 3, transition $65.0\ \mu\text{m}$, lengthscale $180.0\ \mu\text{m}$, drift scale $2.8\ \mu\text{m}$, measurement noise $0.12\ \mu\text{m}$. Jump events: piecewise regions 3, transition $25.0\ \mu\text{m}$, lengthscale $170.0\ \mu\text{m}$, drift scale $2.4\ \mu\text{m}$, measurement noise $0.13\ \mu\text{m}$. Additional defaults: $n\_tracks = 80$, $n\_times = 20$, $n\_seeds = 10$, seed start 42, stationary 0.45, migrating 0.35, smoothness 0.9. Used in Figure 4 and Supplementary Figure 13. |

**Table 2.** Synthetic dataset profiles for association and identity reconstruction benchmarks.

| Profile | Regimes | Core parameters and replication |
| --- | --- | --- |
| DS-PCA-1 | no drift, fixed displacement scale | Blind pairs with displacement scales [0.2, 0.5, 1.0, 2.0, 4.0]. Defaults: $N\_FILES = 3$, $N\_TRACKS = 80$, $N\_TIMES = 20$, $BASE\_SEED = 42$. Motion mix: stationary 0.10, migrating 0.85, smoothness 0.9. Displacements: Gaussian, track smoothness 0.7. Used in Figure 5 and Supplementary Figure 14. |
| DS-PCA-2 | no drift, varying displacement | Varying per track and time: $dx\_min = 0.2\ \mu m$, $dx\_max = 4.0\ \mu m$, uniform distribution, nominal $dx\_scale = 1.0\ \mu m$. Used in Figure 6. |
| DS-PCA-3 | birth and death with stationary subset | Combined with varying displacement, explicit birth and death events, minimum lifetime 5 frames, late-birth and early-death probabilities 0.35, random birth probability 0.5, and a stationary fraction of 0.25 realized as a hard-zero motion subset. Used in Figure 7. |
| DS-PCA-4 | density sweep in fixed volume | Volume size (700, 700, 350), density scales 0.5 to 20.0, initial tracks 80. Combined with varying displacement, birth and death, and stationary settings. Used in Figure 9. |
| DS-PCA-5 | birth and death with drift | Varying displacement and birth and death with an added smooth translation-plus-rotation drift schedule, using the dedicated drift and motion-mix presets from the benchmark generator. Used in Figure 8. |

### Executed benchmark suites

**Table 3.** Benchmark suites executed for drift correction, including the primary sweep axes.

| Suite | Dataset profiles | Methods and sweep highlights |
| --- | --- | --- |
| Baseline suite comparisons | DS-DRIFT-1 | Rigid stepwise, local communities, RBF to reference, space time RBF, and space time RBF mixture of experts. Mixture of experts compared with $k \in \{2, 3, 5\}$. Used in Figures 1 and 2. |
| Baseline fairness sweeps | DS-DRIFT-1 | Rigid stepwise and local-community hyperparameter grids. Rigid: keep fraction $\{0.5, 0.7, 0.85, 0.95\}$ and reestimate iterations $\{5, 15, 30\}$. Local: $k$ neighbors $\{8, 12, 20, 30\}$ and field lengthscale $\{80, 120, 180, 240\} \mu\text{m}$. Used in Supplementary Figure 12. |
| Structured non-rigid suite comparisons | DS-DRIFT-2 | Rigid stepwise, local communities, RBF to reference, space time RBF, and space time RBF mixture of experts on the preset rotation, piecewise-spatial, and jump-event regimes. Used in Figure 4. |
| Structured non-rigid RBF-family sweeps | DS-DRIFT-2 | Stress-suite counterpart of the baseline RBF-family analyses, including robust-mode, expressivity, and stationary-subset evaluations for the structured non-rigid presets. Used in Supplementary Figure 13. |
| RBF robust mode sweep | DS-DRIFT-1 | Robust mode comparison: sample trimming versus IRLS Huber. Used in Figure 3. |
| Space time RBF robust mode sweep | DS-DRIFT-1 | Robust mode comparison: sample trimming, IRLS Huber, and a hybrid track IRLS mode. Used in Figure 3. |
| Space time RBF expressivity sweep | DS-DRIFT-1 | Model expressivity toggles: anisotropy off versus on, linear augmentation off versus on. Used in Figure 3. |
| Space time RBF parameter grid | DS-DRIFT-1 | Grid: $\ell_{\text{space}} \in \{1.0, 2.0, 4.0\}$, $\ell_{\text{time}} \in \{0.4, 0.8, 1.6\}$, typically with 10 seeds. Used in Figure 3. |
| Space time RBF stationary subset tests | DS-DRIFT-1 | Candidate settings $(\ell_{\text{space}}, \ell_{\text{time}}) = (4.0, 0.4)$ and $(1.0, 1.6)$ with stationary subset toggled. Used in Figure 3. |

**Table 4.**
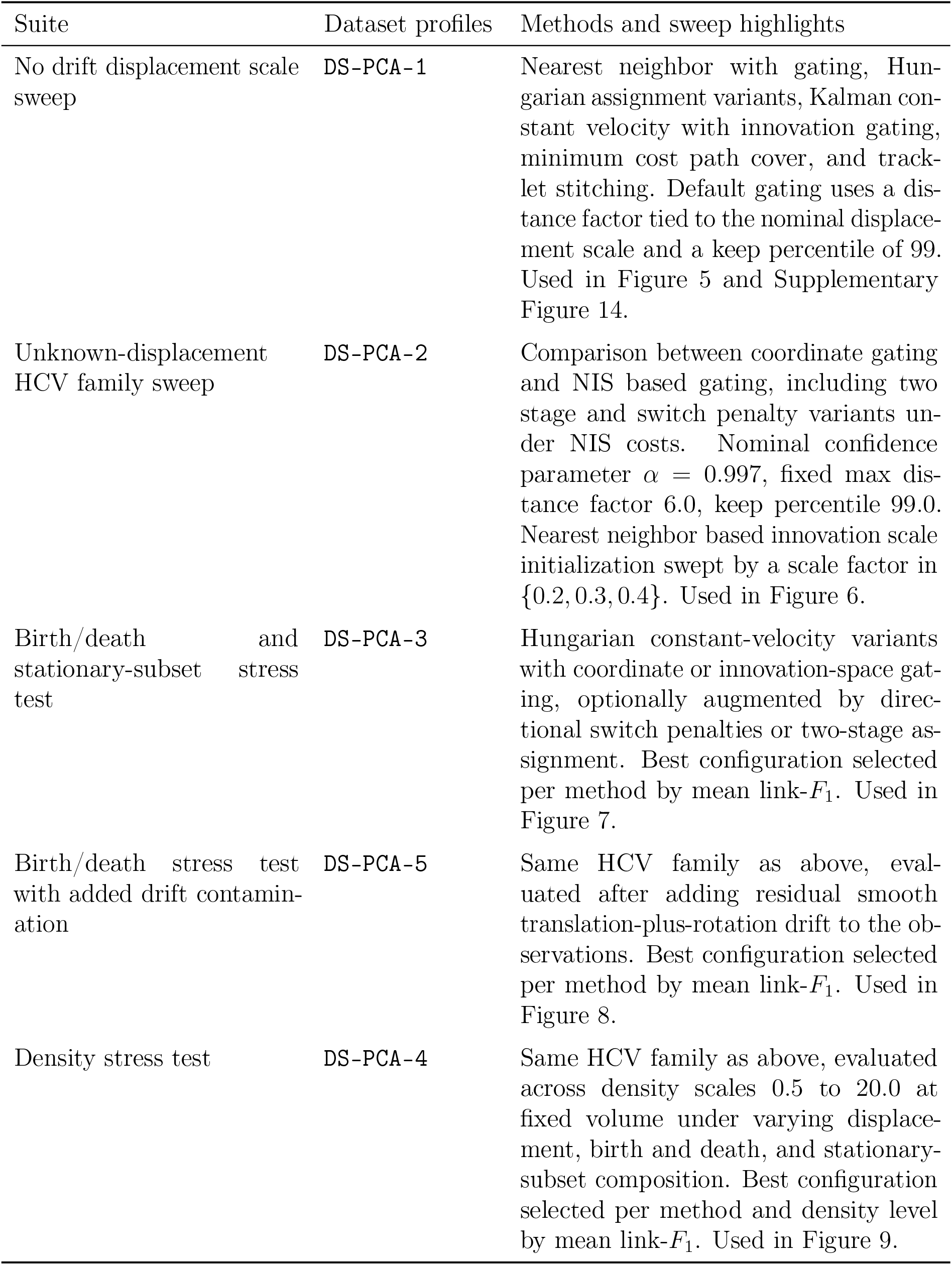
Benchmark suites executed for association and identity reconstruction.

## Supplementary Information synthetic drift-correction benchmarks

### Supplementary Information for baseline spatially varying drift benchmarks

To facilitate direct numerical inspection of the aggregate benchmark results shown in Figure 2, Table 5 reports the exact mean values underlying the heatmaps. The table contains the mean overall RMS error, mean median error, mean *p*95 error, and mean absolute drift-bias error for all seven correction methods across the three baseline drift modes `smooth`, `piecewise_smooth`, and `piecewise_abrupt`. All values are given in *µ*m and were averaged over the five synthetic datasets generated for each mode from profile `DS-DRIFT-1`.

**Table 5:** Exact numerical values underlying Figure 2. Mean benchmark values for all method–mode combinations in the baseline suite. For each of the three drift modes, the table reports mean overall RMS error, mean median error, mean *p*95 error, and mean absolute drift-bias error. All values are reported in *µ*m and correspond to the aggregate heatmap values shown in Figure 2.

| Method | smooth |  |  |  | piecewise_smooth |  |  |  | piecewise_abrupt |  |  |  |
| --- | --- | --- | --- | --- | --- | --- | --- | --- | --- | --- | --- | --- |
| | RMS | Median | $p95$ | Bias | RMS | Median | $p95$ | Bias | RMS | Median | $p95$ | Bias |
| `rigid` | 2.548 | 2.483 | 3.384 | 2.311 | 4.252 | 3.948 | 6.094 | 3.251 | 4.355 | 4.002 | 6.280 | 3.150 |
| `local` | 2.580 | 2.483 | 3.495 | 2.268 | 4.284 | 3.833 | 6.245 | 3.213 | 4.493 | 3.996 | 6.528 | 3.213 |
| `rbf` | 4.484 | 3.343 | 7.979 | 2.507 | 5.976 | 5.067 | 10.048 | 3.486 | 6.142 | 5.272 | 10.257 | 3.430 |
| `st_rbf` | 2.780 | 2.634 | 4.003 | 2.273 | 4.685 | 4.156 | 7.125 | 3.195 | 4.850 | 4.282 | 7.373 | 3.129 |
| `st_rbf_moe_k2` | 3.032 | 2.696 | 4.622 | 2.392 | 4.736 | 4.195 | 7.286 | 3.208 | 4.896 | 4.332 | 7.511 | 3.098 |
| `st_rbf_moe_k3` | 3.548 | 3.004 | 5.562 | 2.537 | 5.113 | 4.460 | 7.975 | 3.175 | 5.318 | 4.444 | 8.406 | 3.019 |
| `st_rbf_moe_k5` | 4.536 | 3.444 | 7.970 | 2.836 | 6.158 | 5.076 | 10.641 | 3.778 | 6.396 | 5.223 | 10.996 | 3.722 |

The baseline-suite comparison in the main text uses fixed hyperparameters for all correction models in order to ensure that the ranking reflects intrinsic model behavior rather than differences in tuning exposure. To verify that the rigid and local baselines were not advantaged or disadvantaged by these default settings, we performed additional parameter sweeps for both baseline families. These sweeps serve purely as a fairness and robustness check and are therefore reported only in the Supplementary Information.

Figure 12 summarizes the results. For both models, the sweeps were evaluated on the same three spatially varying drift regimes used in the main baseline suite (smooth, piecewise smooth, piecewise abrupt), using identical synthetic datasets and aggregating metrics across benchmark realizations. The primary comparison metric is the mean overall corrected-versus-ground-truth RMS error.

#### Rigid model sweep

For the rigid stepwise model (Eq. (5), Eq. (11)), we varied the parameters controlling the trimmed robust estimation procedure described in Eq. (8). Specifically, we swept the inlier keep fraction *ρ* ∈ { 0.5, 0.7, 0.85, 0.95} and the number of inlier re-estimation iterations {5, 15, 30} . The results are shown in Figure 12a.

Across all three regimes, the rigid model shows only weak sensitivity to these parameters. The dominant effect is the keep fraction *ρ*: values near *ρ* ≈ 0.85 tend to produce the lowest mean RMS, whereas very aggressive trimming (*ρ* = 0.5) or almost no trimming (*ρ* = 0.95) are slightly worse. In contrast, the number of inlier re-estimation iterations has negligible influence once a few iterations are performed. This behavior is expected because the trimmed Kabsch fitting procedure converges quickly once a stable inlier set is identified.

Importantly, the overall performance variation across the full sweep is small. Averaged across regimes, the best configuration improves the mean overall RMS by less than one percent relative to the default configuration used in the baseline-suite experiments. Thus the rigid model’s strong performance in the main benchmark is not the result of a narrowly tuned parameter choice.

#### Local community model sweep

In contrast to the rigid model, the local method exhibits a clearer parameter dependence. The dominant control parameter is the field blending scale *ℓ*_field_, which governs the spatial smoothness of the predicted drift field in Eq. Eq. (21). Larger values consistently reduce the RMS across regimes, with *ℓ*_field_ = 240 *µ*m performing best in the present sweep. The number of neighbors *k* plays a secondary role; once *ℓ*_field_ is sufficiently large, the performance differences between *k* = 12, 20, and 30 are minimal.

Even in this case, however, the overall effect size remains modest. The best configuration improves the mean overall RMS by approximately five percent relative to the default parameter choice used in the main baseline suite. While this indicates that the local model benefits somewhat from increased spatial smoothing, the change does not alter the qualitative ranking of methods observed in the main results.

#### Fairness interpretation

The purpose of these sweeps is not to identify globally optimal parameter settings but to verify that the baseline comparison is not driven by arbitrary defaults. Across the explored parameter ranges, both baseline models show stable behavior and relatively shallow performance landscapes. The rigid model is particularly insensitive to parameter variation, and although the local model benefits from somewhat larger spatial smoothing scales, the overall benchmark conclusions remain unchanged. The strong performance of the rigid and local baselines in the main experiments therefore reflects intrinsic model robustness rather than favorable hyperparameter tuning.

### Supplementary Information for RBF-family benchmarks

The following tables report the exact numerical values underlying the parameter sweeps and ablation analyses shown in Figure 3. All values are aggregated across benchmark realizations for the respective synthetic regimes.

#### Space-time RBF parameter grid sweep

Table 6 reports the aggregated results of the (*ℓ*_space_, *ℓ*_*t*_) grid sweep shown in Figure 3a. The table lists the regime, the parameter pair (*ℓ*_space_, *ℓ*_*t*_), the mean global RMS error 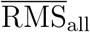, and the mean absolute drift-bias 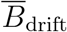.

**Table 6:** Aggregated results of the ST-RBF parameter grid sweep. Values correspond to the heatmaps shown in Figure 3a.

| Regime | $(\ell_{\text{space}}, \ell_t)$ | RMS | Drift | Regime | $(\ell_{\text{space}}, \ell_t)$ | RMS | Drift |
| --- | --- | --- | --- | --- | --- | --- | --- |
| piecewise abrupt | (1.0, 0.1) | 7.378 | 5.090 | piecewise smooth | (6.0, 0.4) | 4.800 | 4.338 |
| piecewise abrupt | (1.0, 0.2) | 7.448 | 5.070 | piecewise smooth | (6.0, 0.8) | 4.781 | 4.302 |
| piecewise abrupt | (1.0, 0.4) | 7.547 | 5.097 | piecewise smooth | (6.0, 1.6) | 4.723 | 4.190 |
| piecewise abrupt | (1.0, 0.8) | 7.605 | 5.149 | piecewise smooth | (6.0, 3.2) | 4.835 | 4.591 |
| piecewise abrupt | (1.0, 1.6) | 7.694 | 5.156 | piecewise smooth | (8.0, 0.1) | 5.011 | 4.530 |
| piecewise abrupt | (1.0, 3.2) | 7.804 | 5.301 | piecewise smooth | (8.0, 0.2) | 4.938 | 4.508 |
| piecewise abrupt | (2.0, 0.1) | 7.212 | 5.335 | piecewise smooth | (8.0, 0.4) | 4.895 | 4.489 |
| piecewise abrupt | (2.0, 0.2) | 7.241 | 5.312 | piecewise smooth | (8.0, 0.8) | 4.862 | 4.453 |
| piecewise abrupt | (2.0, 0.4) | 7.278 | 5.284 | piecewise smooth | (8.0, 1.6) | 4.778 | 4.321 |
| piecewise abrupt | (2.0, 0.8) | 7.291 | 5.213 | piecewise smooth | (8.0, 3.2) | 4.848 | 4.481 |
| piecewise abrupt | (2.0, 1.6) | 7.363 | 5.203 | piecewise smooth | (10.0, 0.1) | 5.101 | 4.628 |
| piecewise abrupt | (2.0, 3.2) | 7.462 | 5.314 | piecewise smooth | (10.0, 0.2) | 5.028 | 4.610 |
| piecewise abrupt | (4.0, 0.1) | 7.146 | 5.314 | piecewise smooth | (10.0, 0.4) | 4.983 | 4.592 |
| piecewise abrupt | (4.0, 0.2) | 7.157 | 5.302 | piecewise smooth | (10.0, 0.8) | 4.947 | 4.558 |
| piecewise abrupt | (4.0, 0.4) | 7.165 | 5.297 | piecewise smooth | (10.0, 1.6) | 4.855 | 4.429 |
| piecewise abrupt | (4.0, 0.8) | 7.172 | 5.294 | piecewise smooth | (10.0, 3.2) | 4.884 | 4.456 |
| piecewise abrupt | (4.0, 1.6) | 7.216 | 5.267 | smooth spacetime | (1.0, 0.1) | 3.934 | 3.972 |
| piecewise abrupt | (4.0, 3.2) | 7.309 | 5.283 | smooth spacetime | (1.0, 0.2) | 3.972 | 4.047 |
| piecewise abrupt | (6.0, 0.1) | 7.164 | 5.386 | smooth spacetime | (1.0, 0.4) | 4.080 | 4.143 |
| piecewise abrupt | (6.0, 0.2) | 7.160 | 5.399 | smooth spacetime | (1.0, 0.8) | 4.250 | 4.187 |
| piecewise abrupt | (6.0, 0.4) | 7.163 | 5.398 | smooth spacetime | (1.0, 1.6) | 4.465 | 4.323 |
| piecewise abrupt | (6.0, 0.8) | 7.166 | 5.392 | smooth spacetime | (1.0, 3.2) | 4.595 | 4.353 |
| piecewise abrupt | (6.0, 1.6) | 7.208 | 5.368 | smooth spacetime | (2.0, 0.1) | 2.988 | 3.420 |
| piecewise abrupt | (6.0, 3.2) | 7.306 | 5.331 | smooth spacetime | (2.0, 0.2) | 2.972 | 3.423 |
| piecewise abrupt | (8.0, 0.1) | 7.184 | 5.405 | smooth spacetime | (2.0, 0.4) | 2.959 | 3.424 |
| piecewise abrupt | (8.0, 0.2) | 7.182 | 5.414 | smooth spacetime | (2.0, 0.8) | 2.969 | 3.422 |
| piecewise abrupt | (8.0, 0.4) | 7.182 | 5.411 | smooth spacetime | (2.0, 1.6) | 2.991 | 3.418 |
| piecewise abrupt | (8.0, 0.8) | 7.184 | 5.405 | smooth spacetime | (2.0, 3.2) | 3.058 | 3.410 |
| piecewise abrupt | (8.0, 1.6) | 7.221 | 5.376 | smooth spacetime | (4.0, 0.1) | 3.505 | 3.438 |
| piecewise abrupt | (8.0, 3.2) | 7.319 | 5.337 | smooth spacetime | (4.0, 0.2) | 3.436 | 3.435 |
| piecewise abrupt | (10.0, 0.1) | 7.198 | 5.415 | smooth spacetime | (4.0, 0.4) | 3.327 | 3.429 |
| piecewise abrupt | (10.0, 0.2) | 7.198 | 5.421 | smooth spacetime | (4.0, 0.8) | 3.164 | 3.423 |
| piecewise abrupt | (10.0, 0.4) | 7.198 | 5.417 | smooth spacetime | (4.0, 1.6) | 3.026 | 3.458 |
| piecewise abrupt | (10.0, 0.8) | 7.198 | 5.411 | smooth spacetime | (4.0, 3.2) | 3.022 | 3.551 |
| piecewise abrupt | (10.0, 1.6) | 7.231 | 5.382 | smooth spacetime | (6.0, 0.1) | 3.963 | 3.473 |
| piecewise abrupt | (10.0, 3.2) | 7.328 | 5.342 | smooth spacetime | (6.0, 0.2) | 3.829 | 3.469 |

#### Stationary-subset ablation

Table 7 summarizes the stationary-subset ablation experiment described in Section 3. The table reports the performance when fitting the ST-RBF model using either the full track set or only the heuristically selected stationary subset. The column `n_seeds_found`from the original CSV has been omitted here.

**Table 7:** Stationary-subset ablation summary for the ST-RBF model.

| Regime | $(\ell_{\text{space}}, \ell_t)$ | Subset mode | Mean RMS [$\mu\text{m}$] | Drift bias [$\mu\text{m}$] | Std RMS [$\mu\text{m}$] |
| --- | --- | --- | --- | --- | --- |
| piecewise abrupt | (1.0, 1.6) | full | 7.694 | 5.156 | 0.778 |
| piecewise abrupt | (1.0, 1.6) | stationary | 14.412 | 11.927 | 8.521 |
| piecewise abrupt | (4.0, 1.6) | full | 7.216 | 5.267 | 0.756 |
| piecewise abrupt | (4.0, 1.6) | stationary | 7.497 | 5.632 | 0.698 |
| piecewise smooth | (1.0, 1.6) | full | 5.821 | 4.893 | 0.916 |
| piecewise smooth | (1.0, 1.6) | stationary | 7.345 | 6.945 | 2.714 |
| piecewise smooth | (4.0, 1.6) | full | 4.683 | 4.056 | 0.709 |
| piecewise smooth | (4.0, 1.6) | stationary | 4.698 | 4.229 | 0.638 |
| smooth spacetime | (1.0, 1.6) | full | 4.465 | 4.323 | 0.992 |
| smooth spacetime | (1.0, 1.6) | stationary | 5.364 | 5.018 | 1.181 |
| smooth spacetime | (4.0, 1.6) | full | 3.026 | 3.458 | 0.680 |
| smooth spacetime | (4.0, 1.6) | stationary | 3.016 | 3.571 | 0.643 |

#### Per-timepoint RBF robustness sweep

Table 8 reports the results of the robust-mode sweep for the per-timepoint RBF model shown in Figure 3b.

**Table 8:**
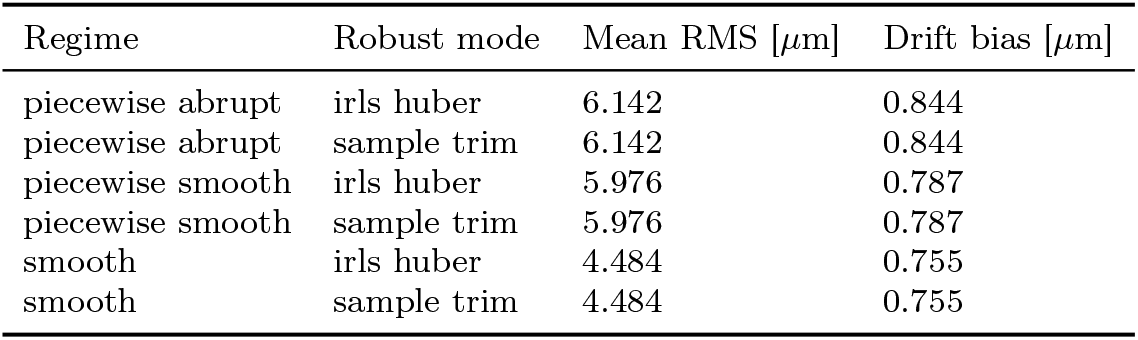
Robust-mode sweep for the per-timepoint RBF model.

| Regime | Robust mode | Mean RMS [$\mu\text{m}$] | Drift bias [$\mu\text{m}$] |
| --- | --- | --- | --- |
| piecewise abrupt | irls huber | 6.142 | 0.844 |
| piecewise abrupt | sample trim | 6.142 | 0.844 |
| piecewise smooth | irls huber | 5.976 | 0.787 |
| piecewise smooth | sample trim | 5.976 | 0.787 |
| smooth | irls huber | 4.484 | 0.755 |
| smooth | sample trim | 4.484 | 0.755 |

#### Space-time RBF robustness sweep

Table 9 reports the corresponding robust-mode sweep for the space-time RBF model (Figure 3c).

**Table 9:** Robust-mode sweep for the space-time RBF model.

| Regime | Robust mode | Mean RMS [$\mu\text{m}$] | Drift bias [$\mu\text{m}$] |
| --- | --- | --- | --- |
| piecewise abrupt | hybrid track irls | 4.819 | 0.777 |
| piecewise abrupt | irls huber | 4.882 | 0.814 |
| piecewise abrupt | sample trim | 4.850 | 0.798 |
| piecewise smooth | hybrid track irls | 4.666 | 0.735 |
| piecewise smooth | irls huber | 4.718 | 0.760 |
| piecewise smooth | sample trim | 4.685 | 0.749 |
| smooth | hybrid track irls | 2.750 | 0.848 |
| smooth | irls huber | 2.834 | 0.880 |
| smooth | sample trim | 2.780 | 0.840 |

#### Space-time RBF expressivity sweep

Table 10 reports the results of the expressivity sweep comparing isotropic versus anisotropic kernels and linear-term augmentation (Figure 3d).

**Table 10:** Expressivity sweep for the space-time RBF model.

| Regime | Model configuration | Mean RMS [$\mu\text{m}$] | Drift bias [$\mu\text{m}$] |
| --- | --- | --- | --- |
| piecewise abrupt | anisotropic off | 5.023 | 0.828 |
| piecewise abrupt | anisotropic on | 5.103 | 0.822 |
| piecewise abrupt | linear terms off | 5.023 | 0.828 |
| piecewise abrupt | linear terms on | 5.045 | 0.827 |
| piecewise smooth | anisotropic off | 4.835 | 0.760 |
| piecewise smooth | anisotropic on | 4.907 | 0.754 |
| piecewise smooth | linear terms off | 4.835 | 0.760 |
| piecewise smooth | linear terms on | 4.857 | 0.757 |
| smooth spacetime | anisotropic off | 3.078 | 0.684 |
| smooth spacetime | anisotropic on | 3.131 | 0.676 |
| smooth spacetime | linear terms off | 3.078 | 0.684 |
| smooth spacetime | linear terms on | 3.092 | 0.681 |

**Table 11:** Aggregated fragmentation statistics in synthetic point-cloud association baseline benchmarks. Shown values correspond to the heatmaps in Figure 5.

| Regime | Model configuration | Mean fragmentation rate | Std. dev. |
| --- | --- | --- | --- |
| 0.2 | hungarian_dummy_unmatched_cp | 0.9417 | 0.0191 |
| 0.2 | hungarian_gating_cp | 0.9417 | 0.0191 |
| 0.2 | hungarian_gating_cv | 1.0000 | 0.0000 |
| 0.2 | hungarian_huber_dummy_cp | 0.2750 | 0.0217 |
| 0.2 | hungarian_mahalanobis_diag_cp | 0.8833 | 0.0144 |
| 0.2 | kalman_cv_innovation_gating | 0.0000 | 0.0000 |
| 0.2 | min_cost_flow_cp | 0.9417 | 0.0191 |
| 0.2 | min_cost_flow_cv | 0.9417 | 0.0191 |
| 0.2 | nearest_gating_cp | 0.9417 | 0.0191 |
| 0.2 | tracklet_stitching_cv | 0.3000 | 0.0866 |
| 0.5 | hungarian_dummy_unmatched_cp | 0.6125 | 0.0375 |
| 0.5 | hungarian_gating_cp | 0.6125 | 0.0375 |
| 0.5 | hungarian_gating_cv | 0.4292 | 0.0505 |
| 0.5 | hungarian_huber_dummy_cp | 0.6125 | 0.0375 |
| 0.5 | hungarian_mahalanobis_diag_cp | 0.8583 | 0.0072 |
| 0.5 | kalman_cv_innovation_gating | 0.0083 | 0.0144 |
| 0.5 | min_cost_flow_cp | 0.6125 | 0.0375 |
| 0.5 | min_cost_flow_cv | 0.6125 | 0.0375 |
| 0.5 | nearest_gating_cp | 0.6125 | 0.0375 |
| 0.5 | tracklet_stitching_cv | 0.0125 | 0.0125 |
| 1.0 | hungarian_dummy_unmatched_cp | 0.4625 | 0.0433 |
| 1.0 | hungarian_gating_cp | 0.4625 | 0.0433 |
| 1.0 | hungarian_gating_cv | 0.3083 | 0.0711 |
| 1.0 | hungarian_huber_dummy_cp | 0.7958 | 0.0191 |
| 1.0 | hungarian_mahalanobis_diag_cp | 0.8833 | 0.0191 |
| 1.0 | kalman_cv_innovation_gating | 0.2583 | 0.0577 |
| 1.0 | min_cost_flow_cp | 0.7625 | 0.0217 |
| 1.0 | min_cost_flow_cv | 0.7625 | 0.0217 |
| 1.0 | nearest_gating_cp | 0.4625 | 0.0433 |
| 1.0 | tracklet_stitching_cv | 0.6208 | 0.0144 |
| 2.0 | hungarian_dummy_unmatched_cp | 0.3958 | 0.0382 |
| 2.0 | hungarian_gating_cp | 0.3958 | 0.0382 |
| 2.0 | hungarian_gating_cv | 0.2750 | 0.0545 |
| 2.0 | hungarian_huber_dummy_cp | 0.8375 | 0.0125 |
| 2.0 | hungarian_mahalanobis_diag_cp | 0.9000 | 0.0000 |
| 2.0 | kalman_cv_innovation_gating | 0.6375 | 0.0331 |
| 2.0 | min_cost_flow_cp | 0.9000 | 0.0000 |
| 2.0 | min_cost_flow_cv | 0.9000 | 0.0000 |
| 2.0 | nearest_gating_cp | 0.3958 | 0.0382 |
| 2.0 | tracklet_stitching_cv | 0.8917 | 0.0072 |
| 4.0 | hungarian_dummy_unmatched_cp | 0.3708 | 0.0382 |
| 4.0 | hungarian_gating_cp | 0.3708 | 0.0382 |
| 4.0 | hungarian_gating_cv | 0.2708 | 0.0577 |
| 4.0 | hungarian_huber_dummy_cp | 0.8542 | 0.0289 |
| 4.0 | hungarian_mahalanobis_diag_cp | 0.9000 | 0.0000 |
| 4.0 | kalman_cv_innovation_gating | 0.8167 | 0.0072 |
| 4.0 | min_cost_flow_cp | 0.9000 | 0.0000 |
| 4.0 | min_cost_flow_cv | 0.9000 | 0.0000 |
| 4.0 | nearest_gating_cp | 0.3958 | 0.0361 |
| 4.0 | tracklet_stitching_cv | 0.9000 | 0.0000 |

**Table 12:** Aggregated link F1 statistics in synthetic point-cloud association baseline benchmarks. Shown values correspond to the heatmaps in Figure 5.

| Regime | Model configuration | Mean link F1 | Std. dev. |
| --- | --- | --- | --- |
| 0.2 | hungarian_dummy_unmatched_cp | 0.8596 | 0.0131 |
| 0.2 | hungarian_gating_cp | 0.8596 | 0.0131 |
| 0.2 | hungarian_gating_cv | 0.7543 | 0.0122 |
| 0.2 | hungarian_huber_dummy_cp | 0.9907 | 0.0016 |
| 0.2 | hungarian_mahalanobis_diag_cp | 0.9113 | 0.0060 |
| 0.2 | kalman_cv_innovation_gating | 1.0000 | 0.0000 |
| 0.2 | min_cost_flow_cp | 0.8596 | 0.0131 |
| 0.2 | min_cost_flow_cv | 0.8443 | 0.0154 |
| 0.2 | nearest_gating_cp | 0.8596 | 0.0131 |
| 0.2 | tracklet_stitching_cv | 0.9911 | 0.0028 |
| 0.5 | hungarian_dummy_unmatched_cp | 0.9624 | 0.0093 |
| 0.5 | hungarian_gating_cp | 0.9624 | 0.0093 |
| 0.5 | hungarian_gating_cv | 0.9769 | 0.0054 |
| 0.5 | hungarian_huber_dummy_cp | 0.9624 | 0.0093 |
| 0.5 | hungarian_mahalanobis_diag_cp | 0.8309 | 0.0158 |
| 0.5 | kalman_cv_innovation_gating | 0.9998 | 0.0004 |
| 0.5 | min_cost_flow_cp | 0.9624 | 0.0093 |
| 0.5 | min_cost_flow_cv | 0.9624 | 0.0093 |
| 0.5 | nearest_gating_cp | 0.9624 | 0.0093 |
| 0.5 | tracklet_stitching_cv | 0.9997 | 0.0003 |
| 1.0 | hungarian_dummy_unmatched_cp | 0.9737 | 0.0080 |
| 1.0 | hungarian_gating_cp | 0.9737 | 0.0080 |
| 1.0 | hungarian_gating_cv | 0.9845 | 0.0054 |
| 1.0 | hungarian_huber_dummy_cp | 0.8891 | 0.0184 |
| 1.0 | hungarian_mahalanobis_diag_cp | 0.6653 | 0.0201 |
| 1.0 | kalman_cv_innovation_gating | 0.9874 | 0.0042 |
| 1.0 | min_cost_flow_cp | 0.9093 | 0.0138 |
| 1.0 | min_cost_flow_cv | 0.9093 | 0.0138 |
| 1.0 | nearest_gating_cp | 0.9737 | 0.0080 |
| 1.0 | tracklet_stitching_cv | 0.9426 | 0.0075 |
| 2.0 | hungarian_dummy_unmatched_cp | 0.9776 | 0.0056 |
| 2.0 | hungarian_gating_cp | 0.9776 | 0.0056 |
| 2.0 | hungarian_gating_cv | 0.9866 | 0.0032 |
| 2.0 | hungarian_huber_dummy_cp | 0.8171 | 0.0265 |
| 2.0 | hungarian_mahalanobis_diag_cp | 0.4860 | 0.0172 |
| 2.0 | kalman_cv_innovation_gating | 0.9304 | 0.0102 |
| 2.0 | min_cost_flow_cp | 0.5500 | 0.0137 |
| 2.0 | min_cost_flow_cv | 0.5500 | 0.0137 |
| 2.0 | nearest_gating_cp | 0.9776 | 0.0056 |
| 2.0 | tracklet_stitching_cv | 0.6066 | 0.0112 |
| 4.0 | hungarian_dummy_unmatched_cp | 0.9800 | 0.0035 |
| 4.0 | hungarian_gating_cp | 0.9800 | 0.0035 |
| 4.0 | hungarian_gating_cv | 0.9862 | 0.0033 |
| 4.0 | hungarian_huber_dummy_cp | 0.7844 | 0.0237 |
| 4.0 | hungarian_mahalanobis_diag_cp | 0.3881 | 0.0257 |
| 4.0 | kalman_cv_innovation_gating | 0.8047 | 0.0225 |
| 4.0 | min_cost_flow_cp | 0.3131 | 0.0206 |
| 4.0 | min_cost_flow_cv | 0.3097 | 0.0237 |
| 4.0 | nearest_gating_cp | 0.9780 | 0.0045 |
| 4.0 | tracklet_stitching_cv | 0.3293 | 0.0217 |

**Table 13:** Aggregated ID-switch statistics in synthetic point-cloud association baseline benchmarks. Shown values correspond to the heatmaps in Figure 5.

| Regime | Model configuration | Mean ID switches per track | Std. dev. |
| --- | --- | --- | --- |
| 0.2 | hungarian_dummy_unmatched_cp | 4.6083 | 0.3833 |
| 0.2 | hungarian_gating_cp | 4.6083 | 0.3833 |
| 0.2 | hungarian_gating_cv | 7.4542 | 0.3001 |
| 0.2 | hungarian_huber_dummy_cp | 0.3458 | 0.0591 |
| 0.2 | hungarian_mahalanobis_diag_cp | 3.0542 | 0.1954 |
| 0.2 | kalman_cv_innovation_gating | 0.0000 | 0.0000 |
| 0.2 | min_cost_flow_cp | 4.6083 | 0.3833 |
| 0.2 | min_cost_flow_cv | 4.8208 | 0.3934 |
| 0.2 | nearest_gating_cp | 4.6083 | 0.3833 |
| 0.2 | tracklet_stitching_cv | 0.3333 | 0.1048 |
| 0.5 | hungarian_dummy_unmatched_cp | 1.3292 | 0.3098 |
| 0.5 | hungarian_gating_cp | 1.3292 | 0.3098 |
| 0.5 | hungarian_gating_cv | 0.8167 | 0.1821 |
| 0.5 | hungarian_huber_dummy_cp | 1.3292 | 0.3098 |
| 0.5 | hungarian_mahalanobis_diag_cp | 5.3042 | 0.4438 |
| 0.5 | kalman_cv_innovation_gating | 0.0083 | 0.0144 |
| 0.5 | min_cost_flow_cp | 1.3292 | 0.3098 |
| 0.5 | min_cost_flow_cv | 1.3292 | 0.3098 |
| 0.5 | nearest_gating_cp | 1.3292 | 0.3098 |
| 0.5 | tracklet_stitching_cv | 0.0125 | 0.0125 |
| 1.0 | hungarian_dummy_unmatched_cp | 0.9333 | 0.2717 |
| 1.0 | hungarian_gating_cp | 0.9333 | 0.2717 |
| 1.0 | hungarian_gating_cv | 0.5417 | 0.1954 |
| 1.0 | hungarian_huber_dummy_cp | 3.6083 | 0.5635 |
| 1.0 | hungarian_mahalanobis_diag_cp | 9.3250 | 0.4564 |
| 1.0 | kalman_cv_innovation_gating | 0.4333 | 0.1371 |
| 1.0 | min_cost_flow_cp | 2.9958 | 0.4390 |
| 1.0 | min_cost_flow_cv | 2.9958 | 0.4390 |
| 1.0 | nearest_gating_cp | 0.9333 | 0.2717 |
| 1.0 | tracklet_stitching_cv | 1.8917 | 0.2526 |
| 2.0 | hungarian_dummy_unmatched_cp | 0.7875 | 0.1887 |
| 2.0 | hungarian_gating_cp | 0.7875 | 0.1887 |
| 2.0 | hungarian_gating_cv | 0.4667 | 0.1188 |
| 2.0 | hungarian_huber_dummy_cp | 5.6042 | 0.7140 |
| 2.0 | hungarian_mahalanobis_diag_cp | 12.7833 | 0.2765 |
| 2.0 | kalman_cv_innovation_gating | 2.1708 | 0.3260 |
| 2.0 | min_cost_flow_cp | 11.6375 | 0.2385 |
| 2.0 | min_cost_flow_cv | 11.6375 | 0.2385 |
| 2.0 | nearest_gating_cp | 0.7875 | 0.1887 |
| 2.0 | tracklet_stitching_cv | 10.5375 | 0.2315 |
| 4.0 | hungarian_dummy_unmatched_cp | 0.6875 | 0.1068 |
| 4.0 | hungarian_gating_cp | 0.6875 | 0.1068 |
| 4.0 | hungarian_gating_cv | 0.4500 | 0.0901 |
| 4.0 | hungarian_huber_dummy_cp | 6.4292 | 0.6101 |
| 4.0 | hungarian_mahalanobis_diag_cp | 14.3417 | 0.3767 |
| 4.0 | kalman_cv_innovation_gating | 5.7417 | 0.6362 |
| 4.0 | min_cost_flow_cp | 15.4333 | 0.2810 |
| 4.0 | min_cost_flow_cv | 15.4708 | 0.3146 |
| 4.0 | nearest_gating_cp | 0.7292 | 0.1301 |
| 4.0 | tracklet_stitching_cv | 15.2042 | 0.3001 |

#### RBF-family robustness analyses and hyperparameter sweeps across baseline stress regimes

## Supplementary Information synthetic point-cloud association benchmarks

## Supplementary Methods for biological example workflows

### Biological datasets

We evaluated the workflow on two publicly available biological time-lapse datasets chosen to represent complementary regimes of geometric complexity and image formation.

The first example was taken from the dataset *Test datasets for segmentation (2D, 3D, timelapses)* by Kapoor (Kapoor, 2022). From this collection, we used a low-SNR carcinoma-cell volume sequence acquired as a 3D+*t* fluorescence stack. This dataset was originally released as a segmentation stress test for denoising and instance-segmentation models under adverse imaging conditions. In the present work, it serves as a controlled volumetric benchmark in which the biological objects are sparse enough to permit construction of a segmentation-derived trajectory reference, while the image quality remains sufficiently challenging to expose errors in segmentation, temporal linking, and downstream identity reconstruction.

The second example was taken from the dataset *Paired bright-field and DAPI time-lapse microscopy dataset of fibroblast-like cells (synoviocytes) from osteoarthritic knee tissue* by Radvansky *et al*. (Radvansky et al., 2026). We used the DAPI fluorescence sequence corresponding to a single field of view of primary human fibroblast-like synoviocytes derived from osteoarthritic synovial tissue. In contrast to the carcinoma example, these data form a 2D+*t* time series without axial sampling and exhibit substantial experimental drift, bleaching-related intensity variation, dense cell populations, and long observation duration. This dataset was therefore used as a realistic planar stress case in which drift, imperfect detection continuity, and trajectory fragmentation arise intrinsically from the acquisition rather than from an imposed synthetic perturbation.

Taken together, these two examples probe complementary aspects of the pipeline: the carcinoma sequence provides a volumetric benchmark with known imposed drift and an explicit geometric reference, whereas the synoviocyte DAPI sequence provides a planar benchmark with native experimental drift and substantially greater temporal extent and population density.

### Three-dimensional carcinoma benchmark with synthetic rigid drift

#### Volume segmentation and construction of a proxy track reference

The carcinoma example consists of a 3D+*t* fluorescence stack with *T* = 12 time points and volumetric observations on a Cartesian grid. For each time point, segmentation was performed independently in three spatial dimensions. Let *I*_*t*_(*z, y, x*) denote the observed intensity volume at time *t*. To stabilize contrast across frames, each volume was first intensity-normalized by clipping to the empirical 5th and 99.8th percentiles and rescaling to [0, 1]. A mildly smoothed representation was then obtained by anisotropic Gaussian filtering. From this representation, a primary object mask was constructed by a hysteresis-type thresholding procedure: a high threshold was estimated by the Otsu criterion on the smoothed intensities, seed voxels were defined by exceedance of this threshold, and these seeds were propagated into a support region defined by a lower threshold proportional to the high threshold. Morphological closing, hole filling, and component-size filtering were then applied.

To recover dimmer objects that were only weakly represented in the primary branch, a second rescue branch was applied after coarse background subtraction by subtracting a broader Gaussian background estimate from the normalized volume. The enhanced image was thresholded again, followed by the same propagation step, morphological regularization, and component filtering. Rescue components were accepted only if both their size and their peak response exceeded prescribed minimum values. The final segmentation at time *t* was defined as the union of the primary and rescue masks, followed by a final round of binary regularization and connected-component labeling.

These per-frame components were then linked across time to obtain a proxy reference trajectory table. Linking proceeded in two stages. First, each component at time *t* + 1 was assigned, whenever possible, to the component label at time *t* with maximal voxel overlap. Second, components without overlap-based continuation were matched by nearest centroid continuation subject to a maximal admissible displacement. Unassigned components initiated new trajectories. In addition, components that gave rise to trajectories present in only a single frame were removed by a post hoc clean-up step before downstream analysis, as such one-frame detections were judged biologically implausible in this example and are more consistent with segmentation artefacts than with transient true cells.

The resulting tracked centroids constitute a *proxy ground truth* rather than a manually curated truth set. Its failure modes are therefore inherited from both segmentation and temporal linking. These include missed low-contrast cells, spurious detections, temporary mergers of adjacent cells, over-segmentation of elongated objects, and erroneous identity changes induced by overlap ambiguity or large centroid displacements between consecutive frames.

#### Synthetic rigid drift and definition of input tracks

The raw carcinoma stack exhibited no sufficiently strong intrinsic geometric drift to serve as a stress test for point-cloud correction. We therefore imposed a known synthetic drift field on the proxy reference trajectories. Only the rigid-drift regime was retained for the analyses discussed in the manuscript. Let *x*_*i*_(*t*) ∈ ℝ^3^ denote the centroid position of reference trajectory *i* at time *t*.

The observed input coordinates were generated by

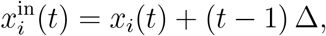

where Δ ∈ ℝ^3^ is a constant translation step applied uniformly to all trajectories. Thus the imposed drift is globally rigid, temporally cumulative, and identical for all objects. This construction provides a controlled stress regime in which the latent trajectories remain fixed while the observations are corrupted by a known acquisition-like bias. In the figures, these rigidly drifted reference trajectories were denoted *input GT*, whereas the undrifted trajectories *x*_*i*_(*t*) were retained as the true positional reference.

#### Blind point-cloud input for association

For point-cloud association, the drifted input trajectories were converted into a blind observation table by retaining only frame index and spatial coordinates and suppressing track identity. Thus, at each time point *t*, the association methods received an unlabeled point set

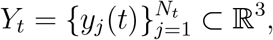

with no explicit information about which points corresponded to the same underlying object across time. The known correspondence between each blind point and the latent input-trajectory identity was stored separately and used only for evaluation.

#### Association and drift-correction pipeline

All six HCV-based association variants were applied to the blind point clouds. For each method, the association parameters were selected from prior synthetic sweeps, and an optional short-gap stitching step was enabled at the track-linking stage to permit continuation across a prescribed number of missing intermediate frames without altering the underlying pairwise assignment model. The resulting inferred trajectories were scored against the proxy reference by link precision, link recall, link *F*_1_, fragmentation, identity-switch rates, and GT-consistent accuracy. The best-performing association method for the present example was then used as input to the drift-correction stage.

The drift-correction methods did not operate on the blind point clouds directly. Instead, they received the best PCA/HCV-inferred trajectory table as input and estimated correction fields from those inferred trajectories. Consequently, the full PCA → DriftCorr workflow is sequential: association first, geometric correction second.

#### One-to-one remapping for GT-based evaluation

A subtle but important consequence of the association step is that the inferred trajectories need not be in one-to-one correspondence with the proxy reference trajectories. In particular, a single reference trajectory may be fragmented into several inferred segments, while in other cases multiple reference trajectories may be merged. To obtain an unambiguous GT-based evaluation of drift correction, we therefore constructed a deterministic one-to-one remapping between inferred and reference identities. Let *c*_*gh*_ denote the number of matched observations supporting an association between reference track *g* and inferred track *h*. Candidate pairs (*g, h*) were sorted in descending order of *c*_*gh*_, and a greedy one-to-one matching was constructed by accepting each pair in turn only if neither *g* nor *h* had been used previously. This procedure yields a disjoint set of matched trajectory pairs.

This remapping was used only for GT-based visualization and metric calculation after association. It was *not* used to fit the drift-correction models themselves, which operated on the full inferred trajectories. Accordingly, if association fragmented a biologically continuous reference track into multiple inferred segments, only one of those segments could survive in the one-to-one GT remapping used for downstream GT-referenced drift-correction panels and RMS-type metrics. This explains why certain inferred subsegments may be visible in the PCA output but absent from GT-referenced DriftCorr overlays.

#### Interpretation of drift-correction errors

Because the true positional reference remains the undrifted proxy trajectory set {*x*_*i*_(*t*)}, GT-based drift-correction RMS errors in this example quantify the performance of the *entire* PCA → DriftCorr pipeline. They therefore conflate two error sources: association errors such as fragmentation, merging, or identity switches, and subsequent errors in the estimated drift field. A low RMS indicates that the combined pipeline reconstructs trajectories close to the undrifted reference. Conversely, an elevated RMS does not imply that the drift-correction model itself is poor in isolation; it may instead arise because the drift estimator was supplied with fragmented or otherwise imperfect trajectories.

### Two-dimensional DAPI benchmark with intrinsic drift

#### Motivation and overall design

The DAPI example differs fundamentally from the carcinoma example in two respects. First, the imaging data are 2D+*t* rather than 3D+*t*. Second, the stack already exhibits substantial intrinsic drift and frame-to-frame intensity variation, so no additional synthetic drift was imposed. This example was therefore used to test the practical behavior of the full PCA→DriftCorr workflow under realistic experimental drift, rather than under a controlled synthetic perturbation. Additionally, this example benchmarked the 2D-capabilities of both the association and drift-correction methods, since the existing mathematical core of these methods is formally three-dimensional but can be applied to planar data by embedding the coordinates into ℝ^3^ with a dummy axial coordinate.

#### Framewise preprocessing and nucleus segmentation

Let *I*_*t*_(*y, x*) denote the fluorescence image at time *t*. To compensate for bleaching and slow temporal intensity changes, each frame was histogram-matched to the first frame before segmentation. Nuclei were then segmented independently in each frame using Cellpose (Stringer et al., 2021) in its nuclear-segmentation mode with a characteristic object radius calibrated empirically for this dataset. Small components and implausibly large components were removed by area filtering. Because the downstream association and drift-correction methods are formulated for three-dimensional coordinates, each extracted centroid (*x*_*i*_(*t*), *y*_*i*_(*t*)) was embedded into ℝ^3^ as

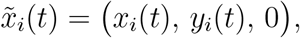

that is, with a dummy axial coordinate fixed to zero.

#### Post hoc temporal linking of segmented nuclei

The Cellpose segmentations were linked across time by two post hoc methods. The baseline method constructed a frame-to-frame overlap graph and took connected components of this graph as trajectories. The main reference method used a Hungarian assignment linker whose cost combined centroid displacement, overlap of segmented masks between consecutive frames, and relative area change. A short-gap option was allowed, so that a trajectory could remain active across a limited number of missing intermediate frames. An additional hard spatial jump constraint was imposed to suppress implausible long-range reassignments. Among the two temporal linkers, the Hungarian-linked trajectories were adopted as the reference trajectory set for the downstream benchmark because they yielded substantially fewer implausible fragments and more visually coherent tracks than the pure overlap-based baseline.

As in the carcinoma example, this linked trajectory table is not a manual cell-tracking truth set. It is a segmentation-derived proxy reference and therefore remains susceptible to false splits, false merges, missed detections, identity switches near close encounters, and artefactual track fragmentation when nuclei are temporarily absent from the segmentation.

#### Blind point-cloud association in the planar setting

The Hungarian-linked DAPI trajectories were converted into blind point clouds by suppressing identity labels and retaining only time and coordinates. Since all axial coordinates were set to zero, the association problem remained formally three-dimensional but was geometrically confined to the plane *z* = 0. This allowed the existing point-cloud association and drift-correction methods to be applied without modifying their mathematical core.

As in the carcinoma example, all six HCV-based association variants were evaluated on these blind observations. A short-gap stitching option was enabled in the association pipeline to reduce artifactual fragmentation caused by brief segmentation dropouts. The best-performing association result according to mean link *F*_1_ was then passed to the drift-correction stage.

#### Why Cellpose was used in the DAPI example

The DAPI dataset contains many small, densely packed nuclei over a long time series with pronounced bleaching and substantial drift. In this setting, a purely threshold-based segmentation produced unacceptable large-scale foreground structures and unstable temporal behavior. A learned nucleus model was therefore preferred because it provides stronger shape priors and is empirically more stable under local intensity variation. By contrast, the carcinoma dataset contains comparatively sparse three-dimensional cellular objects for which a carefully regularized volumetric thresholding and connected-component workflow was sufficient and had the practical advantage of remaining fully deterministic and transparent.

#### Why no additional synthetic drift was applied in the DAPI example

Unlike the carcinoma stack, the DAPI sequence already contains strong experimental drift. Superimposing an additional synthetic transformation would therefore have confounded two distinct sources of motion and would have made interpretation of the corrected trajectories less transparent. The DAPI example was instead treated as a natural stress case in which the observed linked tracks already incorporate unknown geometric bias from the experiment itself.

### Downstream inputs to PCA and DriftCorr

#### Association input

In both biological examples, point-cloud association was applied to blind point sets derived from linked trajectory tables with identities removed. Thus association was tasked with recovering track continuity from unordered per-frame detections alone.

#### Drift-correction input

In both examples, the main DriftCorr analyses were applied not to the proxy reference trajectories, but to the best-performing PCA/HCV-inferred trajectories. Consequently, the drift-correction methods operated on whatever trajectory fragmentation, merging, or identity-switch structure had already been introduced by the association stage. In auxiliary control analyses of the carcinoma example, the same drift-correction methods were also applied directly to identity-labeled input trajectories to isolate the geometric component of the correction problem.

#### GT-referenced versus input-referenced views

In the carcinoma example, the rigidly drifted trajectories serve as the observed input trajectories, whereas the undrifted trajectories remain the true positional reference. In the DAPI example, the Hungarian-linked Cellpose tracks serve as a practical reference for association benchmarking, but no external drift-free positional truth is available. Accordingly, the carcinoma example supports GT-referenced RMS-type geometric evaluation, whereas the DAPI example primarily supports reference-based association evaluation and qualitative or descriptive drift-correction summaries.

### Quantitative summaries and motion-class analyses

#### Association metrics

Association quality was summarized by link precision, link recall, and link *F*_1_, together with fragmentation rates, identity-switch rates, and a GT-consistent accuracy measure. These metrics were computed from the blind-to-reference correspondence table and the inferred link structure, thereby quantifying both local link correctness and trajectory-level identity stability.

#### Track-count and birth–death summaries

For both examples, we additionally summarized the total number of unique trajectories observed across the entire sequence and the number of births and deaths after the initial frame. Here, a birth at time *t* denotes the appearance of a trajectory identity not present at time *t* − 1, and a death denotes the disappearance of a previously present identity.

#### Stationary versus migrating trajectories

To separate weakly fluctuating trajectories from genuinely migrating ones, we computed frame-to-frame step lengths

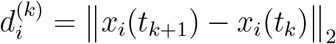

for each trajectory *i*. In the carcinoma example, motion class was based on the per-track median step length,

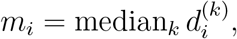

and the stationary–migrating threshold was estimated from the reference distribution of {*m*_*i*_} by fitting a two-component Gaussian mixture to log(1 + *m*_*i*_) and taking the intersection point of the two component densities. In the DAPI example, motion class was based on the per-track mean step length,

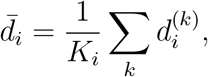

and the threshold was estimated by applying the Otsu criterion to 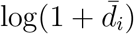 over the reference trajectory set. Trajectories with summary displacement below threshold were labeled stationary, and those above threshold were labeled migrating.

#### Implications and limitations

All reference-dependent metrics in these biological examples must be interpreted with the caveat that the references are segmentation-derived and linker-derived rather than manually curated cell-tracking truths. Thus disagreement between PCA or DriftCorr outputs and the reference can arise either because the method failed or because the constructed reference itself contains merges, splits, missed detections, or identity errors. In the carcinoma example, GT-referenced DriftCorr RMS values therefore quantify end-to-end performance of the combined association-plus-correction pipeline. In the DAPI example, where no drift-free ground truth is available, the downstream analyses quantify relative changes in trajectory consistency and motion statistics rather than absolute geometric recovery error.

